# CCM signaling complex (CSC) is a master regulator governing homeostasis of progestins and their mediated signaling cascades

**DOI:** 10.1101/2020.06.10.145003

**Authors:** Johnathan Abou-Fadel, Xiaoting Jiang, Akhil Padarti, Dinesh Goswami, Mark Smith, Brian Grajeda, Wendy Walker, Jun Zhang

**Author notes:** All correspondence: Jun Zhang, Sc.D., Ph.D. Department of Molecular and Translational Medicine (MTM) Texas Tech University Health Science Center El Paso 5001 El Paso Drive, El Paso, El Paso, TX 79905. TTUS has filed a provisional patent, 63/034,787, regarding certain aspects of this publication.

## Abstract

We demonstrate that a novel signaling network among the CSC and mPRS is dynamically modulated and fine-tuned with intricate feedback regulations in PR negative cells, especially endothelial cells (ECs). Depletion of any of three CCMs (1, 2, 3) genes results in the disruption of non-classic mPRs-mediated signaling *in-vitro* as well as defective homeostasis of PRG *in-vivo*. Therefore, we propose the CSC is a master regulator of homeostasis of PRG and its associated classic and non-classic signaling cascades. Assisted with omic approaches, we identified signaling pathways involved and specific biomarkers associated with hemorrhagic events during CCM pathogenesis *in-vitro* and *in-vivo*. To our knowledge, this is the first report detailing etiology to predict the occurrence of early hemorrhagic events with a set of serum biomarkers.

---

Cerebral cavernous malformations (CCMs) are characterized by abnormally dilated intracranial microvascular sinusoids that result in increased susceptibility to hemorrhagic stroke. At least one of three genes, KRIT1 (CCM1), MGC4607 (CCM2) and PDCD10 (CCM3), is disrupted in most human CCM cases. It has been demonstrated that both CCM1 and CCM3 bind to CCM2 to form a CCM signaling complex (CSC). Recent identification of multiple CCM2 isoforms with distinctive functions (*1*) and extensive efforts on the comparative omics of the CSC (*2, 3*) make the ongoing investigation of CSC signaling cascades even more challenging and exciting. Despite all three CCM proteins being expressed ubiquitously among tissues, differential expression of all three CCM genes were recently discovered in various tissues and cells at both the transcriptional and translational levels (*1–5*). Progesterone (PRG) binds to its nuclear receptor (PR) in classic genomic actions (*6*). PR has two major isoforms, PR1/2 (*7*), and interestingly vascular expression of PR1 isoform is greater in women than in men, suggesting the gender role of PRG in vascular angiogenesis (*8*). PRG can also evoke rapid membrane and cytoplasm changes through ion channel activation, defined as rapid, non-classical or non-genomic actions (*9, 10*). Currently, most studies of PRG have been focused on genomic and non-genomic actions of the classical PR1/2. Recently, two new groups of non-classical PRG receptors (*11*) have been identified as the membrane progesterone receptors (mPRs) (*12, 13*), and the progesterone receptor membrane components (PGRMCs)(*14, 15*). Unlike classic PR, mPRs function as membrane receptors usually coupling to G proteins for their signaling (*16, 17*).

It has been demonstrated that three CCMs (CCM1, CCM2, and CCM3) form a CCM signaling complex (CSC) to mediate angiogenic signaling (*1, 3, 18*). In this report, we elucidate the effects of a disrupted CSC on PRG-mPRs-mediated signaling cascades in PR negative (-) cells both *in-vitro* and *in-vivo*, maintenance and performance of microvascular ECs modulated by PRG-mPRs-mediated signaling, and the role of differentially expressed CCM proteins in the biogenesis of PRG at both the transcriptional and translational levels. We also identified serum biomarkers to predict the occurrence of hemorrhage initiation, which could have enormous clinical application in stroke prevention.

## Results

### Distinct PRG receptor signaling under sex steroid actions in PR(-) cells

#### Unique responses to non-classic progesterone signaling under steroid actions in 293T cells expressing only trace PR1/2

To further elucidate the cellular relationship between PR1/2 and mPRs/PAQRs upon steroid actions, we investigated the relative expression level of total PR1/2 and mPRs/PAQRs under combined steroid treatment (PRG+MIF) on 293T cells which has only traceable levels of PR1/2 (Suppl. Fig. 1A). We initially hypothesized that under the depletion of PR1/2, drastic changes could be observed under steroid actions due to this unbalanced CSC-mPRs signaling cascade. Surprisingly, no change of PR1/2 or mPRs/PAQRs expression was found either at the transcriptional (Figs. 1A) or translational (Figs. 1B) levels, under PRG+MIF treatment in 293T cells. The lack of changes in PR1/2 RNA under PRG+MIF treatment can possibly be explained by its traceable expression, however, mPRs/PAQRs responses are quite different from the data observed in PR(+) T47D cells, recently reported (*19*). Using RNAi, we knocked down CCM1, CCM2, and CCM3 respectively in 293T cells and examined RNA expression levels of total PR1/2 and PAQRs with RT-qPCR analysis. The results showed that changed RNA expression patterns of PAQRs were only observed by silencing CCM2, with increased RNA expression levels of PAQR 6, 7 and 9 (Fig. 1C), similar to the data from PR(+) T47D cells (*19*). The phenomena that significantly changed RNA expression patterns of mPRs/PAQRs by silencing CCM2, suggests the reciprocal regulation between mPRs/PAQRs and the CSC is mediated through CCM2. Increased protein expression levels of most mPRs/PAQRs were observed, especially under CCM1-KD condition which increased PAQR7/8. CCM3-KD resulted in increased expression of PAQR7, but decreased levels of PAQR5/8. CCM2-KD only resulted in significant decreased expression levels of PAQR5/8. These results do not follow the same trend observed in T47D cells (other than decreased PAQR8 expression in CCM2-KD), indicating a different regulation mechanism of the CSC on mPRs/PAQRs expression at the translational level (Fig. 1D). Next, inhibitory roles of PRG/MIF in both RNA and protein expression levels of CCMs (1, 2, 3) genes in 293T cells was examined. RT-PCR data demonstrated that PRG+MIF treatment can only inhibit RNA expression of CCM2 isoforms but had no effect on the RNA expression of either *CCM1 or CCM3* genes (Fig. 1E) which is identical to the data in T47D cells (*19*). These overlaps suggest the phenomena that *CCM2* is the direct target for sex steroid actions at the transcriptional level, and the potential central role of CCM2 in these steroid actions on the CSC, applies to both PR(+) and PR(-) cells. The observation in PR (+) T47D cells, that both progesterone and mifepristone can independently inhibit protein expression levels of CCM1 and CCM3, as well as the combination of both working synergistically to enhance their inhibitory effects (*19*) were also confirmed in 293T cells (Fig. 1F), reaffirming a common mechanism in regulation of inhibitory roles of PRG/MIF in the CSC, regardless of PR(+)/PR(-) cell types. Transient transfection experiments showed that 293T cells expressing CCM2 isoforms with CCM3 binding site (CCM2-100, CCM2-200) can stabilize CCM3 protein levels under PRG+MIF treatments compared to a CCM2 isoform lacking a CCM3 binding site (CCM2-1209, and expression vector) (Fig. 1G), validating the cornerstone role of CCM2 in the CSC under steroid actions. Our data illustrated that silencing major mPRs without steroid actions has little effect on the protein levels of CCM1 and CCM3 (Suppl Fig. 2A), suggesting the CSC may play a major role in this relationship under steroid actions. Finally, increased expression levels of CCM1/3 proteins were observed by silencing PAQR5/7/8 in combination with MIF+PRG treatment (Fig. 1H), identical to the observations seen in T47D cells (*19*), indicating a common mechanism in the feedback regulatory relationship of mPRs/PAQRs to the CSC under steroid actions, regardless of PR(+)/PR(-) cell types. Several mPRs-null mutant fish strains were identified without any visual phenotype (*20*), strongly indicating that mPR genes share redundancy (*21*), which is further supported by our data in this section. Overall, these data also reaffirm previous data that CCM2 isoforms might be the key factors influencing the expression levels of all mPRs/PAQRs at both the transcriptional and post-translational levels, while CCM1/3 proteins might be the key factors to influence the expression levels of mPRs/PAQRs at the post-translational level. Based on our current data, we define 293T cells as “Type-1A” PR(-) cells in response to sex hormone actions.

**Fig. 1.**
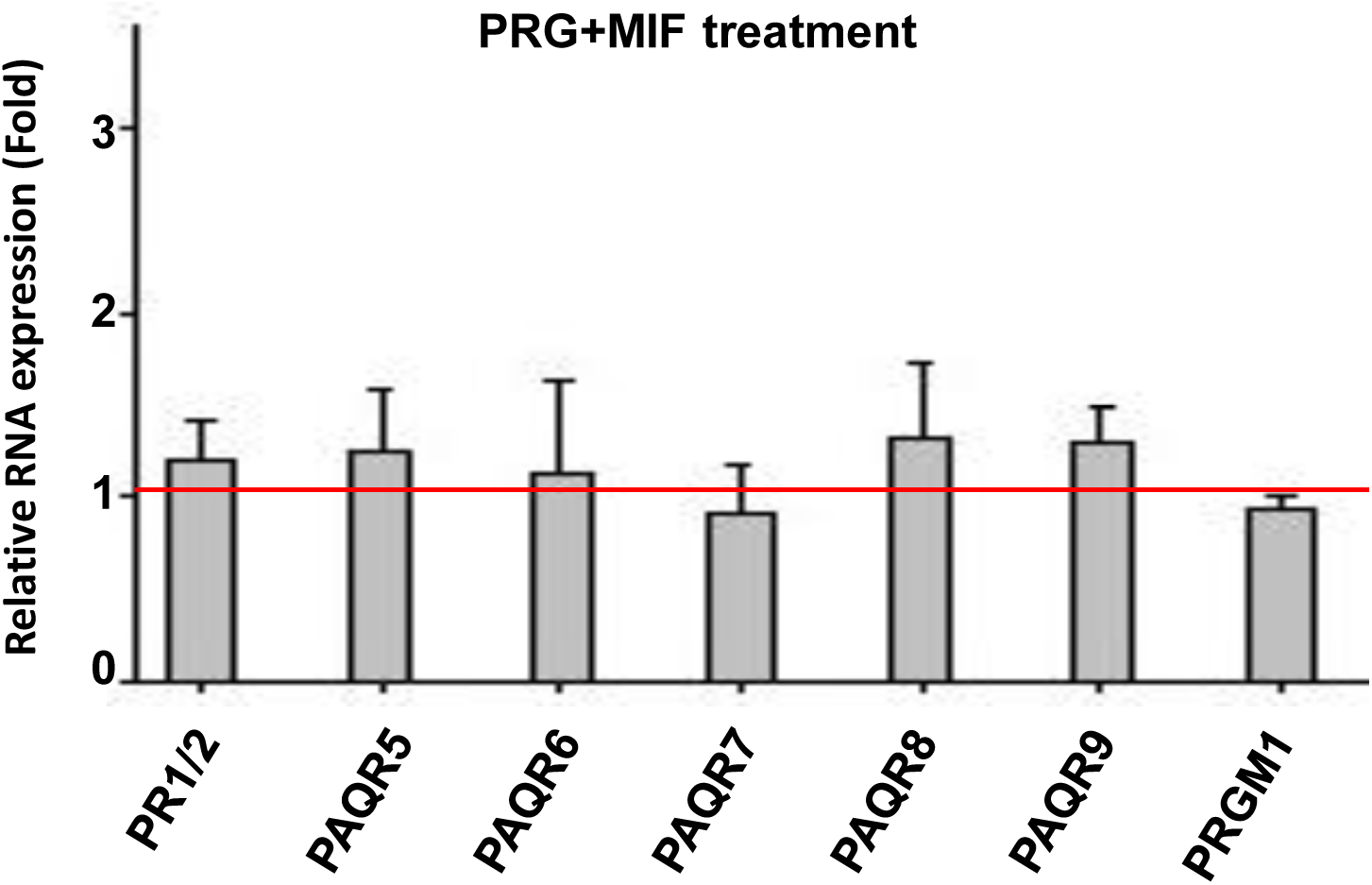
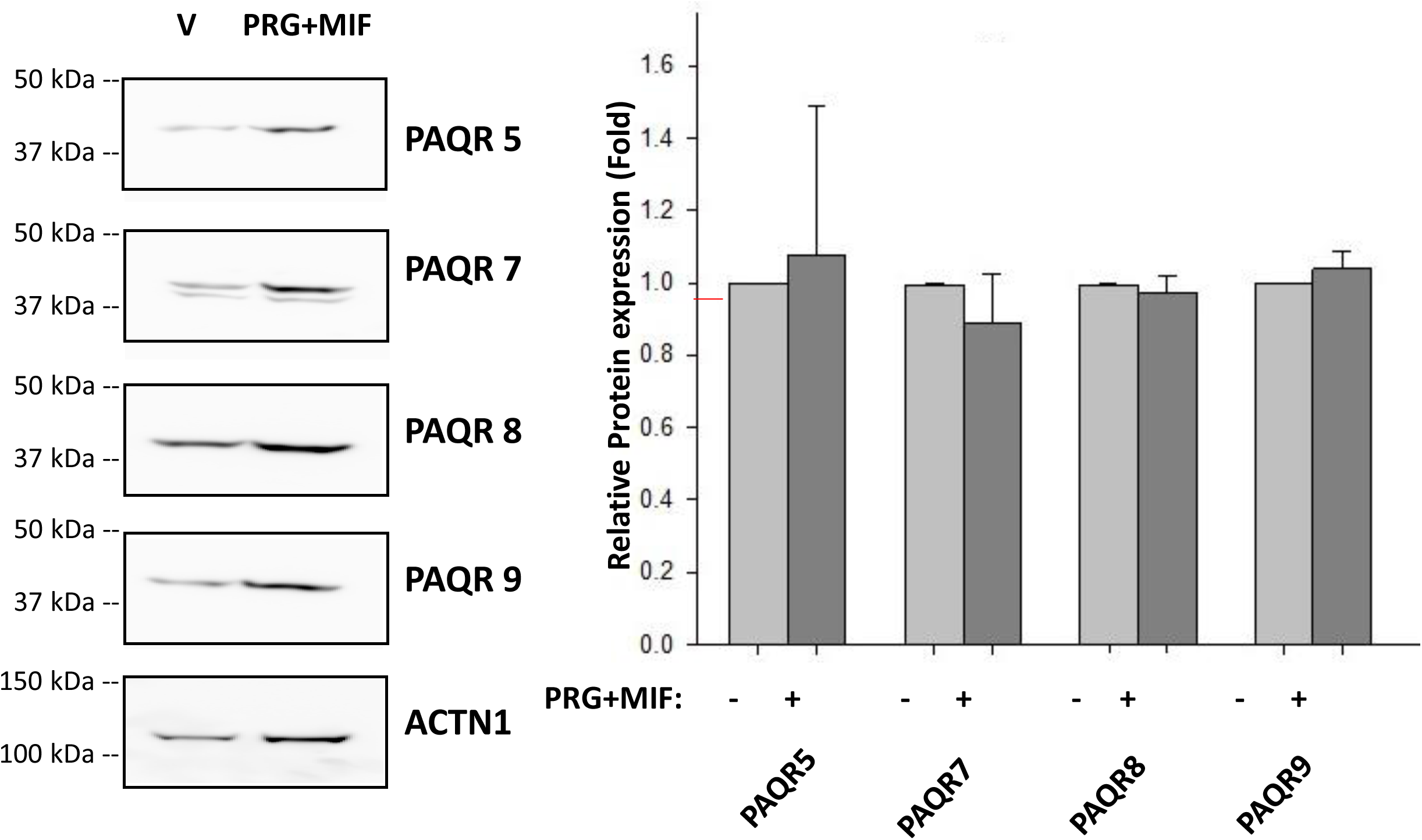
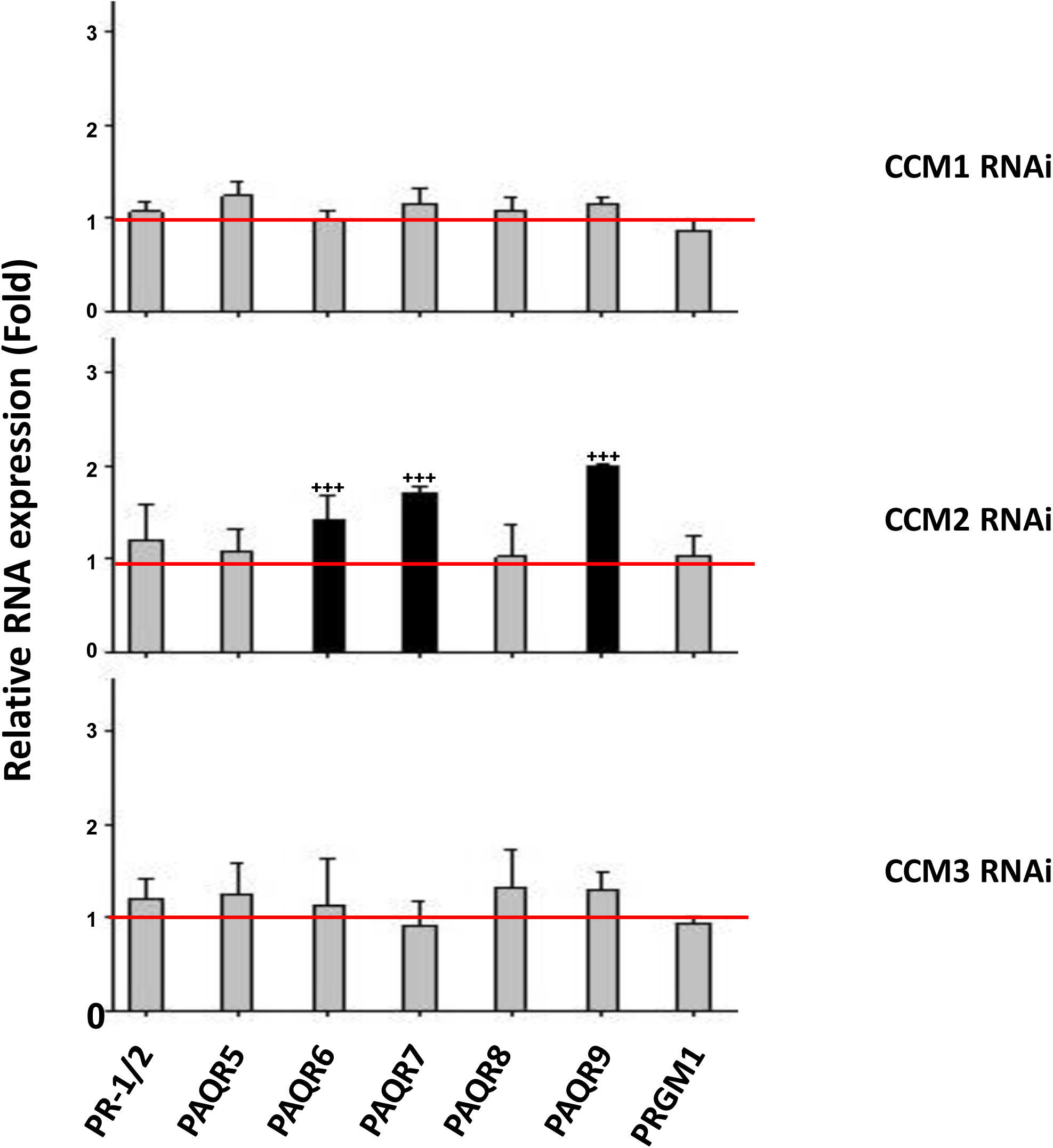
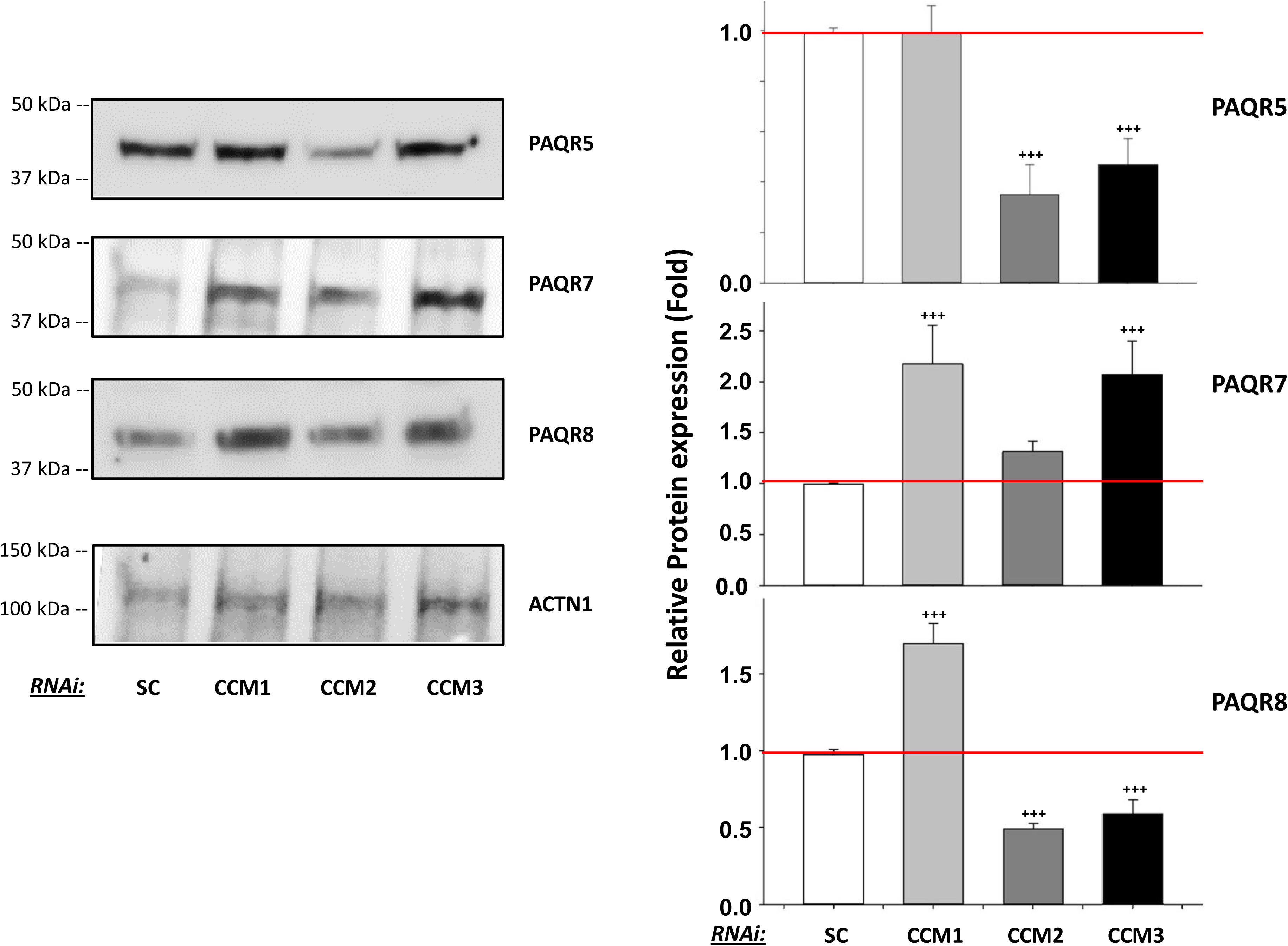
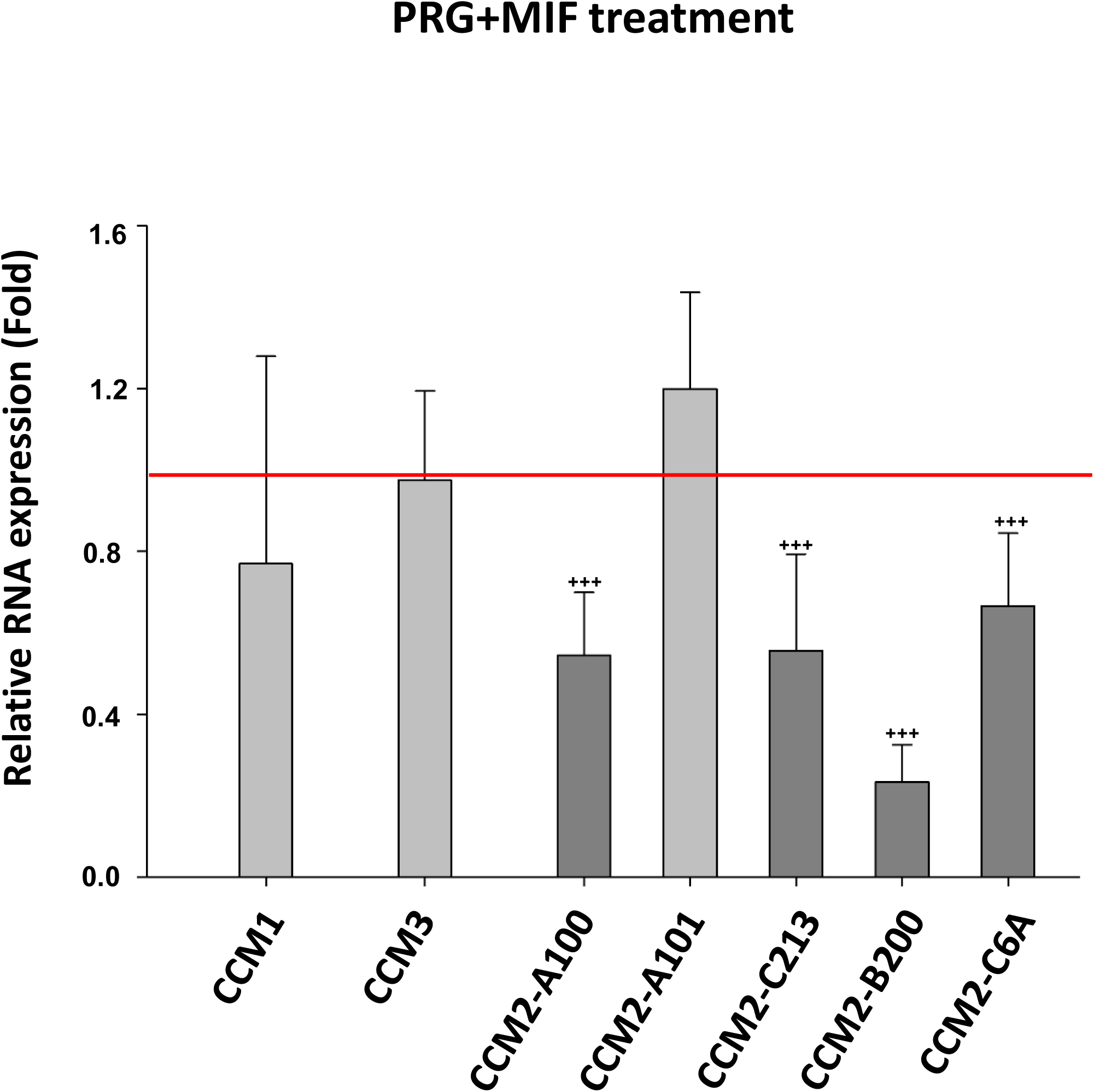
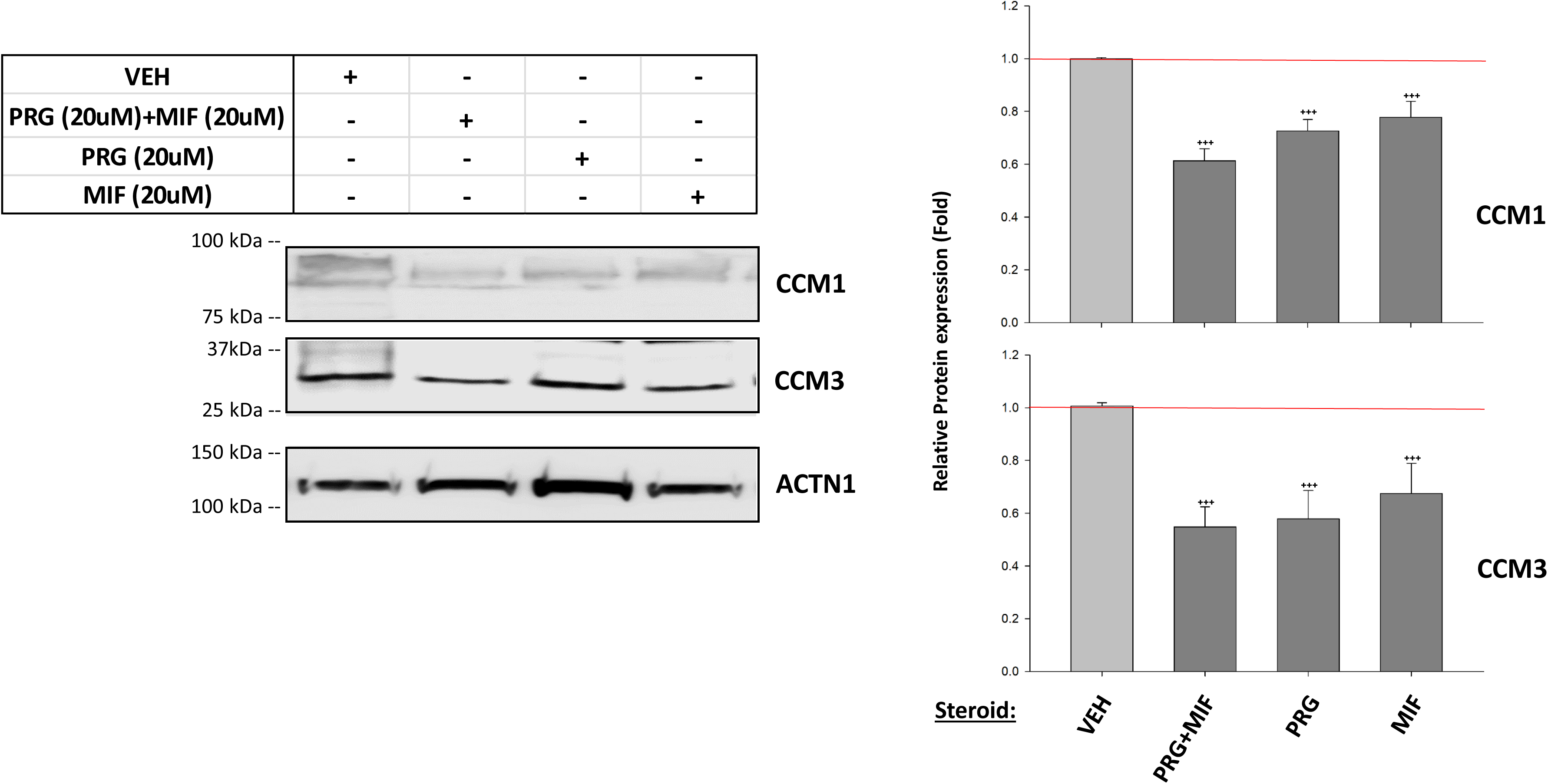
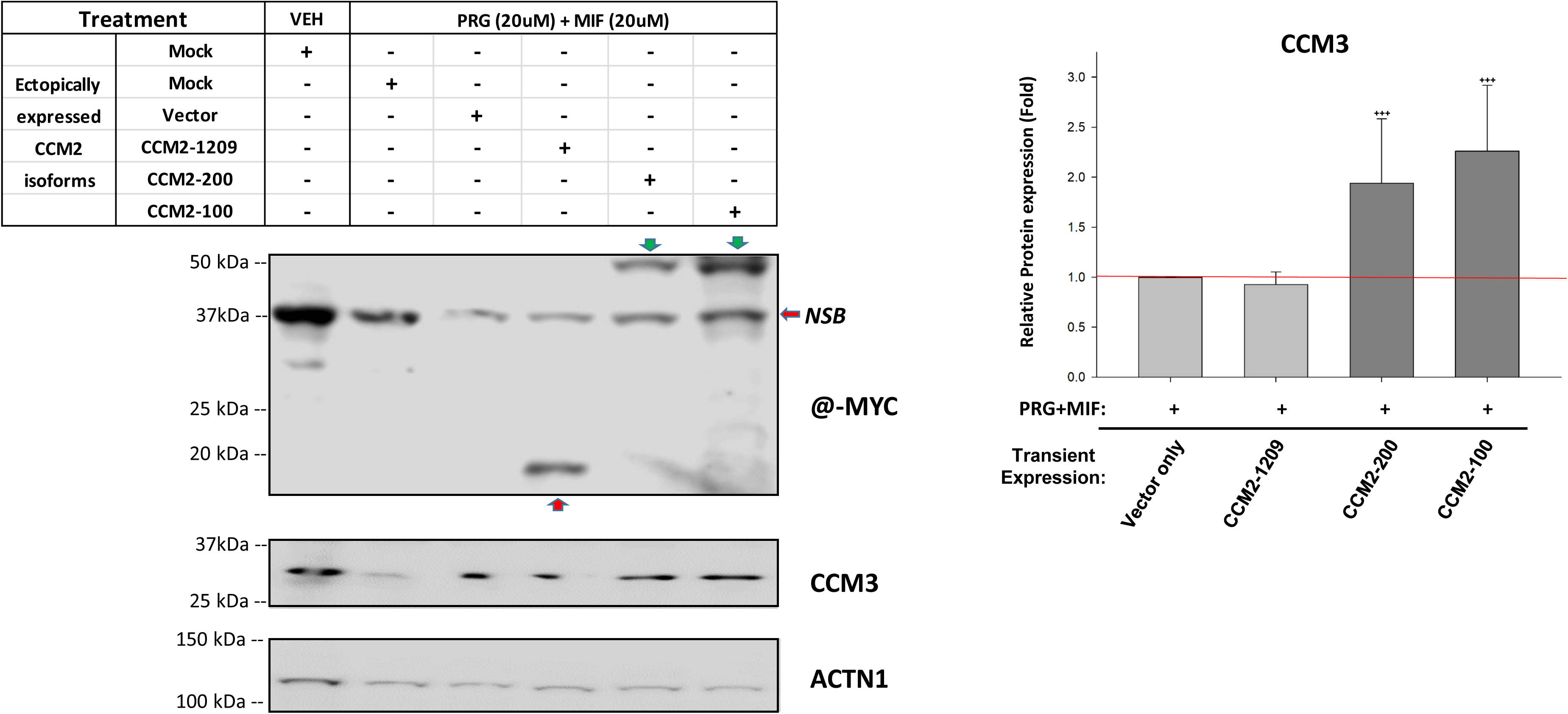
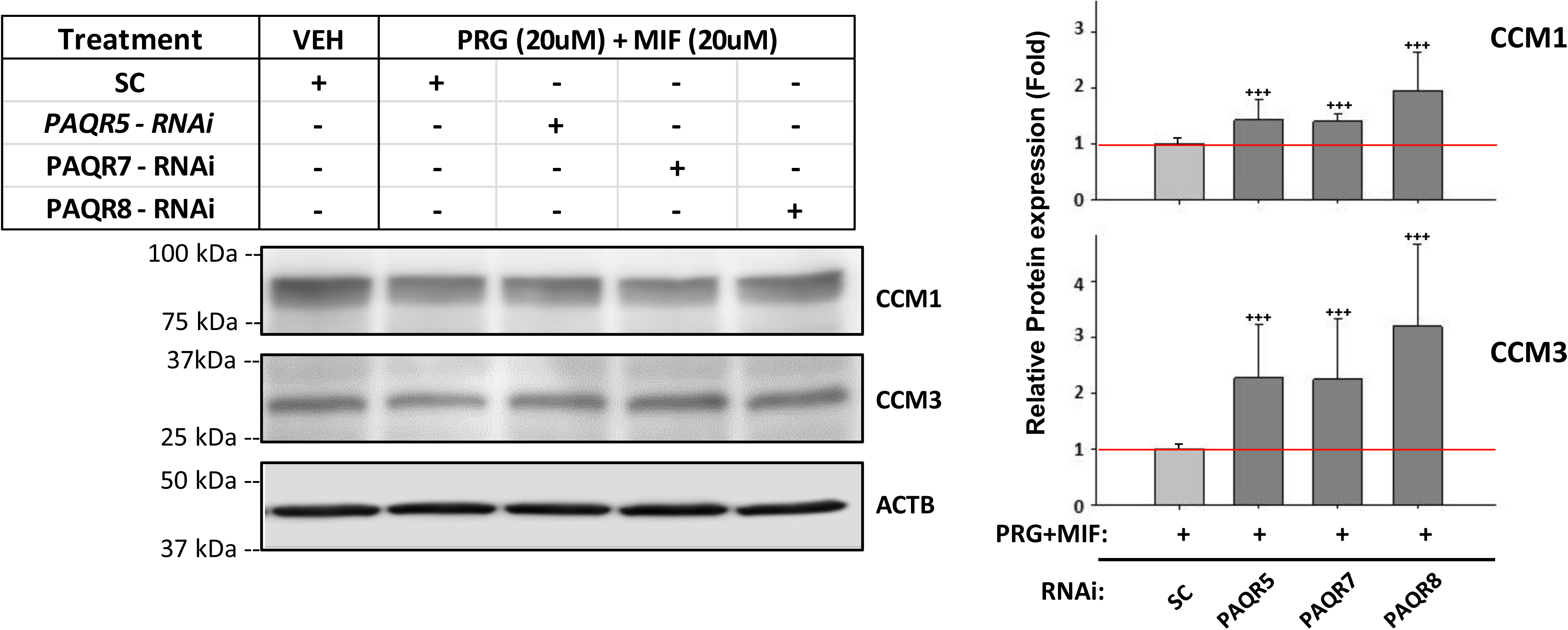
The relationships among PR1/2, PAQRs, PRG/MIF, and the CSC in a PR-traceable cell line, 293T. **A.** RNA expression levels of PR1/2 and mPRs/PAQRs are not influenced by PRG+MIF in 293T cells. Although PR1/2 RNAs are barely detected through RT-PCR in 293T cells (Suppl. Fig. 1), the relative expression changes of PR1/2 and mPRs/PAQRs were examined by RT-qPCR (Fold) in 293T cells after combined steroid treatment (PRG+MIF, 20 µM each) or vehicle control for 48 hrs. No apparent RNA expression change of either PR1/2 or mPRs/PAQRs was observed (n=3). **B.** Protein expression levels of PAQR 5, 7, 8 and 9 are not influenced by PRG+MIF treatment. After PRG+MIF treatment (20 µM each) for 48 hrs, no change of relative expression levels of PAQR 5, 7, 8 or 9 were observed, (n=3). **C**. RNA expression levels of PAQRs are only influenced by CCM2 gene expression in 293T cells. After silencing all three *CCMs* (1, 2, 3) genes for 48 hrs, increased RNA expression levels of PAQR6, 7, and 9 was only observed under CCM2-deficiency while no changed expression levels of mPRs were found under either CCM1 or CCM3-KD conditions. Further, no RNA expression changes of PR1/2 or PGRMC1 was found in 293T cells. The relative RNA expression changes of PR1/2/PAQRs/PGRMC1 were examined by RT-qPCR (Fold) in 293T cells (n=3). **D.** Protein expression levels of PAQR 5, 7 and 8 are modulated by the CSC in 293T cells. After silencing all three *CCMs* (1, 2, 3) genes for 48 hrs, increased PAQR7 expression was observed under CCM1/3-deficiency while significant decreased PAQR5 and PAQR8 protein levels were observed when silencing CCM2/3. PAQR8 expression increased when silencing CCM1, (n=3). **E**. PRG+MIF actions target RNA expression of CCM2 in 293T cells. Only decreased RNA expression of CCM2 was observed under PRG+MIF (20 µM each) for 48 hrs, while expression levels of CCM1/3 genes were not influenced in 293T cells. The relative RNA expression changes of CCM1/3, and 5 isoforms of CCM2 were measured by RT-qPCR (Fold) (n=3). **F.** PRG/MIF work synergistically to enhance their inhibitory roles in protein expression of CCM proteins in 293T cells. Decreased protein expression levels of CCM1/3 were observed under the treatment of progesterone (20 µM), mifepristone (20 µM), and PRG+MIF (20 µM each) for 48 hrs. PRG/MIF can independently inhibit protein expression levels of CCM1/3 and this effect is further amplified in PRG+MIF, demonstrating a synergistic effect to enhance suppression on protein expression levels of CCM1/3 (Left Panel), (n=3). **G.** Three CCM2 isoforms (CCM2-100/200/1209, tagged with MYC) along with expression vector alone were transiently transfected into 293T cells for 24 hrs, followed by PRG+MIF treatment (20 µM each), or vehicle control for an additional 48 hrs. Significant increased expression of CCM3 protein was observed only in ectopically expressed CCM2 isoforms containing aPTB domain (CCM2-100, CCM2-200) (left middle panel). Anti-Myc antibody was used to detect expressed CCM2 isoforms with MYC-tag, (n=3). **H.** Both CCM1 and CCM3 proteins are stabilized to PRG+MIF treatment after silencing mPRs in 293T cells. After silencing PAQR5, 7 or 8 for 24 hrs, followed by PRG+MIF (20 µM each) treatment for additional 48 hrs, increased levels of CCM1/3 were observed in mPRs-KD 293T cells, compared to SC controls (Left panel), (n=4). More details provided in Suppl. Fig. 2C.

#### Angiogenic responses to sex steroid actions through non-classic progesterone signaling in PR(-) vascular endothelial cells (ECs)

Vascular ECs are a group of PR(-) and mPRs-expressing cells (Suppl. Figs. 1B-C) and rapid, nongenomic actions of PRG on ECs have been previously reported (*22*). Therefore, several microvascular EC cells, including human brain microvascular endothelial cells (HBMVEC), human dermal microvascular endothelial cells (HDMVEC), human umbilical vein endothelial cells (HUVEC), and rat brain microvascular endothelial cells (RBMVEC) cells, were utilized to investigate angiogenic signaling modulated by the CSC and mPRs/PAQRs upon combined steroid actions. Interestingly, low protein expression of mPRs/PAQRs were found in these ECs, and no change of PAQR7 protein expression was found among HBMVEC, HDMVEC, HUVEC, and RBMVEC cells under MIF+PRG treatment (Suppl. Fig. 2B), identical to the trends observed in 293T cells (Fig. 1B). The RNA expression pattern of both mPRs/PAQRs and CCMs genes were examined upon steroid actions, and similar increased RNA expression patterns of both CCMs genes (Fig. 2A) and mPRs/PAQRs (Fig. 2B) were observed among HBMVEC, HDMVEC, and HUVEC cells. Likewise, silencing all three *CCMs* (*1, 2, 3*) genes lead to significantly decreased protein expression of PAQR7 in both HBMVEC and HDMVEC cells (Fig. 2C). Finally, the expression levels of CCM1 and CCM3 proteins were examined upon MIF+PRG treatment among HBMVEC, HDMVEC, HUVEC, and RBMVEC cells, reinforcing that PRG works with its antagonist, MIF, synergistically to inhibit the protein expression of CCM1/3 through mPRs (Fig. 2D), as seen previously in PR(-) 293T cells (Fig. 1F) and PR(+) T47D cells (*19*). However, sex hormone inhibition on protein expression of CCM1/3 in EC cells (Fig. 2D) is more dramatic than what was observed in either PR(-) 293T cells (Fig. 1F) or PR(+) T47D cells (*19*), suggesting that steroid hormones have stronger actions on the stability of the CSC through mPRs in ECs. To understand the molecular mechanisms of this CSC-mPRs coupled signaling cascade, the feedback network among the CSC/mPRs under steroid actions in type 1 PR(-) cells was schematically summarized, where key feedback regulatory pathways at both the transcriptional and translational levels are illustrated with detailed supporting data from corresponding experiments. In Type 1A 293T cells, it can be seen that PRG/MIF has an inhibitory effect at both the transcriptional and translational levels for the CSC, but can be stabilized from PRG+MIF actions by transiently expressing CCM2 isoforms (Fig. 2E, left panel). Furthermore, it appears that mPRs have a negative effect on the CSC at the proteomic level, however the CSC also exhibit signs of dual roles in feedback regulation of certain mPRs expression at the same level. Interestingly, the CSC is capable of modulating mPRs expression at the transcriptional level, extending our findings of intricate feedback loops that modulate expression at each level for this CSC-mPRs signaling cascade. In comparison to steroid response profiling of 293T cells (type-1A), we have defined PR(-) ECs as type-1B in response to sex hormone actions. From this schematic diagram, it is clearly shown that the transcriptional levels of both *CCM/mPRs* genes have been enhanced under feedback-regulation to compensate for the destabilization of CCMs/mPRs proteins under steroid actions at the post-translational level (Fig. 2E, right panel).

**Fig. 2.**
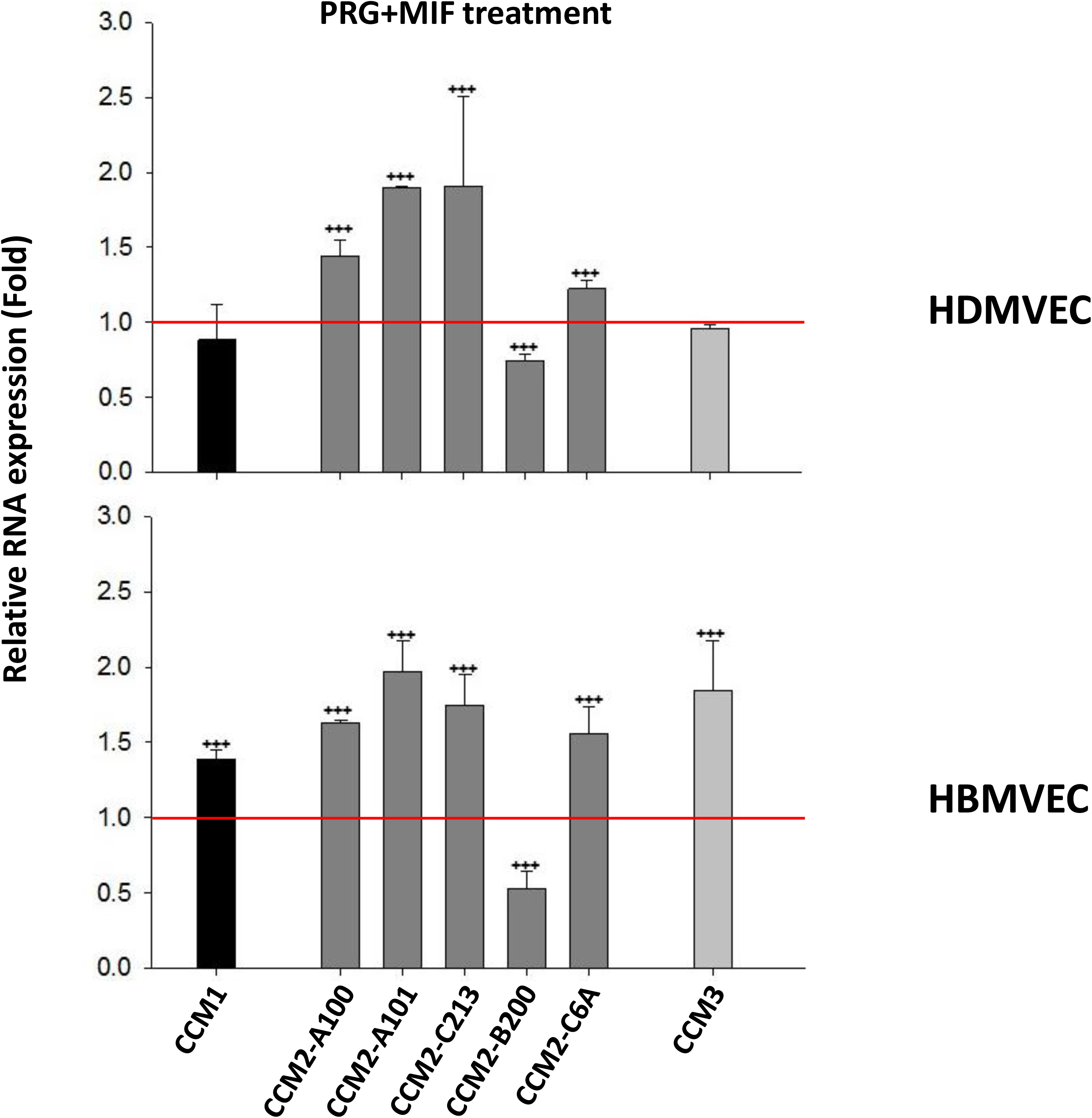
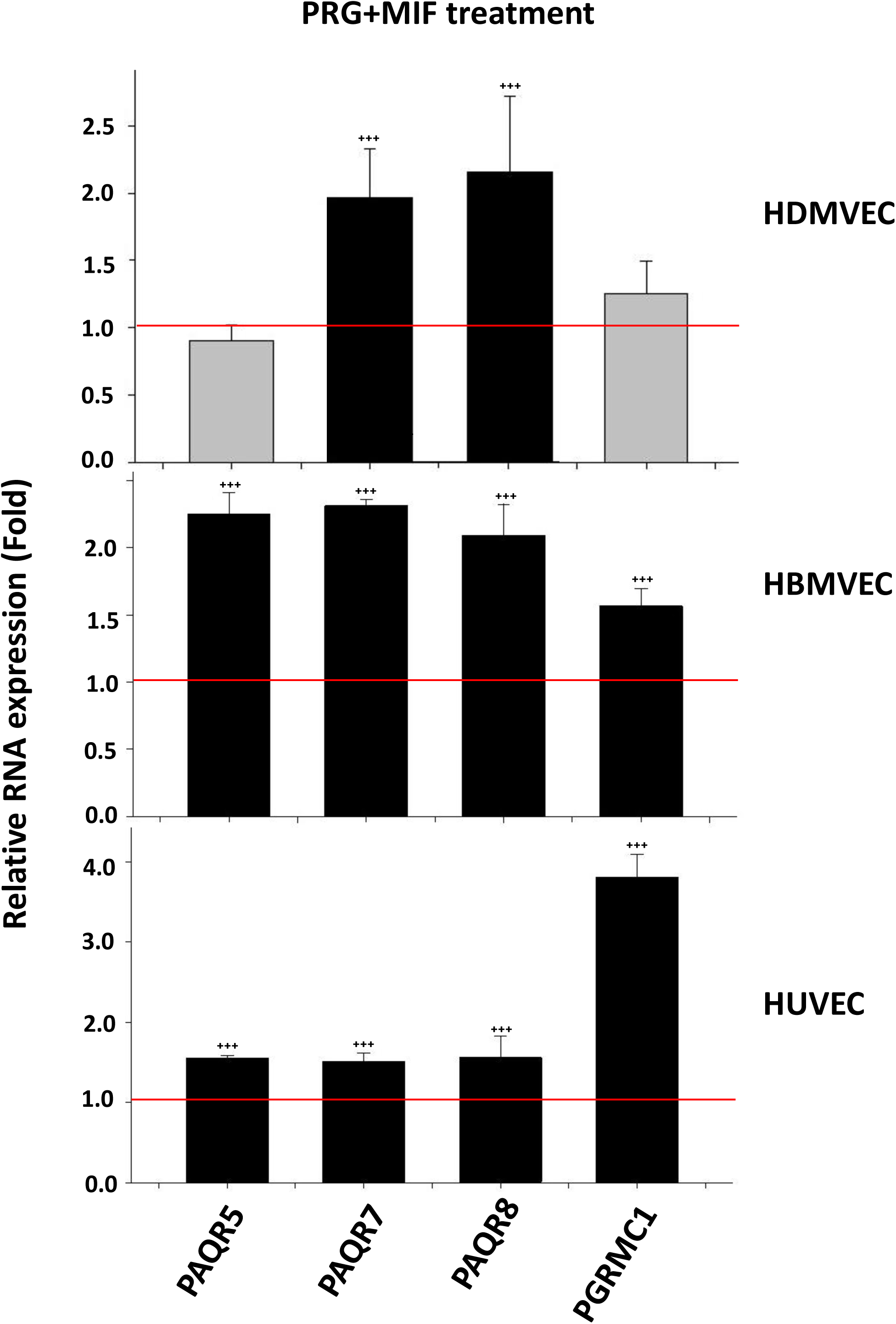
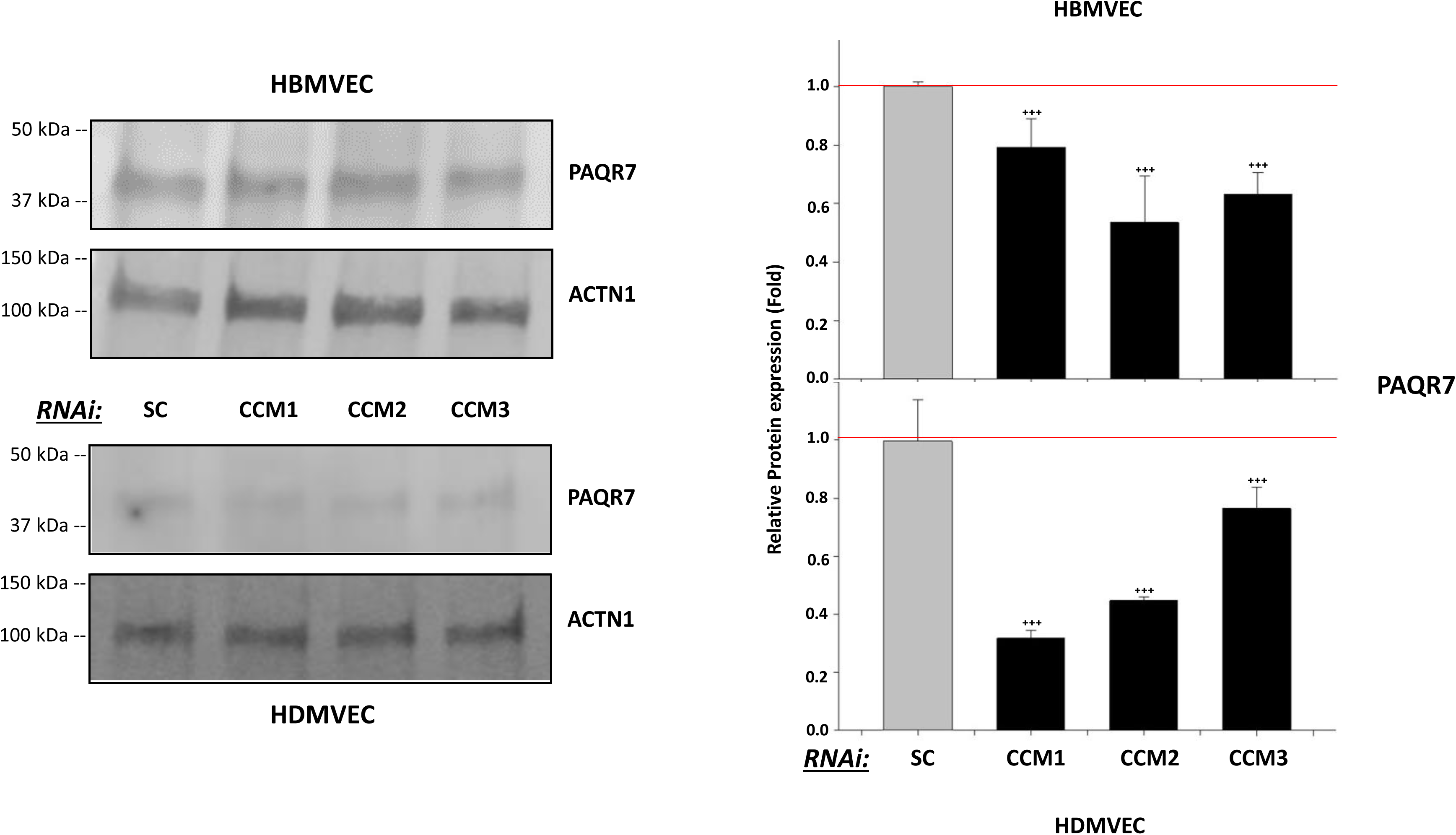
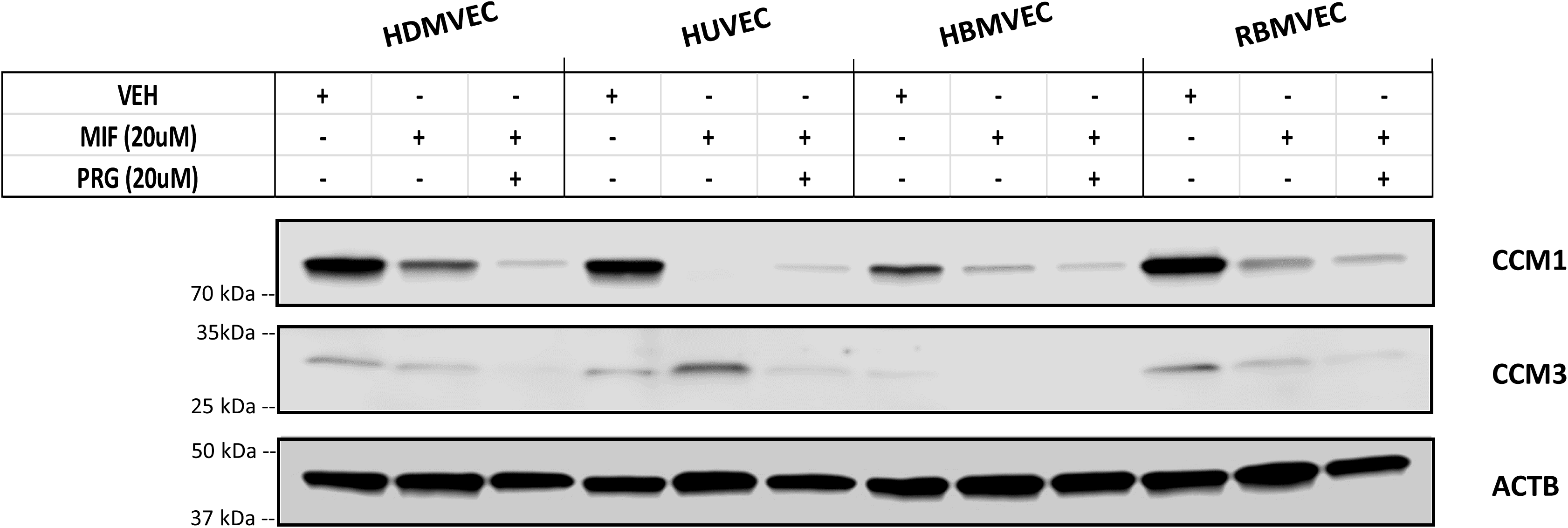
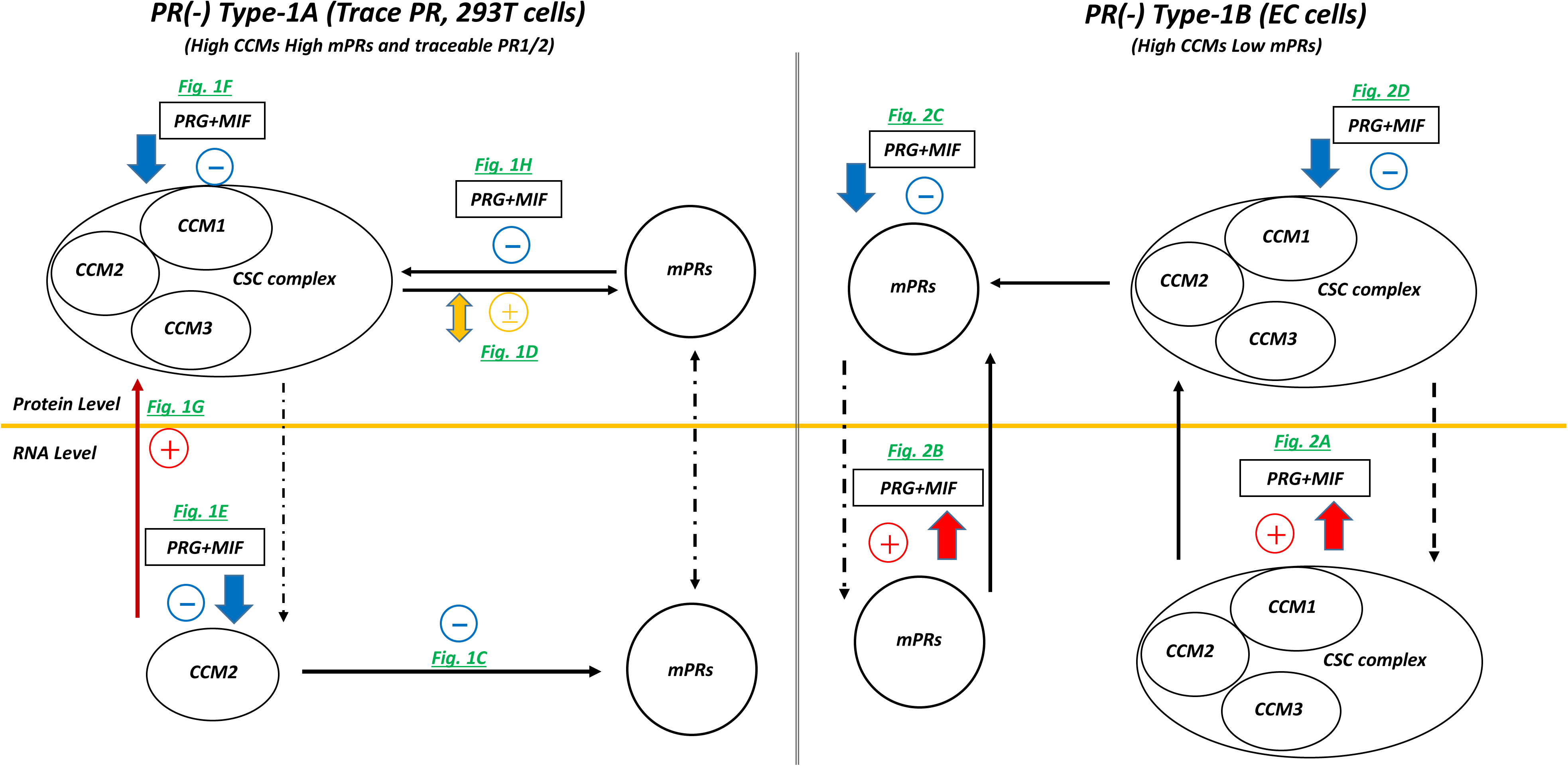
Relationships among PAQRs, its ligand(s) PRG/MIF, and the CSC complex in PR(-) endothelial cell (EC) lines. **A**. Under steroid treatment (PRG+MIF) for 48 hrs, enhanced RNA expression levels of most *CCM2* isoforms was observed in both Human Dermal Microvascular Endothelial cells (HDMVEC) and Human Brain Microvascular Endothelial cells (HBMVEC) cells, while increased RNA expression level of *CCM1*/*3* were only observed in HBMVEC cells. The relative RNA expression level of three *CCMs* (CCM1, CCM2, CCM3) genes were measured through RT-qPCR (Fold changes) (n=3). **B**. Significantly increased RNA expression of PAQR5, 7, 8 and PGRMC1 genes were observed under PRG+MIF treatment in HDMVEC, HBMVEC and Human Umbilical Vascular Endothelial cells (HUVEC) for 48 hrs, suggesting RNA expression of most mPRs can be dramatically enhanced by steroid treatment. The relative RNA expression levels of PAQRs/PGRMC1 genes were measured through RT-qPCR (Fold changes) (n=3). **C**. Protein expression levels of PAQR7 are influenced by all 3 Ccms gene expression in ECs. After silencing all three *CCMs* (1, 2, 3) genes for 48 hrs, decreased protein expression levels of PAQR7 was observed for all 3 Ccms-KD conditions. The relative protein expression changes of PAQR7 were examined by western blots in ECs (n=3). **D**. PRG and MIF can work synergistically to enhance their inhibitory roles in protein expression of CCM proteins in PR(-) endothelial cell (EC) lines. The combination of PRG+MIF (20 µM each) synergistically enhance their inhibitory effects on protein expression levels of CCM1/3 in HDMVEC, HUVEC, HBMVEC, and Rat Brain Microvascular Endothelial cells (RBMVEC), compared to mifepristone only (MIF, 20 µM) or vehicle controls (VEH). **E**. The summarized feedback regulatory networks among the CSC, PR1/2, and mPRs signaling complex under combined steroid treatment for PR(-) cell lines. Type 1A is 293T cells, while Type 1B is PR(-) EC cell lines. Yellow line separates transcriptional and translational levels. The + symbols represent enhancement, - symbols represent inhibition while ± symbol represent various regulation for the expression of targeted genes/proteins. Red colored symbols/lines represent positive effects of combined steroid treatment (PRG+MIF), blue colored symbols/lines represent negative effects of treatment, while orange colored symbols/lines represent variable effects. Dark green colored letters indicate the direct supporting data generated from this experiment. Arrow indicates effect direction, solid line is the direct impact, dotted line for indirect effects. More details provided in Suppl. Fig. 2C.

#### Regulation of signaling cascades among microvascular PR(-) ECs under steroid actions at the transcriptional levels

To further delineate the molecular mechanisms of steroid actions in ECs, we utilized high throughput RNAseq analysis to profile differentially expressed genes (DEGs) in HDMVECS, HBMVECs and RBMVECs after PRG+MIF treatment (Suppl. Table 3A). Identified DEGs were also compared to our previously published omic data generated from *CCMs* (*1, 2, 3*) genes knockdown (KD) experiments in HBMVECs (*2, 3*). As expected, initial profiling of DEGs demonstrated more similarities (in terms of DEGs) between HBMVECs and RBMVECs compared to HDMVEC (Fig. 3A, panels i-iii). When DEGs were evaluated using hierarchical clustering software, the data demonstrated vast differences between control and steroid treated samples among all 3 cell lines, demonstrating the profound impact combined steroid actions has on gene expression, regardless of cell type. Interestingly, it appears that most alterations at the transcriptional level for HDMVECs occur through down-regulation of genes (as seen by blue color in the clustering scheme) compared to HBMVEC and RBMVEC (Fig. 3A, panels iv-vi). Shared disrupted signaling pathways among all three cell lines, when compared to controls, included FOXO, P53, Oocyte meiosis, and progesterone mediated oocyte maturation (Fig. 3A, panels vii-ix). Pathways unique to HDMVEC included HIF-1, RAS, ovarian steroidogenesis and steroid biosynthesis signaling pathways (Fig 3A, panel vii). VEGF signaling, interestingly, was only affected in HBMVECs (Fig. 3A, viii), while disruption of regulation of actin/cytoskeleton, focal adhesion and ErbB signaling pathways was only observed in RBMVECs (Fig. 3A, ix), demonstrating the wide range of effects steroid actions can have on different EC sub-types. Furthermore, a wider view of KEGG pathways enrichment demonstrated a wide range of effects on altering multiple sub-processes including cardiovascular disease, neurodegenerative disease, multiple cancer signaling pathways, development, circulatory system and even aging signaling pathways that were shared among all three cell lines under steroid actions (Fig. 3A, panels x-xii). To distinguish binding partners in the synergistic action with PRG+MIF among the three EC cell lines, we compared DEG expression patterns of HBMVEC, HDMVEC and RBMVEC cells through RNAseq; we were able to visualize hierarchical clustering, and found similar patterns in both intersection and union of DEGs between all three cell lines (Figs. 3Bi-iii and Suppl. Table 3B), suggesting shared signaling cascade variations between them. We then evaluated these overlapped clusters to identify the specific overlapped DEGs, to compare side by side among all three EC cell lines (Fig. 3C and Suppl. Table 3C). Interestingly, the only shared DEGs among all three cell lines were all down-regulated under steroid actions, further confirming shared signaling cascades among all three cell lines. Functional enrichment performed from these overlapped DEGs revealed shared alterations among several key pathways including cell cycle/cell division, cytoskeletal protein binding, oocyte meiosis, progesterone mediated oocyte maturation and p53 signaling pathways (Fig. 3D, Suppl. Fig. 3A-U).

**Figure 3:**
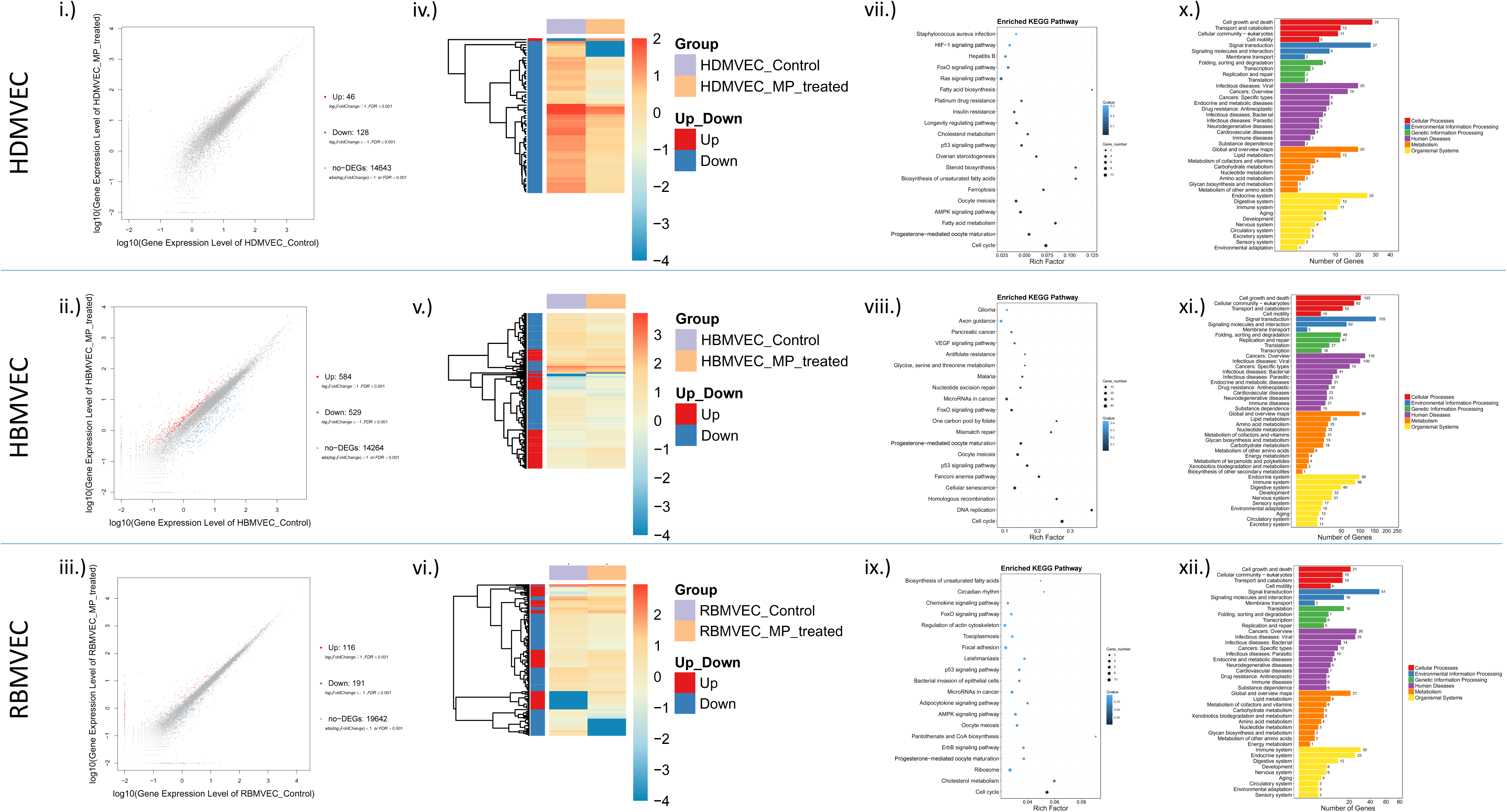
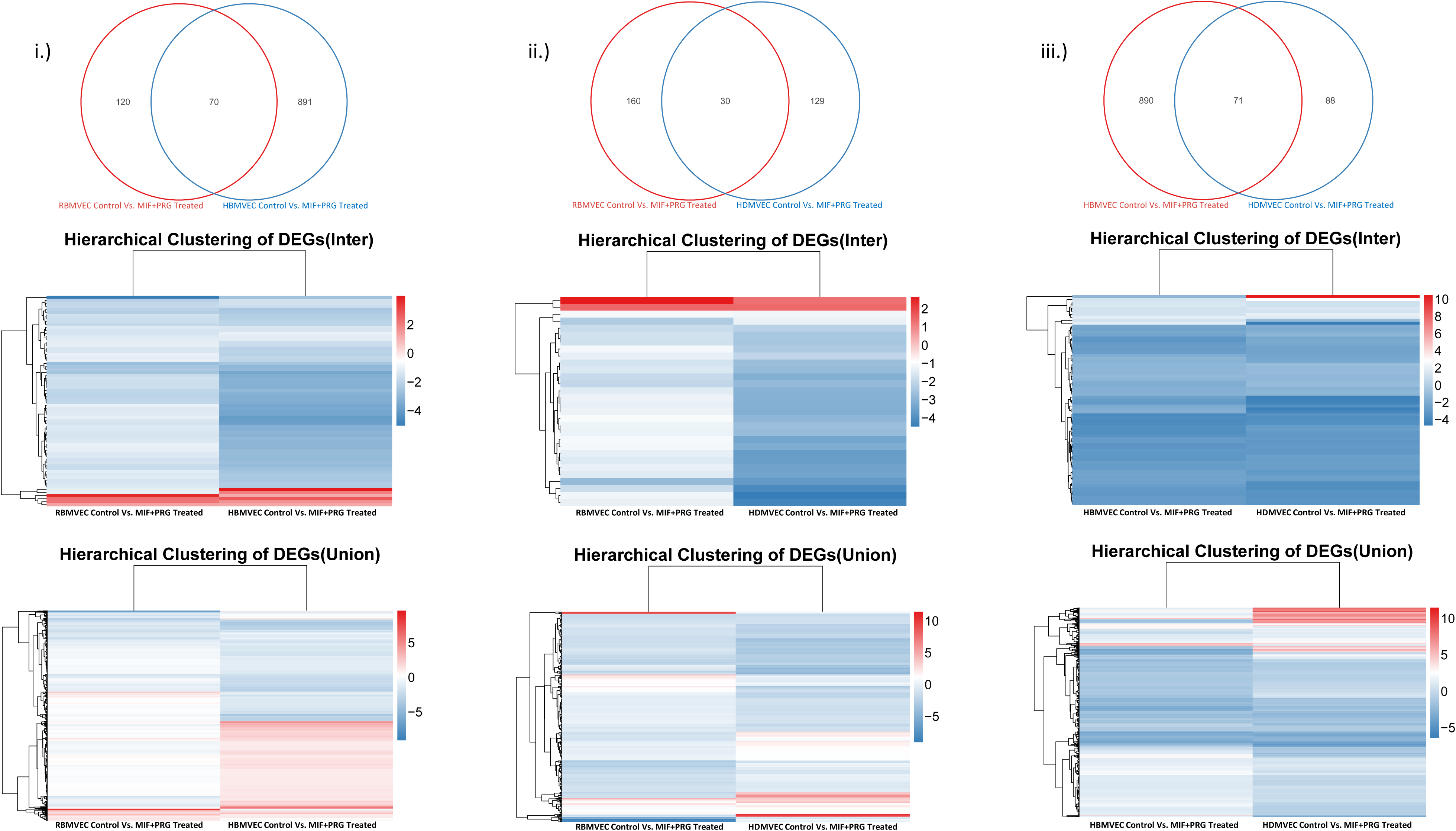
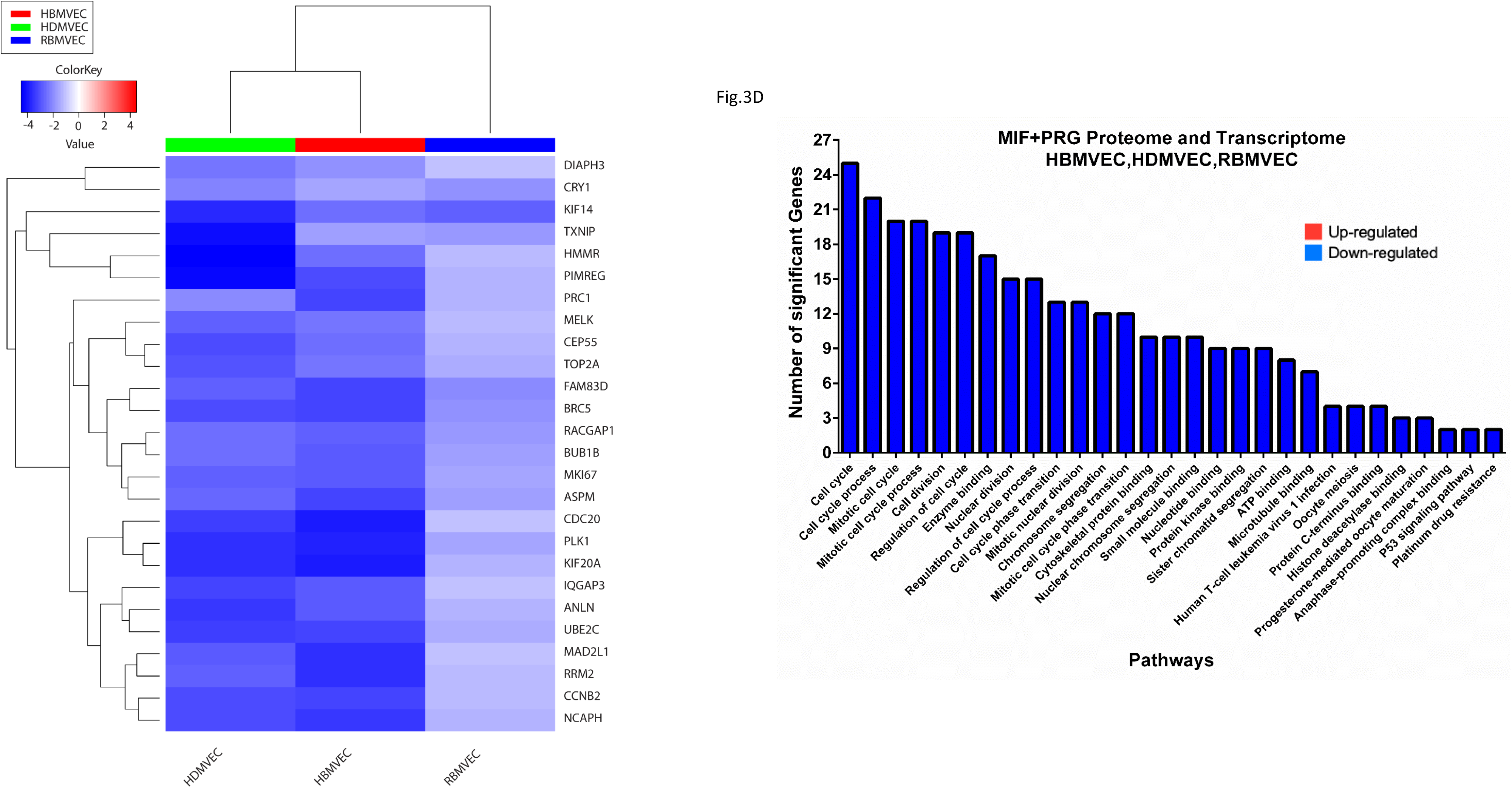
A. Differentially expressed gene (DEG) detection of MIF+PRG treatment on various endothelial cell lines through high-throughput RNAseq: (*i-iii*) Scatter plots of DEGs for MIF+PRG treated HDMVEC cells *(i)* HBMVEC cells *(ii)* or RBMVEC cells *(iii)* is displayed; X axis represents log10 expression of control gene, while Y axis represents log10 expression of MIF+PRG gene expression. (iv*-vi*) Cluster software and Euclidean distance matrixes were used for the hierarchical clustering analysis of the expressed genes (RNA) and sample program at the same time to generate the displayed Heatmap of hierarchical clustering for the DEGs from HDMVEC cells *(iv)* HBMVEC cells *(v)* or RBMVEC cells *(vi)* of expression clustering scheme; x axis represents each comparing sample and Y axis represents DEGs. (*vii-ix*) Pathway functional enrichment for DEGs from HDMVEC cells *(vii)* HBMVEC cells *(viii)* or RBMVEC cells *(ix)*. X axis represents enrichment factor. Y axis represents pathway name. The color indicates the q-value (high: white, low: blue), the lower q-value indicates the more significant enrichment. Point size indicates DEG number (The bigger dots refer to larger amount). Rich Factor refers to the value of enrichment factor, which is the quotient of foreground value (the number of DEGs) and background value (total Gene amount). The larger the value, the more significant enrichment. (*x-xii*) Pathway classification for DEGs from HDMVEC cells *(x)* HBMVEC cells *(xi)* or RBMVEC cells *(xii)*. X axis represents number of DEGs. Y axis represents functional classification of KEGG pathways. There are seven branches for KEGG pathways: Cellular Processes, Environmental Information Processing, Genetic Information Processing, Human Disease (For animals only), Metabolism, Organismal Systems and Drug Development. **B. Comparisons of overlapped/unique DEGs of MIF+PRG treatment on various microvascular ECs:** DEGs were evaluated between endothelial cell lines to identify unique as well as overlapped DEGs to further understand shared altered signaling cascades under MIF+PRG treatment across three EC lines. Analysis between RBMVEC compared to HBMVEC (i), RBMVEC compared to HDMVEC (ii) and HBMVEC compared to HDMVEC (iii) is displayed. Cluster software and Euclidean distance matrixes were used for the hierarchical clustering analysis of the expressed genes (RNA) and sample program at the same time to generate the displayed Heatmap of hierarchical clustering for the intersection of DEGs or union of DEGs of expression clustering scheme (middle panels); x axis represents each comparing sample and Y axis represents DEGs. **C. Comparisons of overlapped DEGs of MIF+PRG treatment on RBMVEC, HBMVEC and HDMVECs:** DEGs were evaluated between endothelial cell lines to identify overlapped DEGs to further understand shared altered signaling cascades under MIF+PRG treatment across all three endothelial cell lines. Cluster software and Euclidean distance matrixes were used for the hierarchical clustering analysis of the expressed genes (RNA) and sample program at the same time to generate the displayed Heatmap of hierarchical clustering for the DEGs of expression clustering scheme (left panel); x axis represents each comparing sample and Y axis represents DEGs. Pathway functional enrichment results for up/down regulation of genes was also performed from the genes shown in the Heatmap (right panel); X axis represents the terms of Pathway, Y axis represents the number of DEGs, For all maps/graphs, Coloring indicates the log2 transformed fold change (high: red, low: blue) (for RAW data, please see suppl. tables 3A-3C).

#### Reciprocal regulations between the CSC and mPRs in PR(-) ECs at both the transcriptional and translational levels

To investigate the role of the CSC in HBMVECs under steroid actions at the translational level, Identified differentially expressed genes/proteins (DEGs/DEPs) were also compared to our previously published omic data generated from *CCMs* (*1, 2, 3*) genes knockdown (KD) experiments in HBMVECs (*2, 3*). Our proteomics data clearly demonstrated overlaps between MIF+PRG treatment and CCMs (1, 2, 3) KD HBMVEC cells as illustrated in the Venn diagrams (Fig. 4A, panels i-iii and Suppl. Table 4A), emphasizing the existence of CSC-mPRs coupled signaling, and the importance of this feedback-regulated signaling cascade in microvascular angiogenesis. To further delineate the role of the CSC and mPRs/PAQRs under steroid actions in HBMVECs, we compared our two omic approaches to identify altered signaling pathways at both the transcriptional and translational levels. First, we identified overlapped, as well as unique genes/proteins at both levels for hormone treatment, and discovered 10 overlapped DEGs/DEPs (Fig. 4B and Suppl. Table 4B). Pathways analysis of the 10 overlapped DEGs/DEPs revealed steroid actions influence several key metabolic processes, cellular homeostasis, regulation of cell cycle arrest, and steroid hormone receptor binding (Fig. 4B, top right panel and Suppl. Table 4C) signaling pathways at both the transcriptional and translational levels in HBMVECs. To further delineate the role of the CSC, involved with mPRs/PAQRs under steroid actions in HBMVECs, we compared our two omic approaches to identify altered signaling pathways at both the transcriptional (hormone treatment) and translational levels (disrupted CSC). We identified 17 DEGs/DEPs among hormone treatment (RNAseq) with a disrupted CSC (pooled, proteomics) (Fig. 4C and Suppl. Table 4B). Functional enrichment performed from these 17 genes demonstrated alterations to key pathways including several cellular response processes, cytokine response, ovulation, IL-12 mediated signaling, and positive regulation of blood vessel endothelial cell migration signaling pathways (Fig. 4C, bottom right panel and Suppl. Table 4C). Based off of these data, we hypothesize that the stability of CCMs proteins in vascular ECs will be the first to be challenged when steroid hormone directed reciprocal signal transduction network between the CSC and mPRs/PAQRs is perturbed. This finding translates to the working hypothesis that under perturbed steroid hormone homeostasis/biogenesis, malformed and leaky vasculatures would be among the first group of anomalies to be detected in animal models. We then performed subsequent *in-vivo* analysis to test this hypothesis.

**Figure 4:**
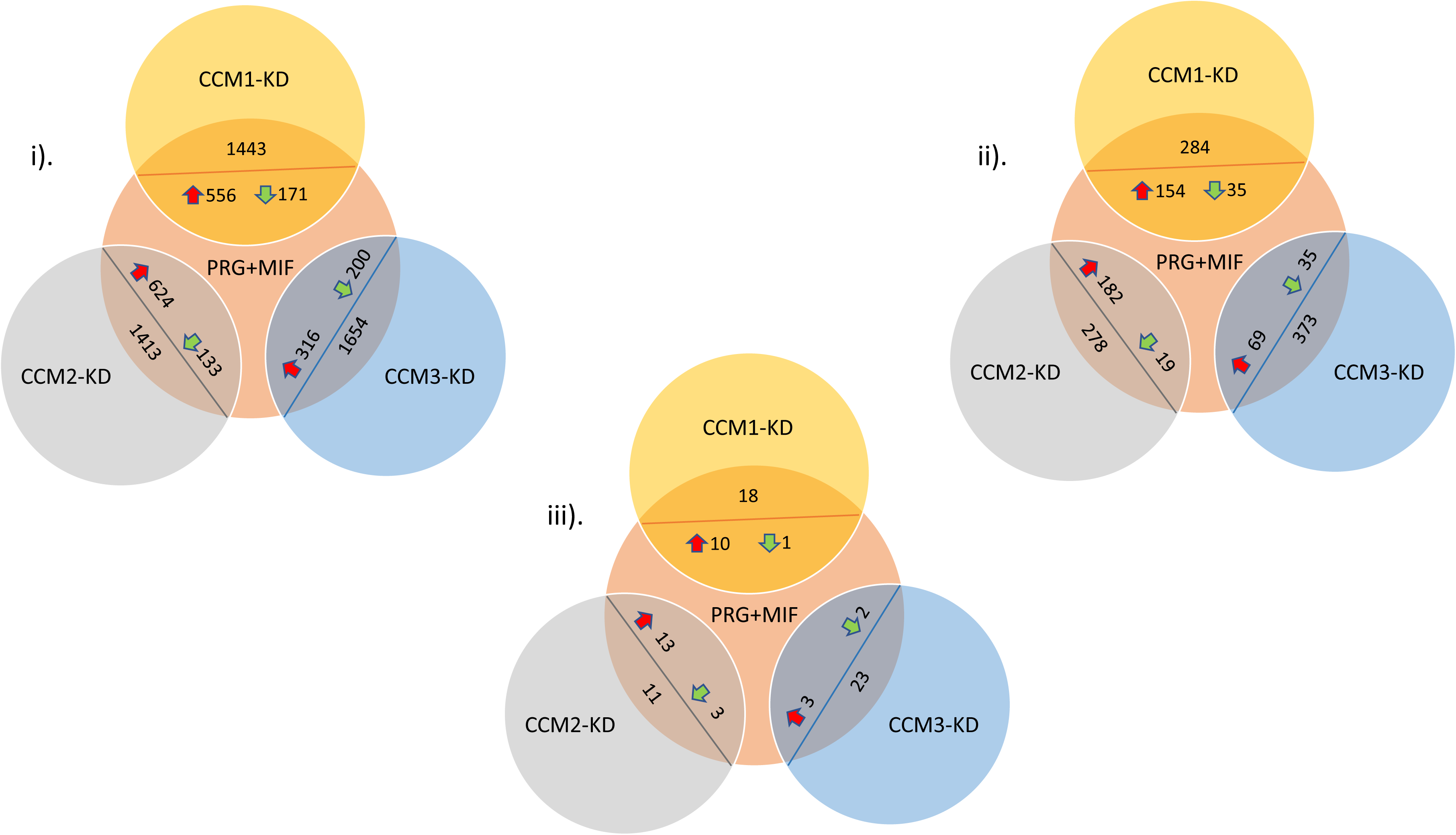
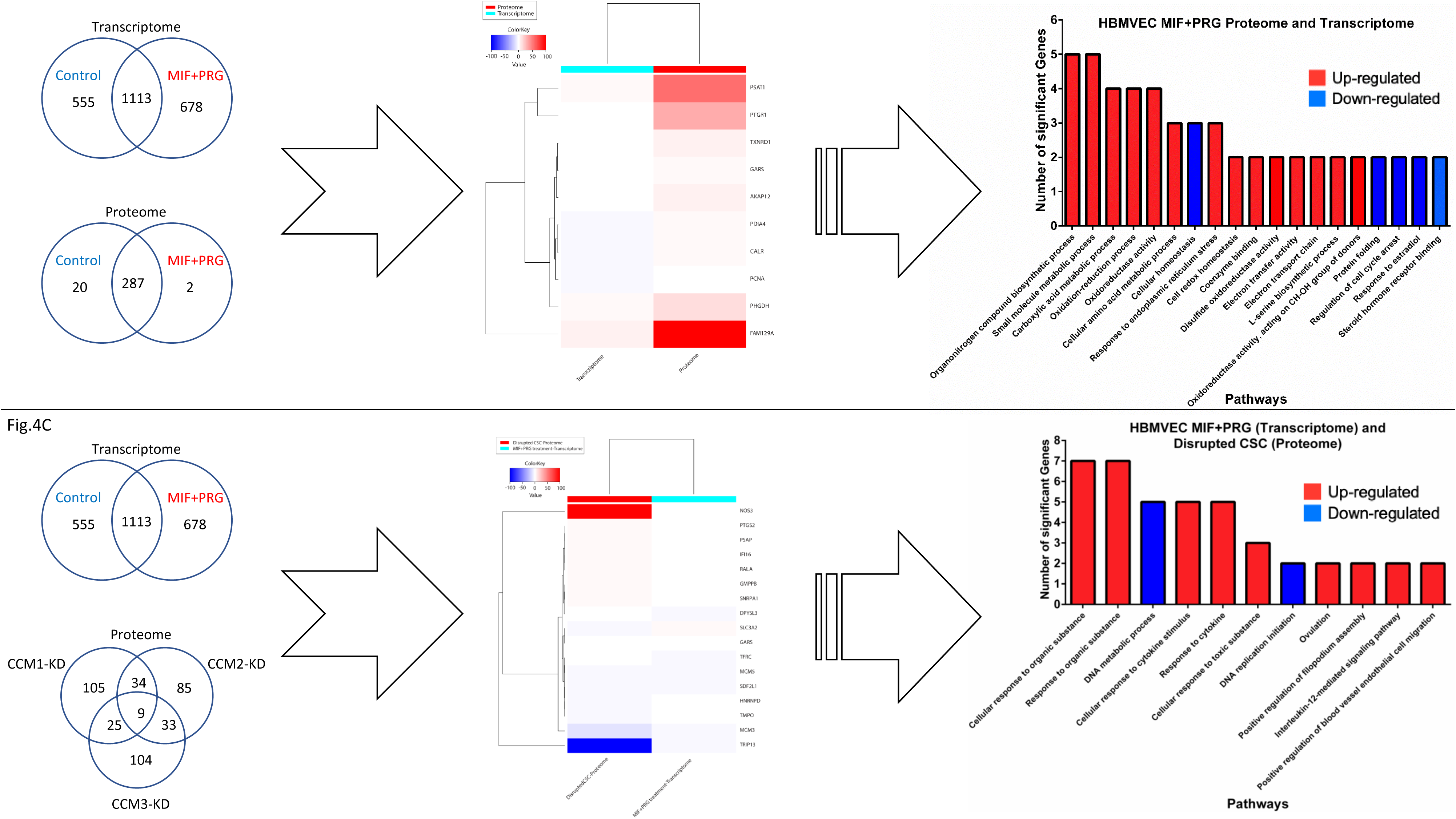
Shared DEGs and differentially expressed proteins (DEPs) between hormone treatment and a disrupted CSC in HBMVECs. A. DEPs detected with MIF+PRG treatment compared with a disrupted CSC in HBMVECs through LC-MS/MS: Venn diagram shows the overlapped DEPs between *CCMs (1, 2, 3)* genes knockdown and PRG+MIF treatment in HBMVECs. DEPs were extracted from the raw data and analyzed for similarities in fold changes between *CCMs (1, 2, 3)* genes knockdown/SC controls and PRG+MIF treated/Vehicle controls. Fold changes were calculated from the original corresponding controls (KD vs WT, and MIF+PRG vs vehicle). i) All DEPs between two groups regardless of significance; ii). DEPs between two groups with at least significance in one treatment group; iii). DEPs between two groups with significance in both treatment groups **B. Comparisons of overlapped DEGs/DEPs of MIF+PRG treatment and disrupted CSC in HBMVEC at the proteome and transcriptome levels:** DEGs were evaluated between HBMVECs at the proteome and transcriptome level to further understand shared altered signaling cascades under MIF+PRG treatment (*Fig. 4B, left upper panel*). Cluster software and Euclidean distance matrixes were used for the hierarchical clustering analysis of the expressed genes (RNA and protein) and sample program at the same time to generate the displayed Heatmap of hierarchical clustering for the DEGs of expression clustering scheme (Fig. 4B, *middle panel*). Pathway functional enrichment results for DEGs was also performed from the genes shown in the Heatmap (Fig. 4B, *right panel*). **C. Comparisons of overlapped DEGs/DEPs between HBMVECs at the proteome and transcriptome level.** To further understand shared altered signaling cascades under MIF+PRG treatment (Transcriptome) and a disrupted CSC (Proteome) (Fig. 4C *left lower panel*). Cluster software and Euclidean distance matrixes were used for the hierarchical clustering analysis of the expressed genes (RNA and protein) and sample program at the same time to generate the displayed Heatmap of hierarchical clustering for the DEGs/DEPs of expression clustering scheme (Fig. 4C, *middle panel*). Pathway functional enrichment results for DEGs/DEPs was also performed from those shown in the Heatmap (*Fig. 4C, right lower panel*). For heat maps, x axis represents each comparing sample and Y axis represents DEGs. For pathways functional enrichment graphs, X axis represents the terms of Pathway, Y axis represents the number of DEGs/DEPs. For all maps/graphs, coloring indicates the log2 transformed fold change (high: red, low: blue) (for RAW data, please see suppl. tables 4A-4C).

### Perturbation of sex hormones modulates mPRs-CSC signaling cascade, which is sufficient for initiation of hemorrhagic events during the pathogenesis of CCMs

It has been proven that falling below the threshold of haploinsufficiency of CCMs proteins in microvascular ECs is an essential step for the pathogenesis of CCM lesions *in-vivo*, generated from both zebrafish (*23, 24*) and mouse Ccms mutant models (*25–27*). Approximately 40% of patients with CCMs have the sporadic form of the disease (*28*), which is difficult to be explained by the current dominant “two-hit” model (*29–31*). Clonal expansion data provided some degree of support for this “two-hit” model (*32, 33*). However, this haploinsufficiency or even null condition of CCM has further been proven to be insufficient for the initiation of hemorrhagic events of CCM lesions in Ccms mutant mouse models (*34*), making this “two-hit” model less likely as a possible common ignition of hemorrhage. Therefore, there must be a molecular “trigger” to initiate the hemorrhagic events of CCM lesions. Based on our *in-vitro* data, we tested the effects of sex hormone (PRG+MIF) actions on the pathogenesis of CCM lesions in mouse WT, Ccm1, Ccm2, and Ccm3 mutant strains.

#### Early appearance of increased permeability of blood vessels in the liver and lung

Using Evans Blue dye (EBD), we quantitatively measured the permeability of blood vessels in the brain, liver, and lung, which are major sites for Ccm pathology and permeability studies (*35*). To test the presence of compromised permeability of blood vessels in the brain, liver, and lung, resulting from vehicle injection, we initially examined EBD data between naïve (untreated with no-injections, N) and vehicle (injection of vehicle, peanut oil) groups among WT, Ccm1, and Ccm2 mutant mice; similar intensity of EBD signals among these control groups were seen (Suppl. Fig. 5A), indicating no apparent influence of vehicle injection on the permeability of blood vessels in these three organs. We then measured permeability of blood vessels in the brain, liver, and lung among all mouse strains, after treatment with PRG+MIF for 30, 60, and 90-day periods, respectively. Increased permeability of blood vessels in the lung and liver were observed in Ccm2 mutant mice as early as 30-days but disappeared for this mutant in 60 and 90 day treatment groups (liver) but remained significantly increased in the lungs for our 60 day treatment group (middle and right panels, Fig. 5A). We also observed increased permeability in the lungs in our 60-day treatment group with CCM3 mutant mice, which remained elevated in our 90 day treatment group as well (middle panel, Fig. 5A). Interestingly, in Ccm3 mutants, increased permeability of blood vessels was seen in the liver for 60-day treatment group, but diminished in our 90-day treatment group, reaffirming the existence of an unknown compensatory mechanism (middle and right panels, Fig. 5A). These results suggest that there may be an innate compensation mechanism to counter the sex steroid induced-blood vessel disruption in Ccm2/3 mutant mice or another possibility is increased efficiency of lymphatic drainage of the liver and lungs over time.

**Fig. 5.**
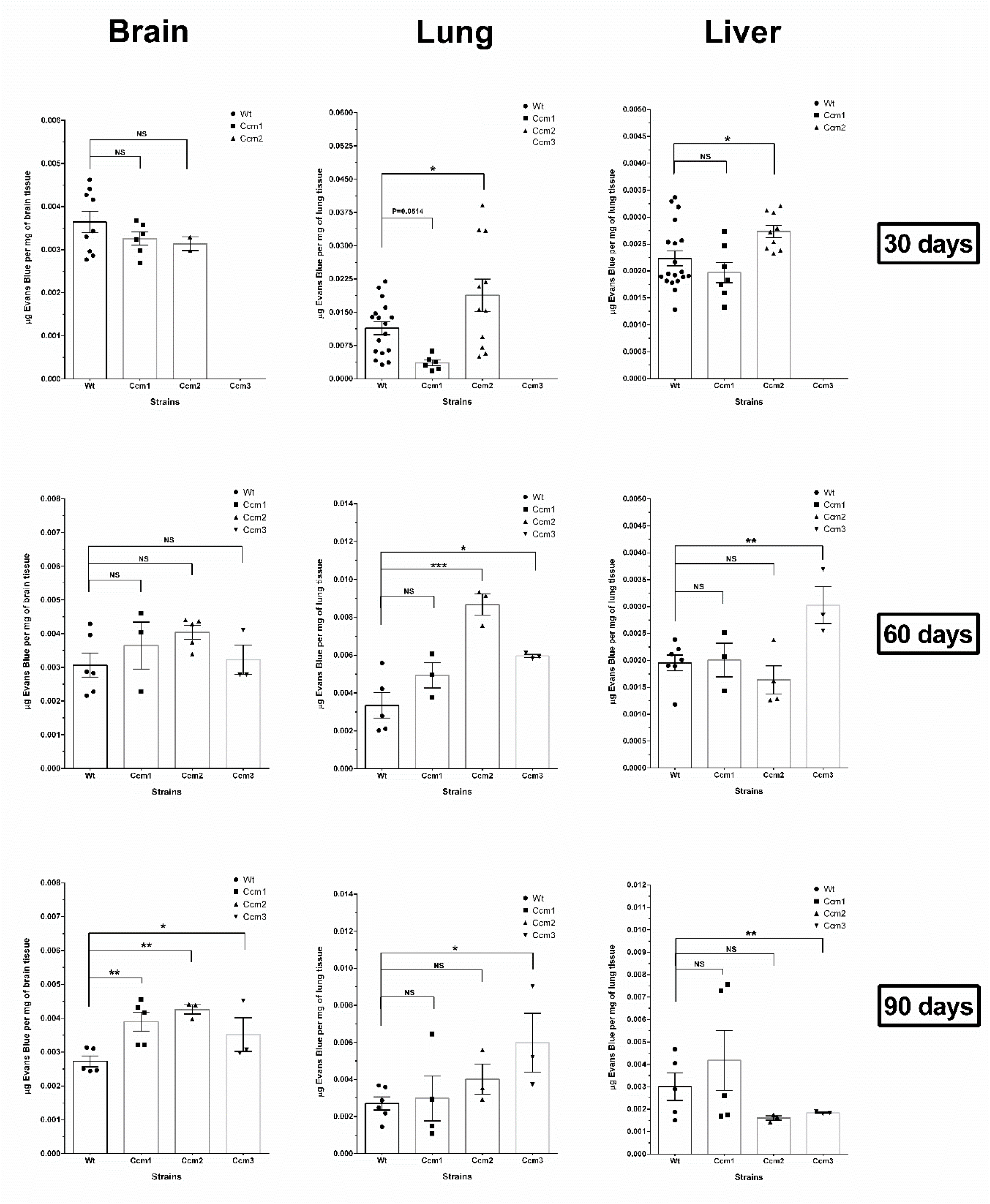
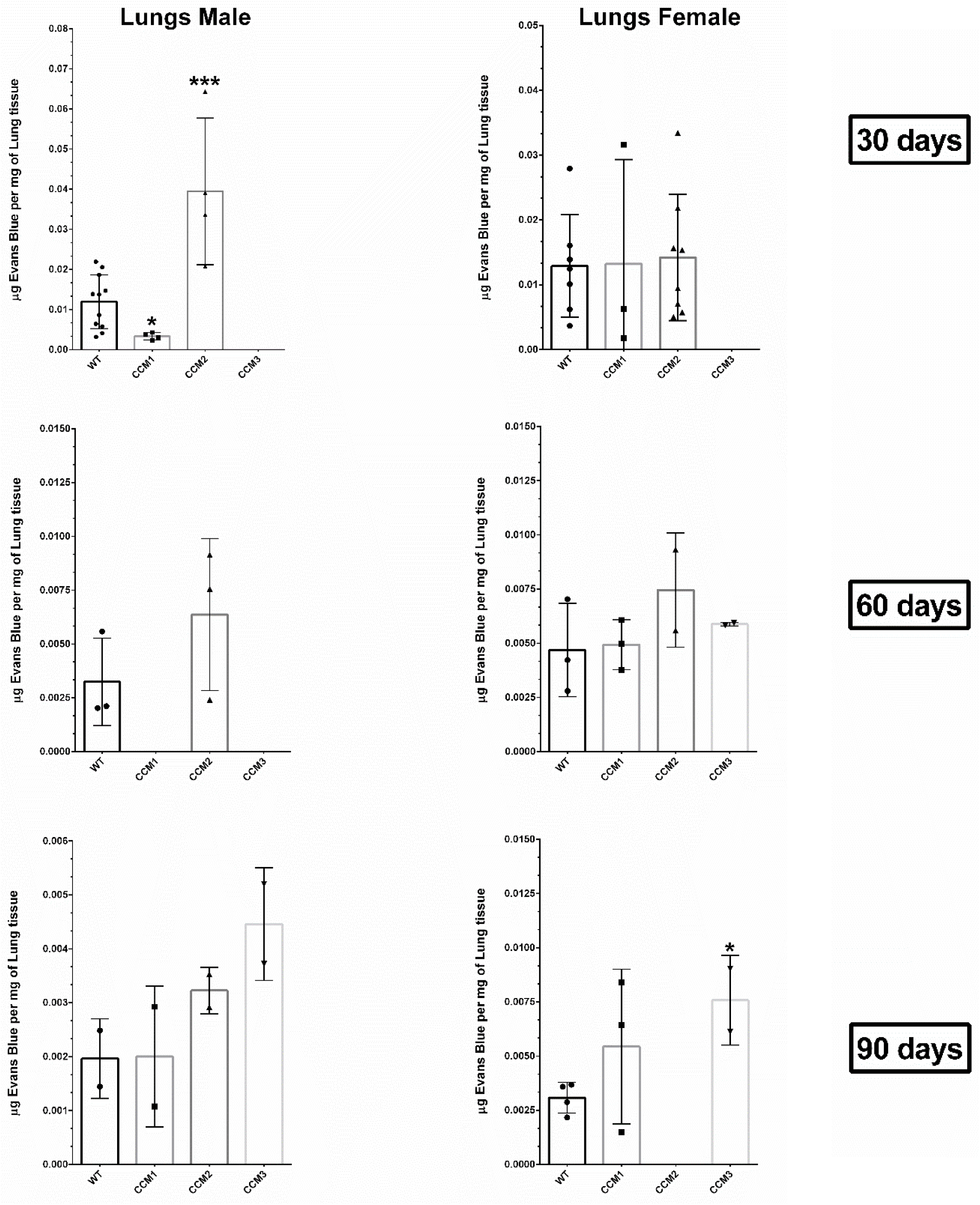
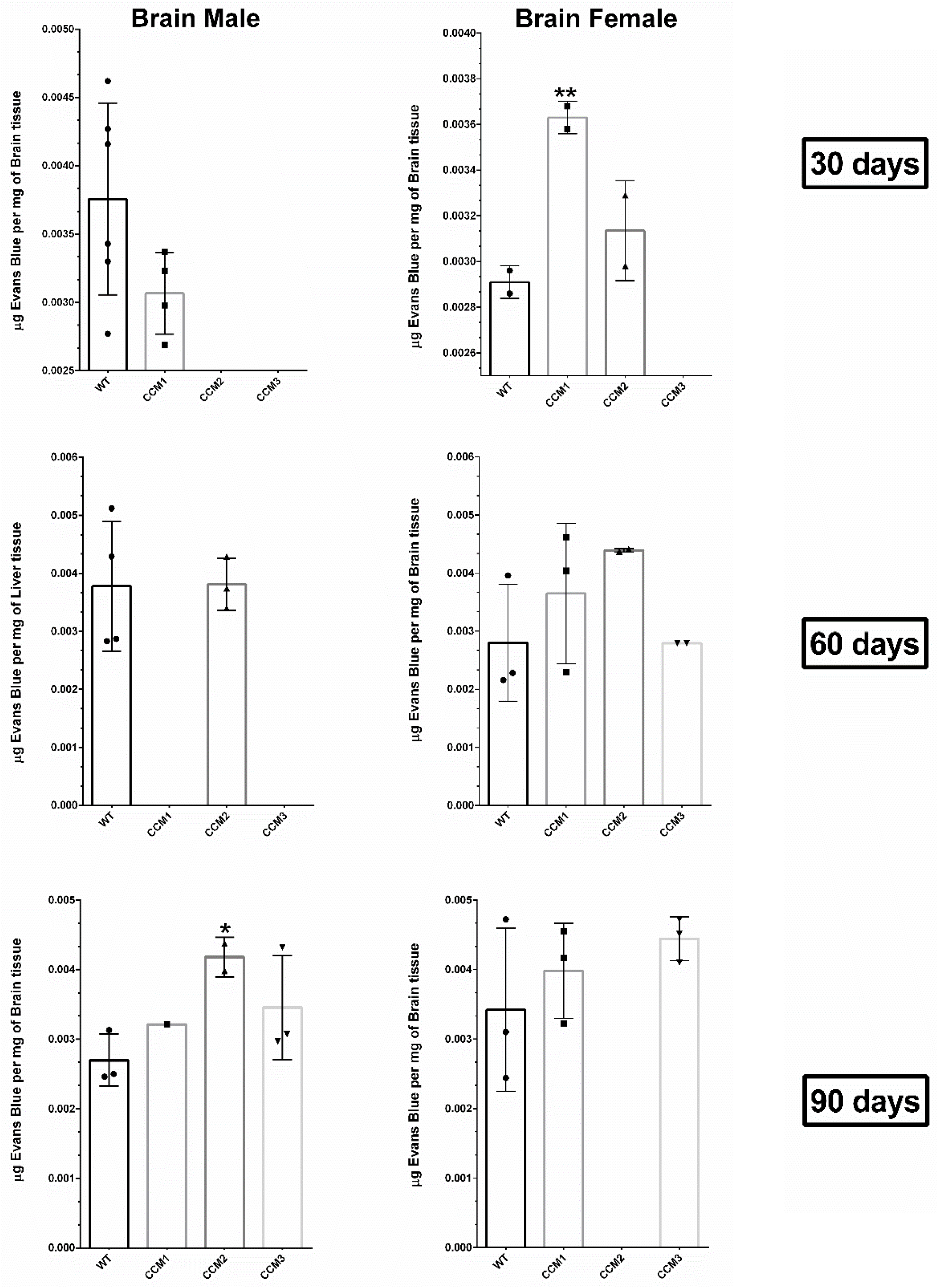
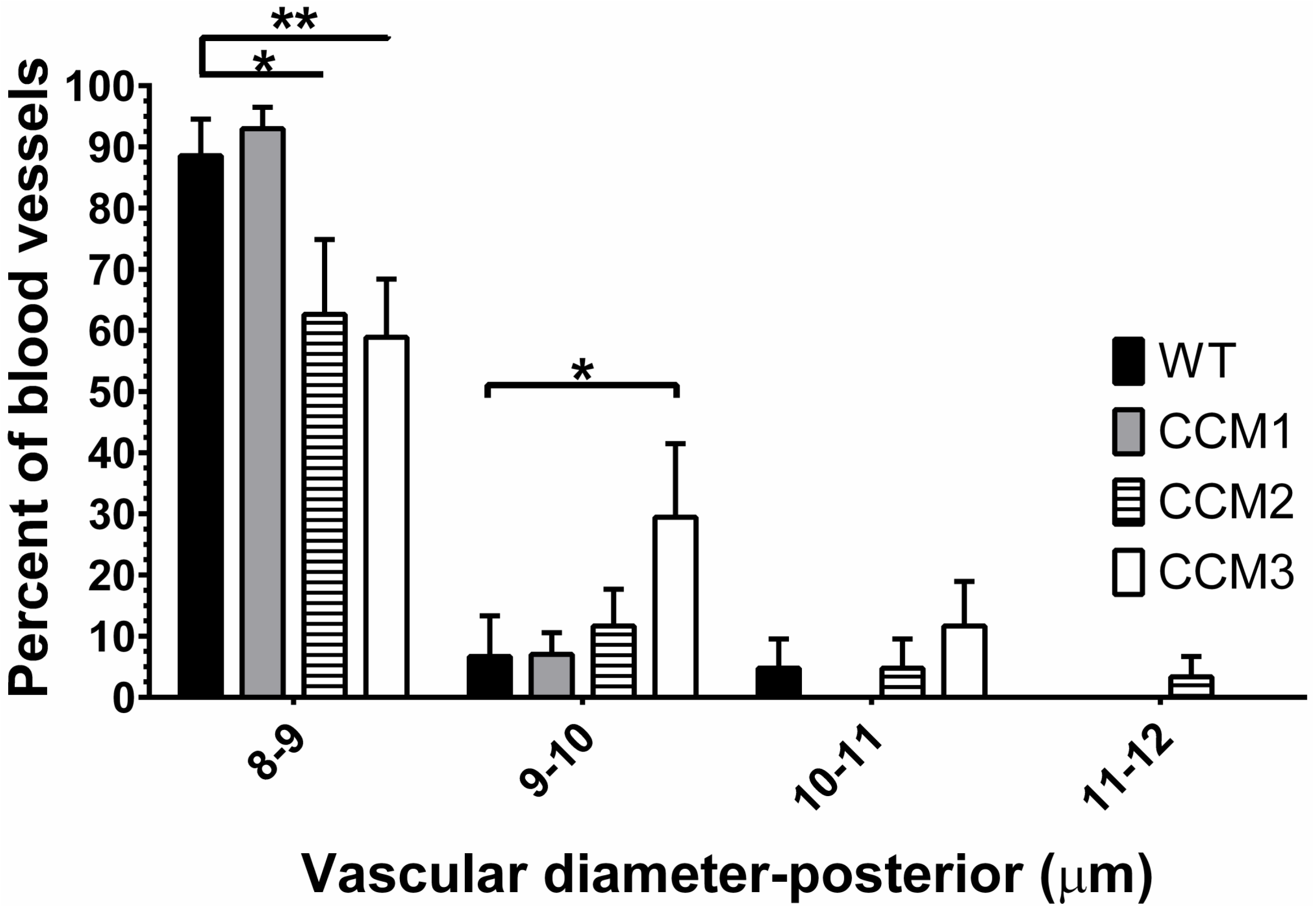
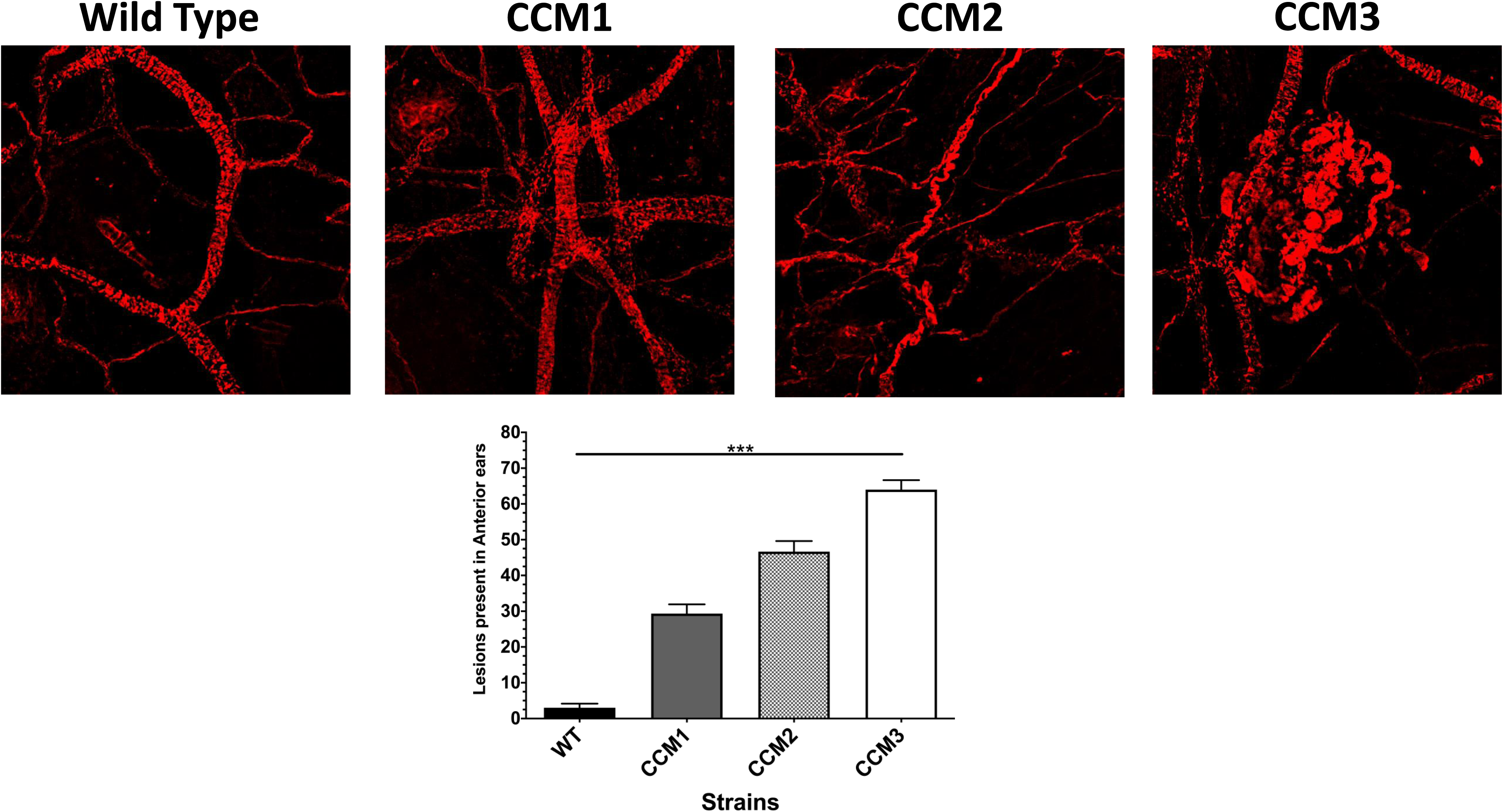
Steroid actions on PR(-) ECs increases microvascular permeability, and is sufficient for the formation of hemorrhagic CCM lesions *in-vivo*. Ccms (1, 2, 3) mutants and WT (C57BL/6J) mice were injected with a combined steroid treatment, a cocktail of progesterone + mifepristone (100 mg/kg body weight) in peanut oil (vehicle), 5 days a week for 30, 60, and 90 days respectively. **A.** Increased microvascular permeability among the lung, liver, and brain are associated with combined actions of PRG+MIF and genotypes. Microvascular leakage in the lung, liver, and brain were observed in PRG+MIF groups of Ccm mutants relative to WT mice. Increased microvascular permeability is detected earlier (30 or 60 days) in lung and liver for Ccm2/3 genotypes, respectively. Blood brain barrier (BBB) permeability is significantly increased at 90 days in all 3 Ccm mutants. **B.** Fluorescence data generated in the previous steroid treatment groups (Fig. 5A) were further stratified by genders. Significant gender differences among Ccms mutant strains were found in the increased microvascular permeability in the lungs. Significantly increased microvascular permeability in the lungs was first observed in male Ccm2 mice in one month (30days) treatment group, while significantly increased microvascular permeability in the lungs was only observed in female Ccm3 mice in three month (90days) treatment group. **C.** Gender differences were also found in the increased BBB permeability. Significantly increased BBB permeability was first observed in female Ccm1 mice in one month (30days) treatment group, while significantly increased BBB permeability was only observed in male Ccm2 mice in three month (90days) treatment group. **D.** Subcutaneous vessel diameters in posterior sections of ears were classified into four subgroups based on the range of the vessel size (diameters): group-I (8-9 µM), group-II (9-10 µM), group-III (10-11 µM), and group-IV (11-12 µM). Significantly decreased percentage of vessels were found in Ccm2/Ccm3 within group-I (8-9 µM), compared to WT in the 90 days PRG+MIF treatment group, suggesting that more vessels in Ccm2/Ccm3 mutants are distributed in larger diameter groups under steroid actions. Indeed, the significant increased percentage of larger vessels in Ccm3 mutant was found in group-II (9-10 µM) compared to WT, and Ccm2 is the only mutant to display vessels in the largest size group, group-IV (11-12 µM) under steroid actions. **E.** Subcutaneous vessel lesions in the anterior side of mouse ears in all Ccm (1, 2, 3) mutant strains can be visually distinguished in 90 day treatment group compared to WT, while CCM3 had the largest number of CCM lesions among Ccm mutants. More details provided in Suppl. Fig. 5G.

#### Late appearance of increased permeability of Blood Brain Barrier (BBB)

Intriguingly, all Ccms (1, 2, 3) mutants were responding uniformly to PRG+MIF treatment in regards to BBB permeability. Similar intensity of EBD signals among Ccm (1, 2, 3) mutants and WT mice for both 30 and 60-day treatment groups were seen, identical to the naïve and vehicle control groups, suggesting no influence of genotypes on the permeability of blood vessels yet (Left panels, Fig. 5A). However, significantly increased permeability of BBB was observed in all Ccms (1, 2, 3) mutant mice in 90-day treatment groups compared to WT mice, indicating that the BBB was initially resistant to the disruption pressure from MIF+PRG actions, but eventually the integrity of the BBB for all Ccms (1, 2, 3) mutants was compromised due to the long lasting effect of steroid actions.

#### Gender differences of disrupted permeability within blood vessels under sex steroid actions

To better understand the underlying mechanisms of the negative effects of PRG+MIF hormones on the permeability of blood vessels, WT and Ccms mutant mice were further stratified into male and female groups. In the lungs, increased permeability of blood vessels was observed in male Ccm2 mutant mice in 30-day treatment groups, but diminished after 60-day treatments, supporting a possible compensatory mechanism (Upper left panel, Fig. 5B). However, increased permeability of blood vessels was only observed in female Ccm3 mutant mice in 90-day treatment groups (Lower right panel, Fig. 5B), suggesting this increased permeability of blood vessels in the lungs may be an accumulative event in female Ccm3 mutants. Significant increased BBB permeability was only observed in female Ccm1 mutants for our 30-day treatment group, suggesting the vulnerability of the BBB to sex steroid hormones in female Ccm1 mutants (Upper right panel, Fig. 5C). However, increased BBB permeability was only observed in male Ccm2 mutant mice in our 90-day treatment group (Lower left panel, Fig. 5C), demonstrating that the BBB is able to initially resist the disrupting pressure from sex steroid hormones in males but is more susceptible for disruption earlier in females, as demonstrated by increased leakage in the 30-day treatment group. Overall, EBD data strongly indicate that the BBB integrity and permeability among females is more susceptible to excessive action of sex steroid hormones to initiate hemorrhagic events earlier than males. It is also important to note the sensitivity of alterations to BBB integrity, as the differences observed in our EBD data among our Ccms mutant mice seem minimal (although statistically significant), but the clinical impacts are drastic enough to induce early hemorrhagic events, such as intracranial bleeding (suppl. Fig. 5B), stroke/ loss of motor functions (Suppl. Video 1) and seizure clinical appearances (Suppl. Videos 2-4)) among multiple Ccms mutant mice in the MIF+PRG treatment groups.

#### Disrupted angiogenesis of blood vessels was further validated in ear vessels

The negative effects of sex steroid hormones on blood vessels was further validated utilizing a mouse ear angiogenesis assay. Microvessels in mouse ears were evaluated by measuring four different parameters: 1). Vascular density, 2). Vascular length density, 3). Vascular diameter, and 4). Number of vessel lesions present in each leaflet. No obvious differences between all four vessel parameters was detected among untreated and vehicle treated mice, suggesting morphologically equivalent vascular conditions among all untreated/vehicle strains (Suppl. Figs. 5C-F). Further, under MIF+PRG treatment, no differences in vascular density (Suppl. Fig. 5C) or vascular length density (Suppl. Fig. 5D), were identified by comparing Ccms mutants and WT mice, suggesting no defects in these vascular parameters. An apparent shift in distribution of vascular diameters from smaller to larger (Fig. 5D) as well as an increased number of vessel lesions (Fig. 5E) in Ccms mutant mice were observed, compared to WT, indicating vascular defects in these two parameters in Ccms hemizygous mice were induced under chronic sex hormone actions.

### Immunosuppression was observed in Ccms mutant mice

#### A local inflammatory response is induced by vehicle injections

Peritoneal inflammation is induced by vehicle (peanut oil) injections, independent of genotype and sex steroid hormone treatments. It has been reported that steroids can influence the gut microbiota and, perturbed gut microbiota can affect the brain through gut microbiota-brain axis (*36*). Further, gut microbiota disturbance has been reported to lead to gastrointestinal inflammation (*37*). Examination of the mice after PRG+MIF treatments revealed abdominal distention, suggesting that peritoneal inflammation may be present. To assess if there was local inflammation in the peritoneal cavity, leukocyte populations were analyzed from this compartment. Animals were euthanized, peritoneal cells were collected via lavage and stained with fluorescent antibodies to quantify myeloid cell populations. We identified monocytes (CD45.2^+^CD11b^+^ Ly6G^-^ Ly6C^high^), neutrophils (CD45.2^+^CD11b^+^ Ly6C^int^Ly6G^high^), large peritoneal macrophages (LPM; CD45.2^+^CD11b^+^ Ly6G^-^Ly6C^-^ F4/80^hi^MHC-II^lo^), and small peritoneal macrophages (SPM; CD45.2^+^CD11b^+^ Ly6G^-^Ly6C^-^F4/80^low^MHC-II^high^) (the FACS gating pipeline is shown in Suppl. Fig. 6A).

We quantified the percentage (Fig. 6A I-IV) and absolute number (Fig. 6A V-VIII) of monocytes, neutrophils, LPM, and SPM in the peritoneal lavage. Naïve mice exhibited almost no monocytes or neutrophils in their peritoneal cavity (Fig. 6A I-II, V-VI), attributed to lack of injections. The majority of myeloid cells present were tissue resident macrophages, including an abundant LPM population and a less abundant SPM population (Fig. 6A III-IV, VII-VIII), consistent with prior reports (*38, 39*). Relative to naïve controls, WT mice injected with PRG+MIF, exhibited a significant increase in monocytes (percentage and absolute number) (Fig. 6A I, V), a significant increase in neutrophils (percentage and absolute number) (Fig. 6A II, VI), and a significant decrease in LPM (percentage and absolute number) (Fig. 6A III, VII). These cellular changes are similar to those observed in prior reports of peritoneal inflammation (*39–41*). Relative to naïve controls, WT-vehicle mice exhibited a significant increase in monocytes (percentage only) (Fig. 6A-I) a significant increase in neutrophils (percentage and absolute number) (Fig. 6A II, VI), and a significant reduction in LPM (percentage and absolute number) (Fig. 6A III, VII). These data suggest that there is a local inflammatory response to vehicle. Relative to vehicle controls, mice injected with PRG+MIF exhibited an increase in the percentage of monocytes in the peritoneal lavage, which was statistically significant (Fig. 6A I), suggesting that PRG+MIF may boost monocyte infiltration. However, the absolute number of monocytes, and the percentage and number of neutrophils and LPM were similar in the PRG+MIF and vehicle groups (Fig. 6A II-III, VI-VII). SPM were not substantially altered by any treatment (Fig. 6A IV, VIII).

**Fig. 6.**
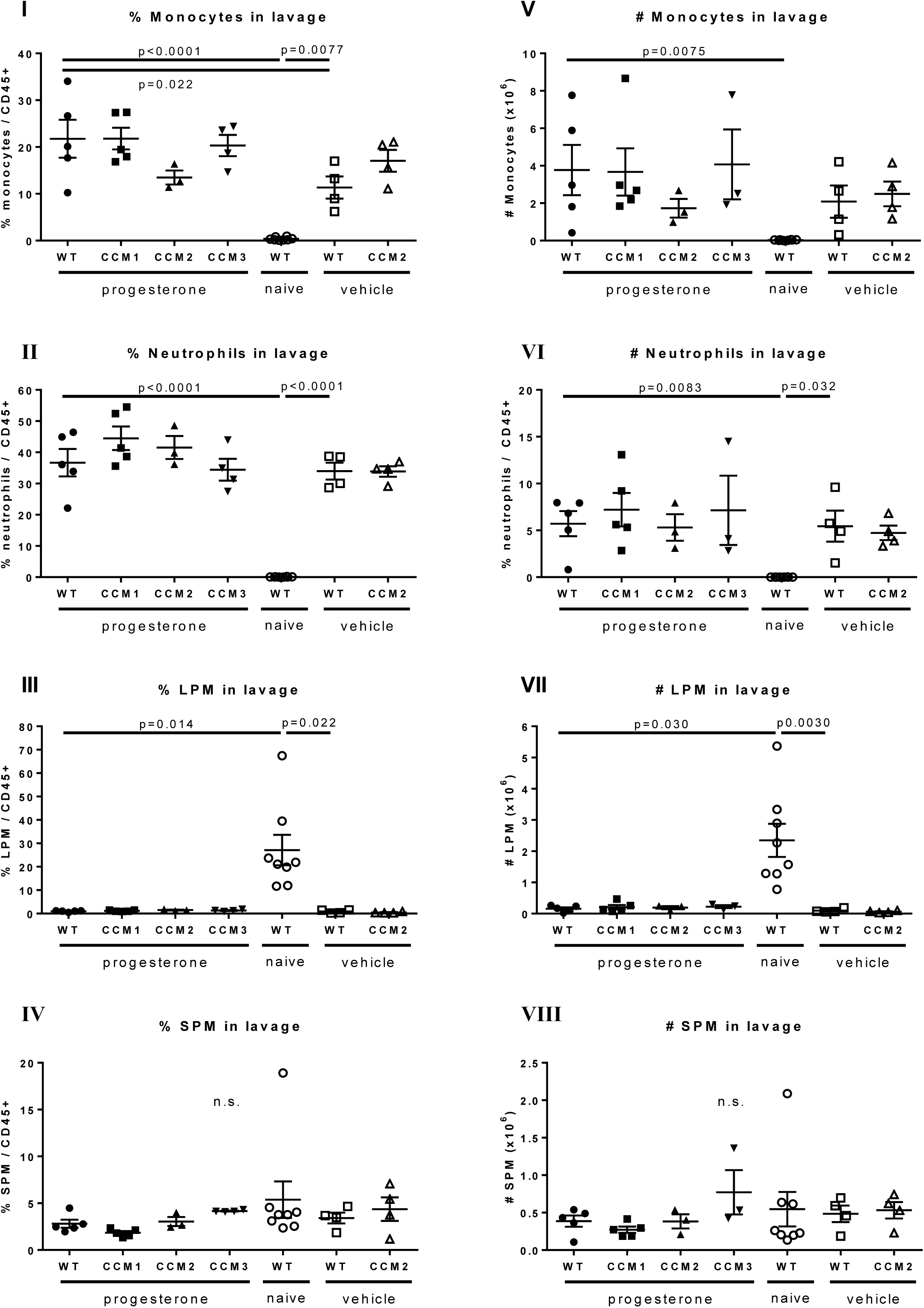
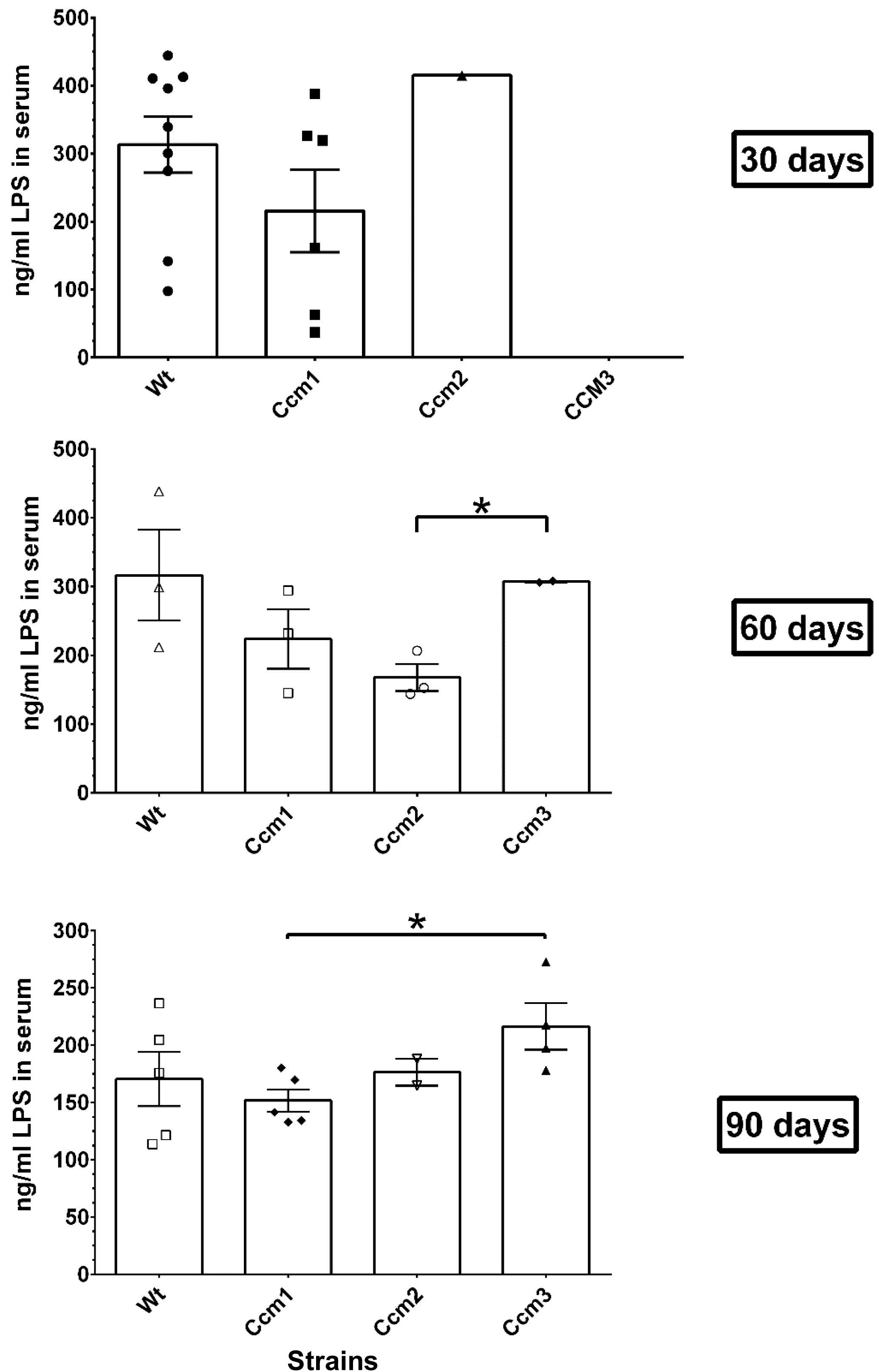
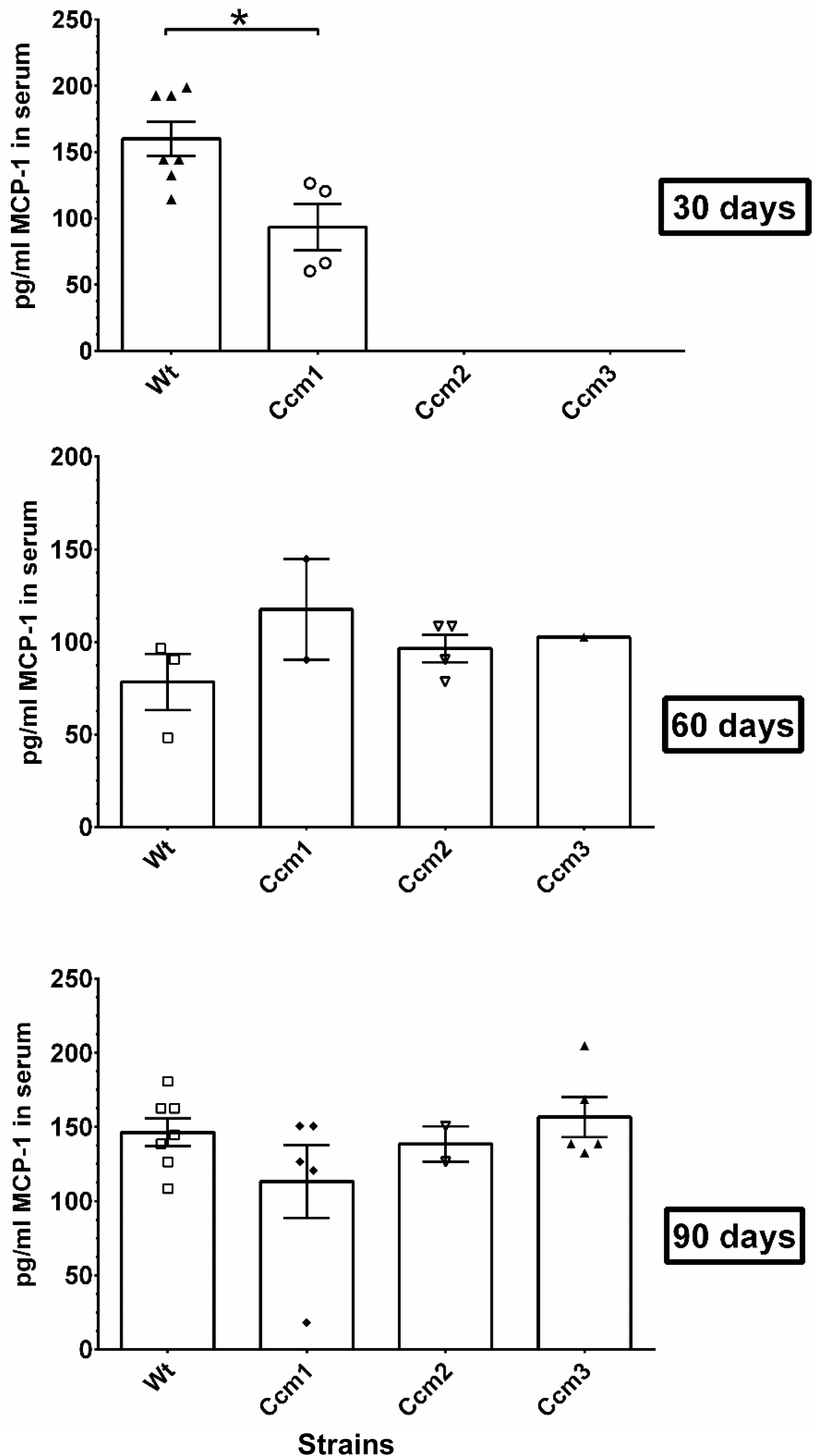
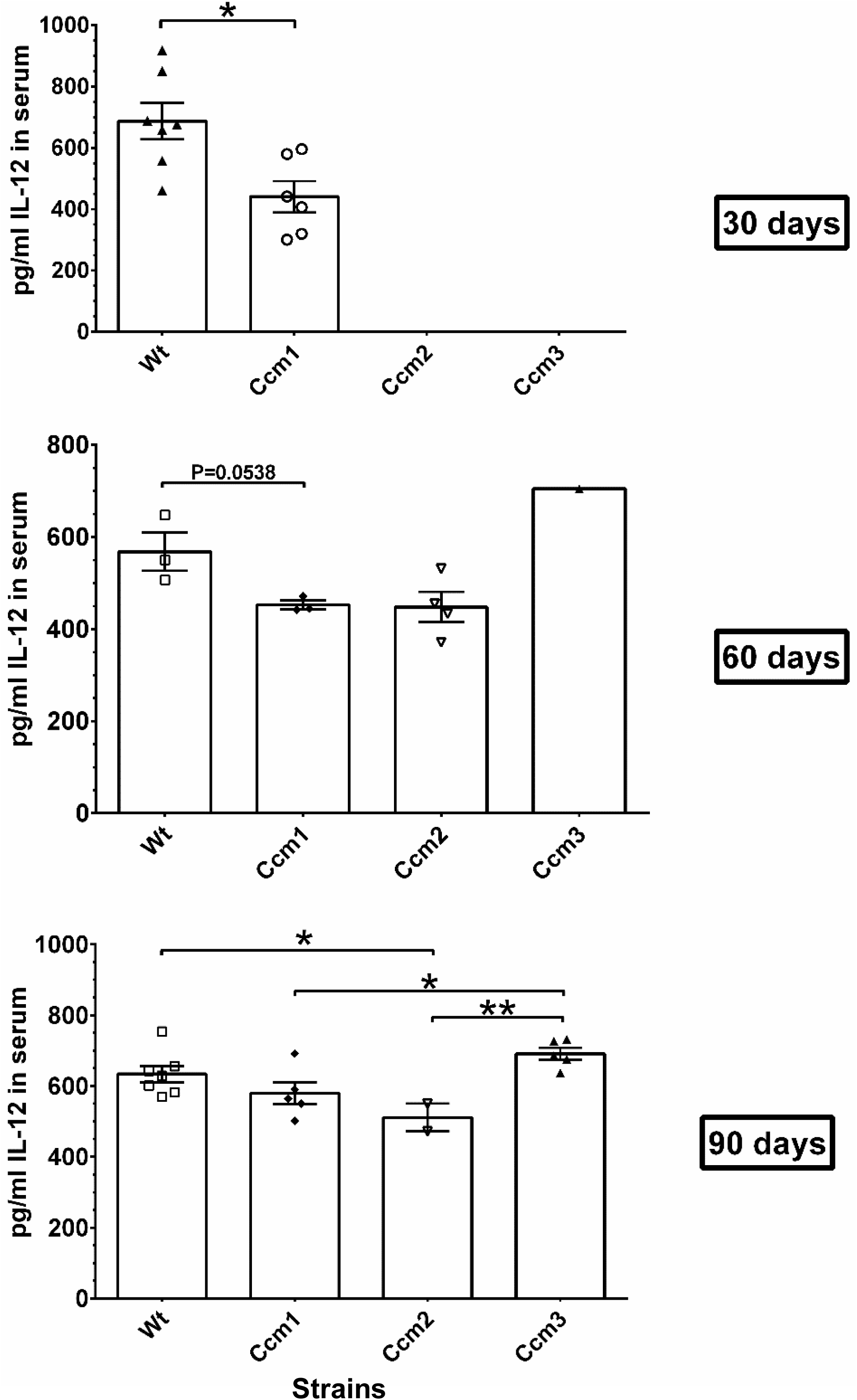
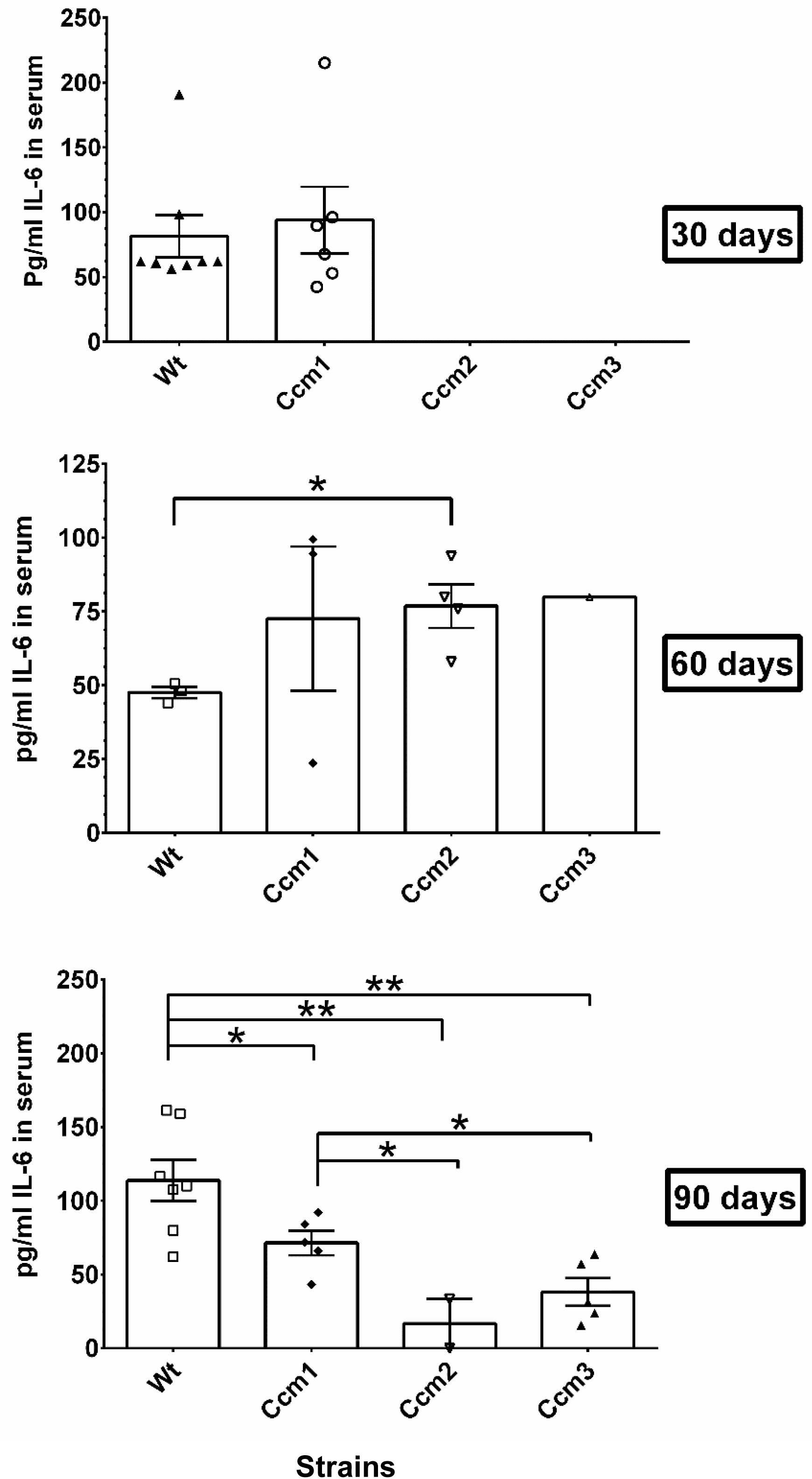
The disrupted brain-blood barrier (BBB) is not caused by local inflammatory reactions in Ccms mutant mice. Localized and systemic inflammations are not caused by either steroid actions or genotypes. **A.** Inflammation in the peritoneal cavity is not caused by either steroid actions or genotypes. Monocytes, neutrophils, large peritoneal macrophages (LPM), and small peritoneal macrophages (SPM) within the peritoneal cell populations were collected through peritoneal lavage. Quantification of percentage of I) Monocytes (CD45.2+CD11b+ Ly6G-Ly6Chigh) II) neutrophils (CD45.2+CD11b+ Ly6CintLy6Ghigh) III) LPM (CD45.2+CD11b+ Ly6G-Ly6C-F4/80hiMHC-IIlo) and IV) SPM (CD45.2+CD11b+ Ly6G-Ly6C-F4/80lowMHC-IIhigh) in peritoneal lavage of mice. The percentage of monocytes and neutrophils was calculated relative to CD45+ leukocytes. Quantification of absolute number of V) Monocytes (CD45.2+CD11b+ Ly6G-Ly6Chigh) VI) neutrophils (CD45.2+CD11b+ Ly6CintLy6Ghigh) VII) LPM (CD45.2+CD11b+ Ly6G-Ly6C-F4/80hiMHC-IIlo) and VIII) SPM (CD45.2+CD11b+ Ly6G-Ly6C-F4/80lowMHC-IIhigh) in peritoneal lavage of mice was performed. Dot plots show mean ± SEM (n=3-8). The only significance that was found was with naive wildtype (untreated), which did not receive any injections. This suggests that vehicle (peanut oil) is causing local inflammatory response. **B.** The disrupted brain-blood barrier (BBB) is not associated with non-immunogenic LPS from *Helicobacter pylori*. Lipopolysaccharide-Based Enzyme-Linked Immunosorbent Assay (LPS-ELISA) was used to measure LPS concentration in mouse serum. Nearly equal amounts of low LPS in the serum of all mouse strains were observed in 30, 60, and 90-day groups with PRG+MIF treatment, suggesting that levels of non-immunogenic LPS is not influenced by either sex steroid actions or genotypes. Relative higher amounts of non-immunogenic LPS in Ccm3 mutant mice were constantly observed from Naïve mice to 90-day treatment group, which also indicates irrelevance of quantity of non-immunogenic LPS towards BBB integrity. **C.** The serum level of inflammatory cytokine, MCP-1, is low and influenced initially by steroid actions in CCM1 mutant only. Nearly equal amounts of MCP-1 in the serum of all mouse strains were observed in 30, 60, and 90-day treatment groups with PRG+MIF but with some notable changes suggesting that MCP-1 may be influenced by either sex steroid action or genotypes at early stages in CCM1 mutant mice. **D.** The serum level of inflammatory cytokine, IL-12, is suppressed by steroid actions and genotypes. Significantly different amounts of IL-12 in the serum of mouse Ccm mutants (Ccm1/2) strains were observed in 30 and 60 (CCM1), and 90-day groups (CCM2) with PRG+MIF treatment respectively, suggesting that IL-12 is suppressed in mouse Ccm mutant (Ccm1/2) strains. **E.** The serum level of inflammatory cytokine, IL-6, is low and not influenced by either sex steroid action or genotypes. Significantly different amounts of IL-6 in the serum of mouse Ccm mutant (1, 2, 3) strains were observed in 90-day group of PRG+MIF treatment, suggesting that IL-6 expression is suppressed in mouse Ccm mutant (1, 2, 3) strains. Although higher amounts of IL-6 were observed in Ccm2 mutant group compared to WT, in 60-day PRG+MIF treatment group, this trend is reversed at 90 days. Significantly decreased amounts of IL-6 were observed in all Ccm mutants compared to WT, in 90- day groups of PRG+MIF treatment, suggesting that IL-6 is initially increased upon treatment but then is

To assess the possible effects of mouse genotype on this process, leukocyte populations were analyzed in the peritoneal cavity of all mouse strains injected with PRG+MIF (Fig. 6A I-VIII). The percentage and number of monocytes (Fig. 6A I, V), neutrophils (Fig. 6A II, VI), LPM (Fig. 6A III, VII) and SPM (Fig. 6A III, VII) was similar among all genotypes. Vehicle treated Ccm mutant mice exhibited a similar response to vehicle treated WT mice, suggesting that peanut oil drives peritoneal inflammation, irrespective of either genotype or sex steroid hormones. These data suggest that mouse CCM genotype does not influence the local inflammatory response observed.

#### Disrupted BBB is not associated with non-immunogenic LPS from H. pylori bacteria

Upon arriving, all our mouse strains were tested positive for murine norovirus, protozoan *tritrichomonas,* and mostly several strains *of Helicobacter pylori* which usually infects the digestive tract. Possible infection of *H. pylori,* a gram-negative bacteria could lead to elevated LPS levels in mouse serum, but is considered to be non-immunogenic (*42–44*). No obvious differences of LPS was detected among untreated (Naïve) and vehicle treated groups, suggesting equal existence of low LPS levels in the serum of all mouse strains (Suppl. Fig. 6B). No significant differences were observed, compared to WT, in the serum of all mouse strains among 30, 60, and 90-day treatment groups respectively, suggesting that existing non-immunogenic LPS in the serum is not influenced by steroid actions (Fig. 6B). Relative higher amounts of non-immunogenic LPS was observed in Ccm3 mutant mice compared to treated groups in both 60 and 90-day treatment groups (Fig. 6B), suggesting Ccm3 mutants may be more susceptible to *H. pylori* bacterial infections.

#### Systemic inflammations remain low in mice injected with either peanut oil or PRG+MIF treatment

Infection of murine model with norovirus and *tritrichomonas* allows us to screen for circulating cytokines to evaluate their expression upon MIF+PRG treatment. Four key markers of systemic inflammation (TNF-α, MCP-1, IL-12 and IL-6) were detected in the serum from all mouse strains.

To assess the possible effects of mouse genotype and injection procedures on this process, four cytokines were analyzed among all mouse strains under naïve (untreated) and vehicle [vehicle (peanut oil)-treated] conditions. No obvious differences were detected among them (Suppl. Figs. 6C-6F), suggesting low existing systemic inflammation, irrespective of either genotype, but possibly related to murine norovirus, or protozoan *tritrichomonas* (or peanut oil). No significant difference was observed in TNF-α, quantities in the serum of all mouse strains among 30, 60, and 90-day treatment groups respectively (Suppl. Fig. 6C), suggesting low existing TNF-α, irrespective of either genotype or sex steroid hormones.

#### Immunosuppression was observed in CCM deficient mice

Interestingly, significant decreased expression of MCP1 was noted in the serum of Ccm1 mutant mice in the 30-day treatment group only, suggesting possible early immunosuppression due to steroid treatment (Fig. 6C). IL-12 was also observed to be significantly decreased in the serum of Ccm1 mutant mice among 30-day treatment group, and though there is a trend of decreased IL-12 in Ccm1 mutant mice at 60 and 90-days, the differences are non-significant (60 day group p=0.058). There was significantly decreased expression of IL-12 in the serum of Ccm2 mutant mice among 90-day treatment group (Fig. 6D), suggesting existing IL-12 cytokines were suppressed, due to the combination of Ccm mutant genotypes and sex steroid hormones. These results validate our functional enrichment analysis illustrating alteration in IL-12 mediated signaling shared between DEGs/DEPs in HBMVECs at the proteome and transcriptome levels with a disrupted CSC under steroid actions (Fig. 4C, right lower panel).

The levels of IL-6 were found to be decreased in the serum of all three Ccms mutant mice among the 90-day treatment group (Fig. 6E), suggesting IL-6 cytokines were suppressed, due to the combination of Ccm mutant genotypes and sex steroid hormones. These data indicate suppression of systemic inflammation occurred in all three Ccms mutant mice under sex steroid actions. These data confirm that the permeability of blood vessels is not driven by a systemic inflammatory response, but rather is an effect of chronic sex steroid hormone actions, which also causes immunosuppression. PRG was reported to play a major role in immunomodulatory action (*45*), likely through its rapid effects on human T cells (*46, 47*). It was found that PRG-induced immunosuppression is mediated neither through PR1/2 (*48–50*) nor by the glucocorticoid receptor (GR) (*51*). Since peripheral blood monocytes and T cells are PR(-) (*52*), non-genomic actions through mPRs are the only viable possibility to explain PRG-induced actions including immunosuppression (*53*). Therefore, the amounts of PRG in serum should be determined to investigate whether this immunosuppression is indeed caused by excessive active “free form” PRG in serum.

### Disruption of PRG homeostasis leads to hemorrhagic events in CCMs

#### Normal homeostasis of PRG, serpin A6 and albumin in the serum of naïve and vehicle-only groups

Over 98% of PRG in blood is believed to be stored and passively transported by plasma proteins (*54*), mainly two major PRG-binding proteins in serum, serpin A6 (bound ∼18% PRG) and albumin, (bound ∼80% PRG) and is physiologically inactive (*55–57*). To further investigate the underlying mechanisms of sudden elevated levels of active PRG in the serum, quantification of PRG, serpin A6 (SERPINA), and Albumin in the serum were performed from naïve and vehicle treated groups. No significant differences were found between naïve (non-injection) and vehicle treated groups for all mouse strains (Suppl. Figs. 7A-7C), suggesting total amounts of PRG, serpin A6 and albumin are evenly distributed among WT and Ccms mutant strains.

#### Excessive active PRG in serum of Ccm2 mutant mice in 90-day treatment group

Quantification of PRG in the serum among all treatment groups was also examined. No significant differences were observed among the different mice strains in the 30 and 60-day treatment groups (Fig. 7A), suggesting homeostasis/biogenesis of PRG is normal among different strains of mice up to two months of PRG+MIF treatment, irrespective of genotype. This indicates that only WT mice have homeostatic capacity of PRG and intrinsic elasticity of PRG metabolism during the full treatment regimen.

**Fig. 7.**
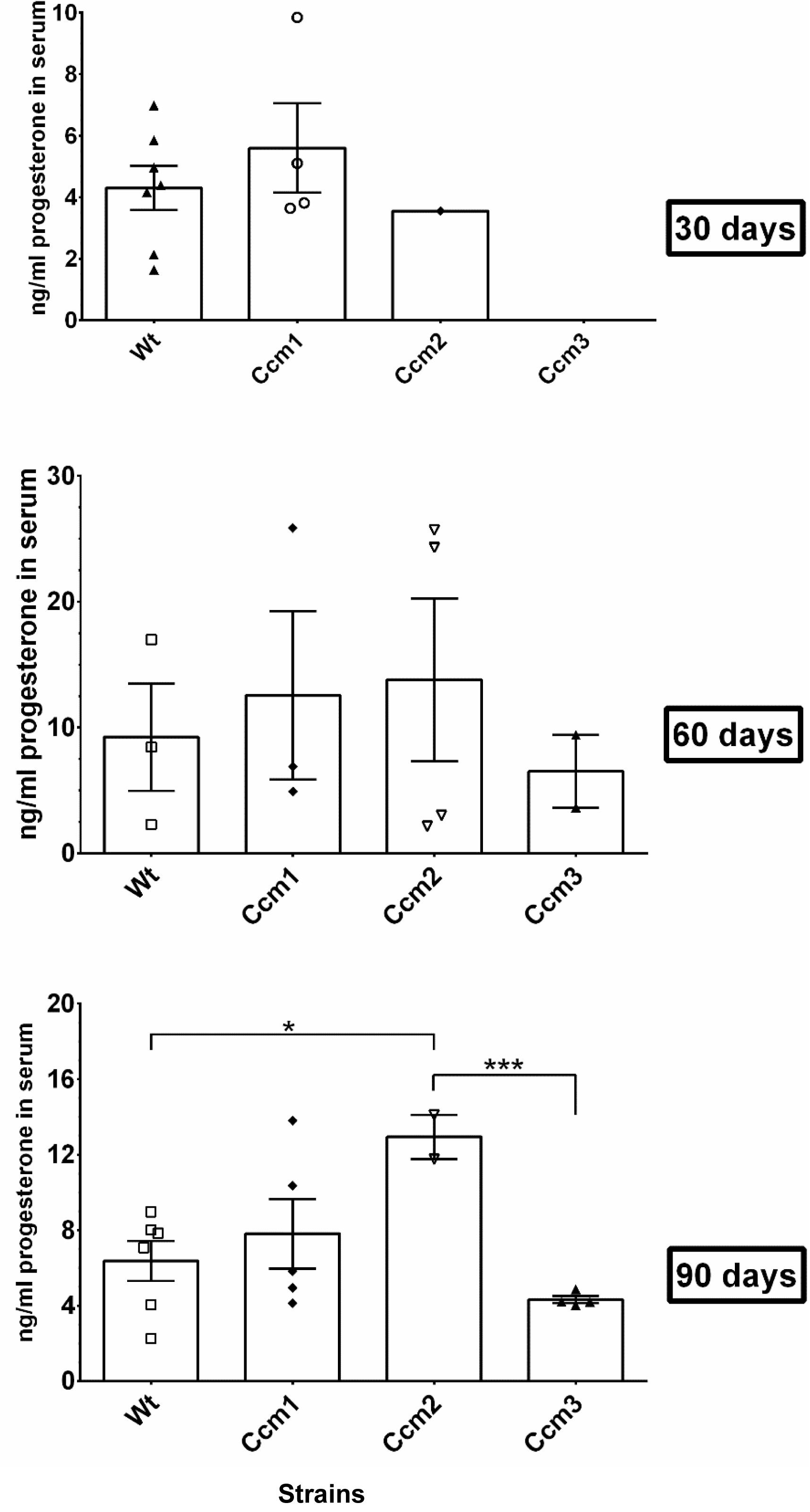
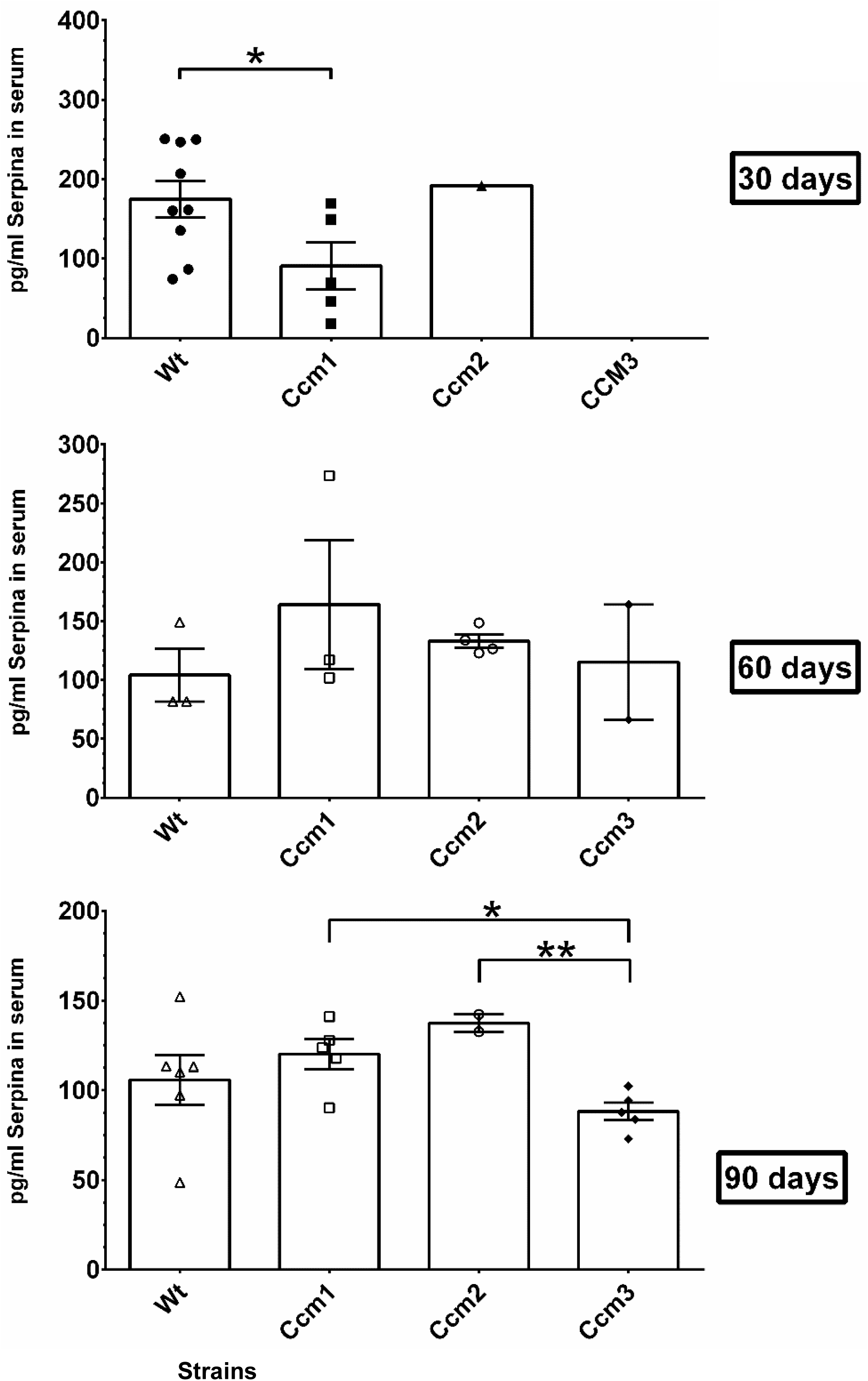
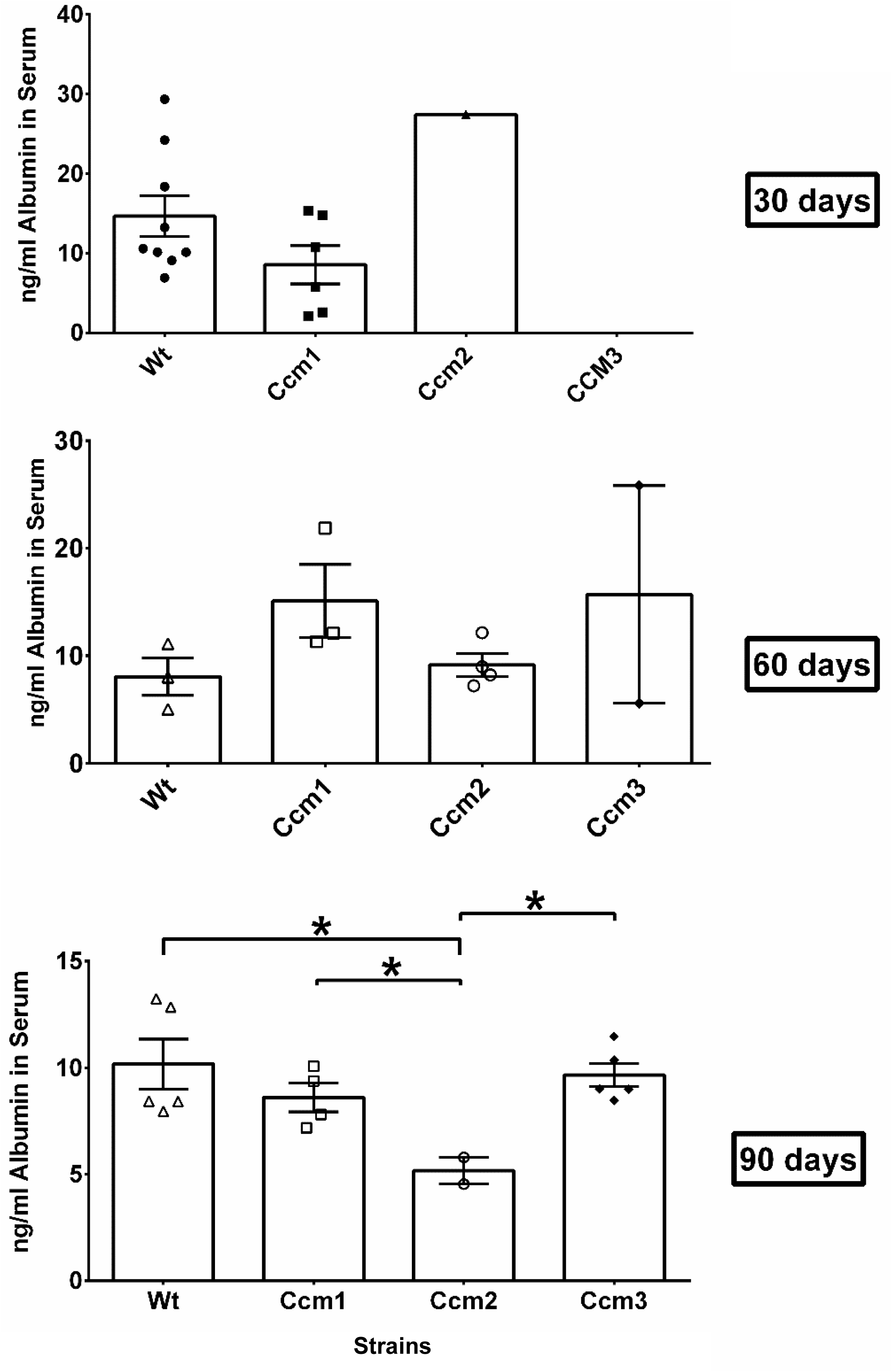
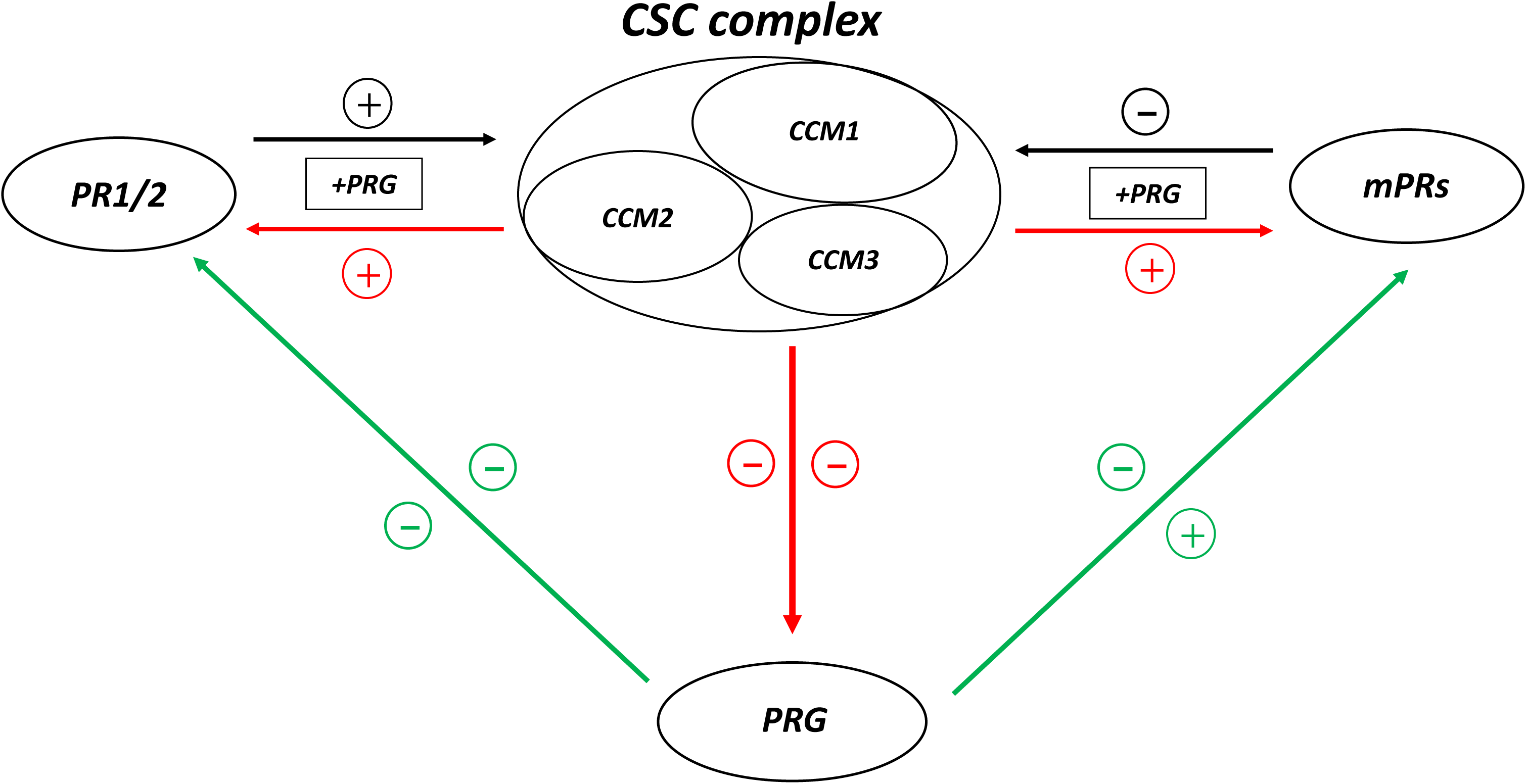
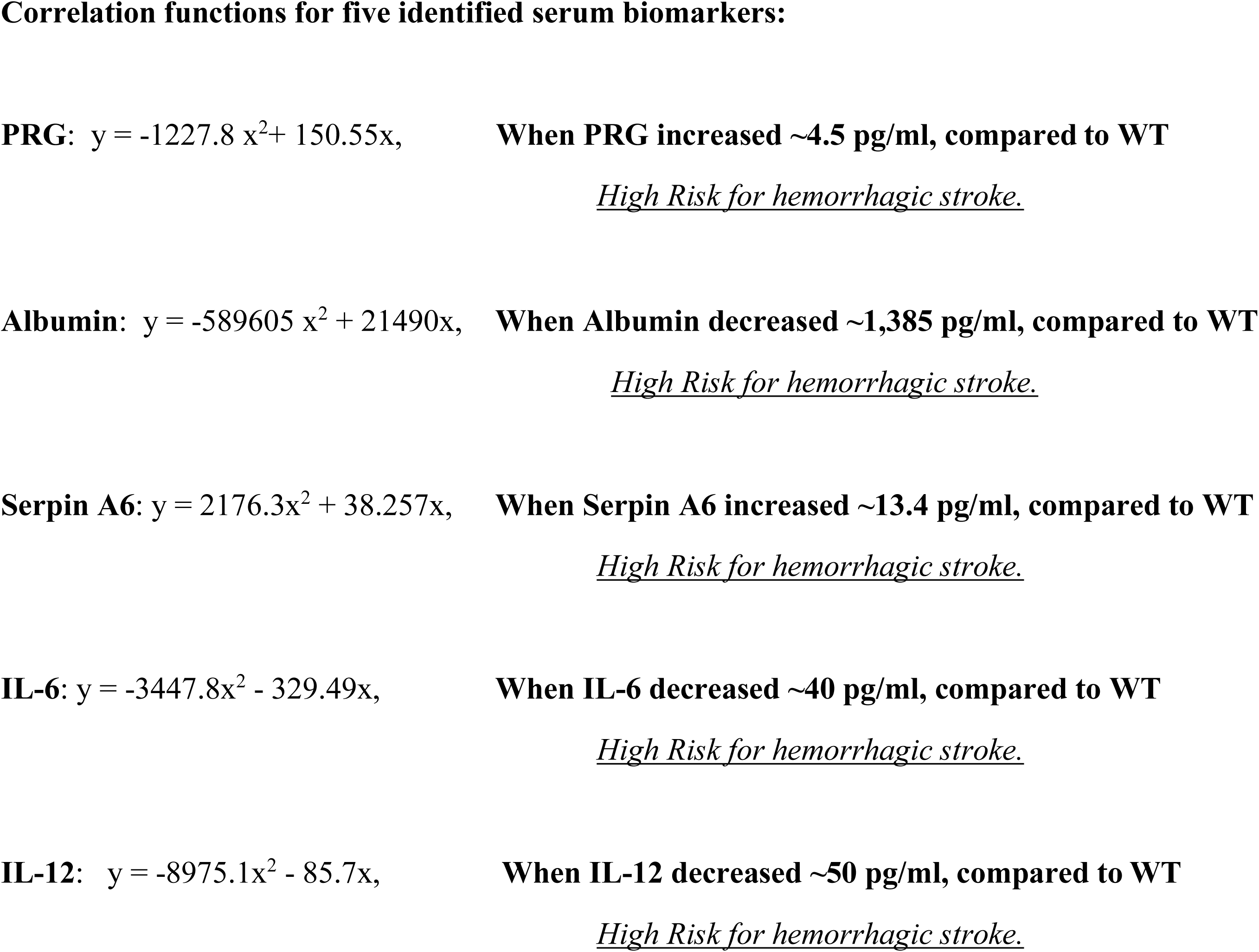
Perturbation of homeostasis/biogenesis with PRG is associated with initial hemorrhagic events in the pathogenesis of CCMs. **A**. Significantly increased PRG levels in serum indicates perturbed homeostasis/biogenesis of PRG in Ccms mutant mice. No obvious differences of PRG levels in mouse serum was observed in 30-day treatment group, suggesting that intrinsic mechanisms of homeostasis and biogenesis of PRG can still regulate PRG to normal levels after 30-days of injecting combined steroid (PRG+MIF) hormones. However, amounts of PRG in the mouse serum have an overall two-fold increase among all treated genotypes in 60-day treatment groups. Especially, amounts of PRG in the Ccm1 and Ccm2 mutant mice are much higher (even though not statistically significant), suggesting possible contribution of the genotype to PRG accumulation in serum. In 90-day treatment groups, amounts of PRG in the mouse serum regressed to normal range, likely though intrinsic mechanism of homeostasis and biogenesis of PRG in WT mice, however, amounts of PRG in the Ccm2 mutant mice stayed significantly higher (similar higher trend in Ccm1 mutant mice as well), indicating that intrinsic mechanisms of homeostasis of PRG was perturbed in Ccm2 (and possibly Ccm1) mutant mice. **B**. Increased quantity of serpin A6 suggest homeostasis of PRG with serpin A6 is functional. Significantly decreased serpin A6 levels were observed in Ccm1 mutant mice in 30-day treatment group only, while increased amounts of serpin A6 were observed in Ccm (1, 2) mutants in both 60 and 90-day treatment groups (not significant yet), suggesting that increased serpin A6 is functionally responding to dramatically increased PRG in serum. **C**. Decreased quantity of albumin suggest homeostasis of PRG through albumin is disrupted. Although albumin levels fluctuated in Ccms mutants in 30-day and 60-day treatment groups, similar to serpin A6 (mostly increased except CCM1 in 30 day group), significantly decreased amounts of albumin were found in Ccm2 mutant in 90-day treatment group (both Ccm1 and Ccm3 also showed decreased levels) compared to WT, suggesting that homeostasis of albumin is perturbed, leading to increased levels of active “free form” PRG in serum. **D**. CCM signaling complex (CSC) as a master regulator of homeostasis of progesterone is summarized in a simplified schematic representation, where key feedback receptors. Green color indicates effect from PRG onto PRG receptors. Black color indicates effect of PRG receptors onto the CSC. + Symbol indicates enhancement, while – symbol demonstrates an inhibitory effect. **E**. Serum levels of certain molecules (PRG, Albumin, Serpin A6, IL-6, and IL-12) were found to correlate with vascular permeability in BBB (leakage), which can be utilized as biomarkers to predict hemorrhagic strokes. The integrated evaluation of these markers will be the key to detect and prevent hemorrhagic stroke. Binomial regression of these five biomarkers in the serum with EBD in the brain was performed.

The amounts of PRG in serum were found to be elevated significantly in Ccm2 mutant mice in the 90-day group compared to WT, and the same trend of increased amounts of PRG in serum were also found in Ccm1 mutant mice but without significance (Fig. 7A, bottom panel), indicating an abrupt disruption of homeostasis of PRG is correlated with initiation of hemorrhagic events (Fig. 5A, bottom left panel) among Ccm2 mutant mice (and possibly Ccm1 mice) after long exposure to sex steroid hormones.

### Perturbation of albumin homeostasis/biogenesis is caused by a disrupted CSC

#### Changed expression levels of PRG-binding proteins in serum of Ccm2 mutant mice in 90-day treatment groups

Significantly decreased levels of serpin A6 in serum was found in Ccm1 mutant mice, compared to WT mice, in the 30-day treatment group. However, an increased trend of serpin A6 in serum was observed in both Ccm1 and Ccm2 mutant mice, compared to WT mice, in the 60 and 90-day treatment groups (although not statistically significant when compared to WT) (Fig. 7B), suggesting that PRG-binding protein, serpin A6, follow the same trend of serum level of PRG. These data suggest that the serum level and function of serpin A6 is normal, irrespective of genotype, and serpin A6 is upregulated to decrease active PRG levels in the serum.

Intriguingly, significantly decreased levels of albumin in serum was found in Ccm-2 mutant mice in the 90-day treatment groups despite no differences in either 30 or 60-day treatment groups (Fig. 7C), suggesting reduced level of albumin in serum might contribute to the perturbed homeostasis/biogenesis of PRG (Compare Fig. 7C and 7A) in Ccm2 mutant mice. These data also suggest that the CSC is highly likely involved in regulation of homeostasis of PRG by modulating its dominant binding protein, albumin, in serum. There is a similar trend between serum levels of albumin and serpin A6 in Ccm1, where both binding proteins are decreased in the 30-day treatment group (significant in serpin A6, non-significant in albumin), but subsequently are increased in 60-day treatment group (non-significant in both). This is correlated to immunosuppression shown previously (Figs. 6B-E), further validating the notion that the CSC modulates biogenesis of albumin, a dominant serum PRG binding protein (over 80%).

Our data indicate that the CSC not only plays a major role in modulating signaling crosstalk between classic and non-classic PRG receptors (*19*), but also influences the homeostasis/biogenesis of PRG, and this intricate relationship is illustrated in Fig. 7D.

#### Biomarkers associated with hemorrhagic strokes defined in Ccms mutant mice

Our data clearly demonstrated that there is a correlation between serum levels of PRG and the initiation of hemorrhagic strokes in Ccms mice. Furthermore, four more biomarkers (Serpin A6, IL-12, IL-6, and Albumin) are all associated with PRG homeostasis/biogenesis in serum, and also correlate well with hemorrhagic events in Ccms mutant mice. Therefore, we hypothesize that these serum molecules can be used as biomarkers to predict the timing of the first hemorrhagic event in CCM pathology. We performed binomial regression with the serum concentration of these four biomarkers with the corresponding EBD levels for the same treatment groups to generate binomial correlation functions (Fig. 7E). Each biomarker has a unique cutoff value indicating elevated risk of hemorrhagic event. These predictive algorithms will be tested through our ongoing collaborative human trials.

## Discussion

### Perturbed homeostasis of PRG is the trigger for hemorrhagic bleeding

#### Female gender is a key risk factor for hemorrhagic bleeding in CCMs

Our key findings is that PR(-) ECs are more susceptible to the perturbation of the CSC-PRG receptor signaling cascades, which is well documented in clinical appearance. CCMs are more common in women and become symptomatic during their reproductive period (30s-40s age range) (*58, 59*). Although no conclusive results have been found (*60*), hormonal changes during pregnancy have long been suggested as significant factors for the increased bleeding (*61–63*), and female gender is a key risk factor for bleeding in CCM patients (*61, 64*). The size of CCM lesions (*65–68*) and more cases of the hemorrhagic CCMs during pregnancy have been documented (*62, 69–74*), suggesting that pregnancy is associated with an increased risk of hemorrhage. Increased risk for acute CCM bleedings (*60, 62, 63, 72, 74, 75*), or formation of a de novo CCM lesion (*72, 75, 76*) have previously been reported during pregnancy.

Spinal hemangiomas have been defined as a pregnancy-related vascular disorder (*77–80*), and CCMs account for half of them (*81, 82*). Hormonal changes during pregnancy are defined as determining factors in initiating and enhancing the vertebral hemangiomas (*78, 83*). Therefore, it has been long speculated that the flux of hormones during pregnancy may predispose CCMs to hemorrhage (*61, 63, 70, 72, 73*).

#### PRG is a key risk factor for hemorrhagic CCMs

Significantly increased PRG levels during early pregnancy (*84, 85*) has been indicated to enhance the progression of vertebral hemangiomas (*86*), possibly through its ability to induce structural changes within the vessel wall (*87*). Negative staining with EST but positive staining for PRG in all 12 orbital CCM samples, suggested the role of PRG signaling in the pathogenesis of orbital CCMs (*88*). Unfortunately, screening results of CCM lesions from 12 patients showed all negative staining with both EST receptor (ER) and classic PRG receptor (PR1/2), failing to link both ER and PR hormonal signaling pathways to the pathogenesis of CCMs (*89*). Data in PR(-) blood monocytes and T cells (*46*) and PR(+) T47D cells (*19*) indicate that mPRs, is quite possibly, the only target for PRG actions in PR(-) cells, such as microvascular ECs.

#### New paradigm for hemorrhagic progression events in CCMs

In clinical diagnosis, hemorrhage is often rooted from defective endothelial cell junctions, and microvessel rupture is a result of compromised integrity of BBB in CCMs (*90*). Many bleeding mechanisms have been proposed, two major theories are anticoagulant vascular domain theory and gut microbiota theory. In the anticoagulant vascular domain theory, local increases in the endothelial cofactors that generate anticoagulant APC (activated protein C) could contribute to recurrent bleeding in CCM lesions (*91*). In the microbiota theory, gram-negative bacterial signaling through the lipopolysaccharide (LPS)-activated immune receptor, Toll-like receptor 4 (TLR4), promotes hemorrhagic bleeding in both Ccm1 and Ccm2 mutant mice, emphasizing the important roles for the gut microbiome and innate immune signaling in the pathogenesis of CCMs (*34*). This microbiota-gut-brain axis theory recently focuses on the importance of gut microbiota in influencing the interaction direction through inducing inflammatory gut milieu, leads to systemic inflamed milieu that might exacerbate the inflammatory response in the brain, and promote detrimental effects on BBB (*92*). However, LPS-induced Ccm hemorrhagic mouse models induce a large CCM burden with massive bleeding which contributes to lethality at the early stages of life, uncharacteristic of human CCMs. Milder bleeding Ccm mouse models have been generated in an effort to resolve these discrepancies (*93, 94*). Furthermore, it is well known that immunogenic LPS will compromise vascular function and integrity in general (*95*) including BBB (*96*), and neuroinflammation (*97, 98*), irrespective of either genotype or inflammatory response.

Nonetheless, neither of previous theories could address a key issue of gender discrepancy in the pathogenesis of CCMs, demanding further evaluation for proposed causes of hemorrhagic stroke. Although it is still under debate (*72, 99, 100*), female dominance in CCM patients have been long suggested (*72, 101, 102*), and consensus has been reached on the more severe bleeding with worse neurological outcomes in females (*61, 102*). This relatively aggressive course of hemorrhagic lesions in females has been proposed to be consequent to endocrine influences (*72, 101, 102*). Our data that enhanced PRG-mPRs signaling due to perturbed homeostasis of PRG leads to CCM hemorrhagic bleeding, in addition to evidence that long exposure to hormonal contraceptives increase risk of cerebral venous sinus thrombosis (CVST) (*103*), seems incongruent with the anticoagulant APC theory (*91*). Furthermore, our findings that immunosuppression caused by steroid actions in CCM deficient mice is associated with CCM hemorrhagic bleeding, also disagree with the microbiota theory (*34*). Therefore, this report proposes a new paradigm for the molecular mechanisms of initiating CCM hemorrhagic events. In PR(-) ECs, the feedback loops between the CSC, mPRs/PAQRs, and steroid actions appear to be quite sensitive and perturbation of this intricate balance, such as hormone therapy or hormonal contraception regimens could result in increased risks in early hemorrhagic events. Our new paradigm does not intend to rule out the possible involvement from the previously proposed theories, but instead provides a theory that is in line with the clinically observed human CCM conditions.

### Novel biomarkers shed light for the first time to predict the occurrence of hemorrhagic stroke

Biomarkers have been long sought for early diagnosis of strokes (*104–107*), but only with a limited success in ischemic strokes (*108–110*). Efforts to define diagnostic biomarkers for hemorrhagic strokes are underway (*111–114*). Inflammatory cytokines have been on target as risk factors or biomarkers for stroke (*115*). Circulating levels of TNF-α (*116–118*), MCP-1 (*119*), and IL-6 (*120–122*) as biomarkers have all been investigated. Among two cytokines from candidate plasma biomarkers in a small cohort (*113*), IL-1β demonstrated slightly elevated levels, while IL-6 plasma level was lower in hemorrhagic patients compared to stable subjects, which is in accordance with our serum IL-6 data in Ccms mutant mice.

Plasma levels of serpin A6 is nearly 20 fold higher from women relative to men, suggesting gender differences in response to PRG signaling (*123*). While no differences were found in women, a significant inverse association between plasma serpin A6 levels in men were detected, indicating a protective role of serpin A6 in strokes (*123*). Serum albumin levels have been found to correlate with stroke incidence and outcome (*124–130*), but the majority of this work only investigate ischemic stroke (*110, 131–136*) but found serum albumin levels can predict hemorrhagic stroke (*137, 138*) and is negatively associated between them (*138*). Since the protective role is so significant, high-dose albumin has been shown to be highly neuroprotective during strokes (*139–141*), and multi-center clinical trials have been carried out (*142–144*), but the neuroprotective function of albumin remains unknown and our data shed light on these possible mechanisms. Ironically, PRG has long been sought as a potential therapeutic drug for post-stroke treatment for both ischemic and hemorrhagic strokes in animal models (*145, 146*) and now there are even some on-going large clinical trials. The neuroprotective function of PRG has been well defined (*147, 148*), mainly due to its ability of immunosuppression (*149, 150*), therefore, it should be cautious to evaluate its potential application as treatment options when current data show the potential role of excessive PRG in inducing hemorrhagic events. Collectively, our findings reinforce the idea that there are gender and sex hormone-associated differences in stroke pathophysiology, and suggest that PRG-mediated signaling should be investigated further as a potential molecular mediator in strokes.

## Conclusion

In this report we extended our previous finding of the intricate feedback regulation between the CSC and PRG receptors mediated signaling into PR(-) cells. We found that the CSC controls the homeostasis of PRG, linking perturbed homeostasis of PRG to the center stage of hemorrhagic events of CCMs, in both *in-vitro* PR negative ECs and *in-vivo* mouse models which is further supported through our functional enrichment analysis of DEGs/DEPs from a disrupted CSC under steroid actions. This project provides new insights into cross-talk between PRG modulated signaling and CSC-mediated signaling pathways toward angiogenesis which helps maintain the integrity of the BBB, revolutionizing the current concepts of vascular malformations and molecular mechanisms of hemorrhagic occurrences, leading to new therapeutic strategies.

## Acknowledgments

We wish to thank Dr. Douglas Marchuk at Duke University for providing us heterozygous Ccms mutant strains of mouse model; Elias Gonzalez, Muaz Bhalli, Alexander Le, Khalid Shoukat, Deepak Muthyala, Mike Yao, Yanchun Qu, Shen Sheng, Ahmed Badr, Junli Zhang, Amna Siddiqui, P. Dubey, Saafan Malik, and Edna Lopez at Texas Tech University Health Science Center El Paso (TTUHSCEP) for their technical help during the experiments.

## Author contributions

JZ: Conceptualization, Methodology, Investigation, Writing-Original draft preparation, Writing-Reviewing and Editing; JAF: Investigation, Software, Data curation, Validation, Writing-Reviewing and Editing, XTJ: Investigation, Visualization; AP: Investigation, Writing-Original draft preparation, DG: Investigation, Data curation; DM: Investigation; MS: Investigation; BG: Software, Data curation, Validation; WW: Investigation, Data curation.

## Data and materials availability

All data is available in the main text or the supplementary materials.

## Supplemental information

### Abbreviations

CCM: Cerebral cavernous malformation
CSC: CCM signaling complex
PRG: Progesterone
MIF: mifepristone
PR1/2: progesterone receptors isoform A/B
mPRS: membrane progesterone receptors
PAQRs: progestin and adipoQ receptor

## Materials and Methods

### Cell Culture and Treatment, Real Time Quantitative PCR analysis (qPCR), and Western blots

Multiple primary endothelial (HUVEC, HDMVEC, HBMVEC, RBMVEC) and 293T cell lines were cultured following manufacturers recommendations (ATCC) and as previously described (*1–6*). Briefly, when cells reached 80% confluency, cells were treated with either vehicle control (ethanol/DMSO, VEH), mifepristone (MIF, 20 µM), progesterone (PRG, 20 µM), combined hormones (PRG+MIF; 20µM each), or media only (Untreated) respectively for steroid treatments. For RNA knockdown experiments, 80% confluent 293T and ECs were transfected with a set of siRNAs, targeting specific genes, by RNAiMAX (Life Technologies) as described before (*7–9*). For transient transfection, 293T cells were transfected with different CCM2 isoform-MYC constructs by Lipofectamine2000 (Life Technologies) when cells reached 75% confluency. CCM2 isoforms were inserted into pcDNA3.1/V5-His-TOPO and Topo pCR®II plasmid vectors (Invitrogen) respectively (*1-6, 10, 11*) Jiang, 2019 #1996; Abou-Fadel, 2020 #1482}. The expression of *CCM* genes at both the transcriptional and translational levels was confirmed through both Real-time quantitative PCR (qPCR) and Western blots as previously described (*7*) with detailed information provided (Suppl. Table 1). Briefly, after various treatments, cells were harvested, and RNA expression levels of *CCMs, PR1/2* and *mPRs* genes were determined through quantitative PCR (qPCR) using Power SYBR Green Master Mix with ViiA 7 Real-Time PCR System (Applied Biosystems), and data were analyzed with DataAssist (ABI) and Rest 2009 software (Qiagen). All experiments were performed with triplicates as described before (*7, 8*). The relative expression levels of candidate proteins were measured with Western blots (WB) as described before (*7–9*).

### Omic data collection and bioinformatics analysis of DEGs/DEPs to evaluate altered signaling cascades

All LC-MS/MS details, and bioinformatics processing of both RNAseq and Proteomics data was performed as described before (*12*). Briefly, For RNAseq workflow, we removed the reads mapped to rRNAs to get the raw data. We then filter the low quality reads (more than 20% of the bases qualities are lower than 10), with adaptors and reads with unknown bases (N bases more than 5%) to get the clean reads. We then assembled those clean reads into Unigenes, followed with Unigene functional annotation and calculate the Unigene expression levels and SNPs of each sample. Finally, we identify DEGs (differential expressed genes) between samples and do clustering analysis and functional annotations. Briefly, for Proteomics workflow, database searching was performed in Sequest with a fragment ion mass tolerance of 0.020 Da and a parent ion tolerance of 10.0 PPM. Carbamidomethyl of cysteine was specified in Sequest as a fixed modification. Oxidation of methionine and acetyl of the n-terminus were specified in Sequest as variable modifications. For protein identification, Scaffold (version Scaffold_4.8.7, Proteome Software Inc., Portland, OR) was used to validate MS/MS based peptide and protein identifications. Peptide identifications were accepted if they could be established at greater than 95.0% probability by the Peptide Prophet algorithm (*13*) with Scaffold delta-mass correction. Protein identifications were accepted if they could be established at greater than 99.0% probability and contained at least 1 identified peptide. Protein probabilities were assigned by the Protein Prophet algorithm (*14*). Proteins that contained similar peptides and could not be differentiated based on MS/MS analysis alone were grouped to satisfy the principles of parsimony. Proteins sharing significant peptide evidence were grouped into clusters. Proteins were annotated with GO terms from NCBI (downloaded Oct 25, 2018) (*15*).

### Evaluation of Blood Brain Barrier with Evan’s Blue Dye from peripheral/central organs

#### Control and treatment groups

Ccms (1, 2, 3) mutants and WT (C57BL/6J) mice were injected with a combined steroid treatment, a cocktail of progesterone + mifepristone (100 mg/kg body weight) in peanut oil (vehicle), 5 days a week for 30, 60, and 90 days, respectively. Mice were also injected with only peanut oil or left untreated as vehicle and naïve controls, respectively. Upon completion of the last injection, mice were injected intravenously with EBD (500µG/25G body weight) and allowed to circulate for 3 hours followed by euthanasia and transcardial perfusion with sterile ice-cold 1X PBS; organs were harvested and snap frozen in liquid nitrogen for future use. Individual points on graph represent a mouse, and sample size, depending on genotype and treatment group normally ranged from N=3-20.

#### Extraction of Evan’s Blue Dye (EBD) from peripheral/central organs

Tissues were homogenized in 1.5mL tubes using pre-sterilized pestles in 600 µl of ice cold PBS. All samples were then loaded on a bead beater machine and agitated at max speed for 1 minute and quickly placed back on ice. After centrifugation at 15,000 RPM at 4°C, supernatant was transferred to a new tube with 300 µl of 100% TCA for a final volume of 900 µl. Samples were rocked overnight at 4°C and again centrifuged at 15,000 RPM at 4°C. 600 µl of supernatant was then transferred to a new tube with 300 µl of ethanol before measuring. The 900 µl of each sample is loaded in 8 wells of a 384 well black Costar plate to serve as technical replicates. The plate was then analyzed using a flex station 3 with fluorescence being read using 620nm excitation and 680 emission, with 6 readings per well, auto PMT sensitivity, and column wavelength priority set during the reads.

### Ear Collection and Preparation

Previous described procedure (*16*) was adapted and optimized. Whole ears were collected from WT (C57BL/6J) and Ccm (1, 2, 3) mutant mice after being treated for 90 days with MIF+PRG in carrier oil (peanut oil), peanut oil as a vehicle control or untreated as a naïve control. Ears were kept frozen at −80°C and thawed at 4°C before preparation. Ears were placed in Hanks Balanced Salt Solution (HBSS) so that hairs and excess tissue could be removed with scissors under a dissection microscope. Posterior and Anterior leaflets of the ears were separated with each leaflet being placed in a separate 24 well plate with 1 mL of 4% paraformaldehyde at 4°C for 15 min. Ears were washed with 1 mL PBS with 0.2% triton X-100 (Washing Buffer), then remaining cartilage and hair was removed and washed again before proceeding to staining procedures.

### Immunofluorescence (IF) Staining

Previous described procedure (*16*) was adapted and optimized. Briefly, prepared ears were placed in 24 well plates (all wash/incubation steps performed in separate wells) with 0.05% (*w/v*) Pronase (antigen retrieval) in PBS with 0.2% triton X-100, and incubated with gentle agitation at room temperature for 30 min. Ears were washed with wash buffer (1X PBST), followed by blocking buffer (10%BSA+0.2% triton X-100 (TX-100)), incubated at room temperature for 30 min, followed with wash buffer. Ears were then incubated with PECAM-1 conjugated antibody (FITC, 488nm) diluted in blocking buffer to a 1/50 concentration. The plate was covered with aluminum foil and incubated for 1 hour at room temperature. Rhodamine Phallodin conjugated antibody (TRITC, 565nm) and DAPI (408nm) diluted to a 1/200 concentration for each was then added into the well without removing PECAM-1 solution and incubated overnight at 4°C in the dark.

### Preparation of Gelatin Slides

A gelatin coating solution was prepared by dissolving 5g of gelatin in 1 L of heated water; 0.5g of chromium potassium sulfate was then dissolved in the gelatin. The solution was filtered and applied to slides in a histological dipping tank. Briefly, cleaned slides were placed in a rack and dipped in the gelatin coating solution 3 to 5 times (5 sec. each). The slides were allowed to dry at room temperature for 48 hrs and then stored at 4°C until ready to use.

### Mounting on Slides and Imaging

Stained ear leaflets were placed in a petri dish with PBS to prevent drying out during this procedure. Any remaining hairs, cartilage tissue, dust and fibers remaining after IF staining were removed from the ears under a dissection microscope. The ear leaflets were transferred to a gelatin slide with the inside of the leaflet facing up. Prolong Gold was placed on the ear leaflets to mount. Ears were clamped and placed under a box covered in aluminum to dry. After drying the slides were sealed with nail polish and allowed to dry.

The slides were imaged using a Nikon Eclipse Ti confocal microscope with the appropriate lasers for the antibodies used (Suppl. Table 1). 10-20 random images (anterior and posterior) from 3 mice from each strain in the 90 day treatment groups (spanning the full surface area of the ear) were taken for a representative sample of the ears depending on the size of the ear using z-stacks. Counts of lesions present for the treatment groups, along with the CCM2 naïve and vehicle controls were also obtained during this time. The images were saved as an 8-bit multicolor, black and white (required for use in the ImageJ software with vessel analysis package). Briefly, images were first inverted then converted to a bit map then analyzed for Vessel Diameter, Vessel Length Density, and Vascular Density. Statistical significance was determined using unpaired t-test.

### Measurement of cytokines and serum biomarkers

#### Estimation of monocytes and neutrophils in the peritoneal lavage

Peritoneal lavage was collected from Ccms (1, 2, 3) mutants and WT mice as previously described (*17*). Briefly, after euthanasia, 3mL PBS was injected into the peritoneal cavity of the mouse. The abdomen was gently massaged for 2min. Subsequently, lavage fluid was recovered using a pipette. Cells were counted on a hemocytometer slide and 1X10^6^ cells were resuspended in staining buffer (PBS with 2% FBS). Staining was carried out as described previously (*17*). Briefly, cells were incubated with CD16/CD32 antibody (Fc shield) for 5 min on ice, to prevent non-specific binding. Subsequently, cells were incubated for 60 minutes on ice with antibodies against cell surface markers (Suppl. Table 2). Cells were fixed for 5 minutes on ice with IC fixation buffer (eBiosciences, cat#00-8222-49). Labeled cells were analyzed on a BD FACSCantoII flow cytometer (BD Biosciences) and data was analyzed using FlowJo software (FlowJo). Sample size ranged from N=3-8, depending on strain and treatment group.

#### Measurement of cytokines and other biomarkers in mouse serum with ELISA assays

The serum levels of four serum cytokines, TNF-α, MCP-1, IL-12 and IL-6 as well as PRG and PRG binding proteins, were measured using corresponding ELISA kits (Suppl. Table 2) following manufacturer’s instructions.

## Supplementary Text

Supplemental Figures/tables are numbered to closely follow along with the numbering of figures in the main text, although there might be some inconsistent gaps in the numbering of the supplemental figures/tables. We have tried our best to aid in readability. There is no Supplement Figure 4, which we left empty to keep numbering consistent when referencing main text figures.

---

**Fig. S1.**
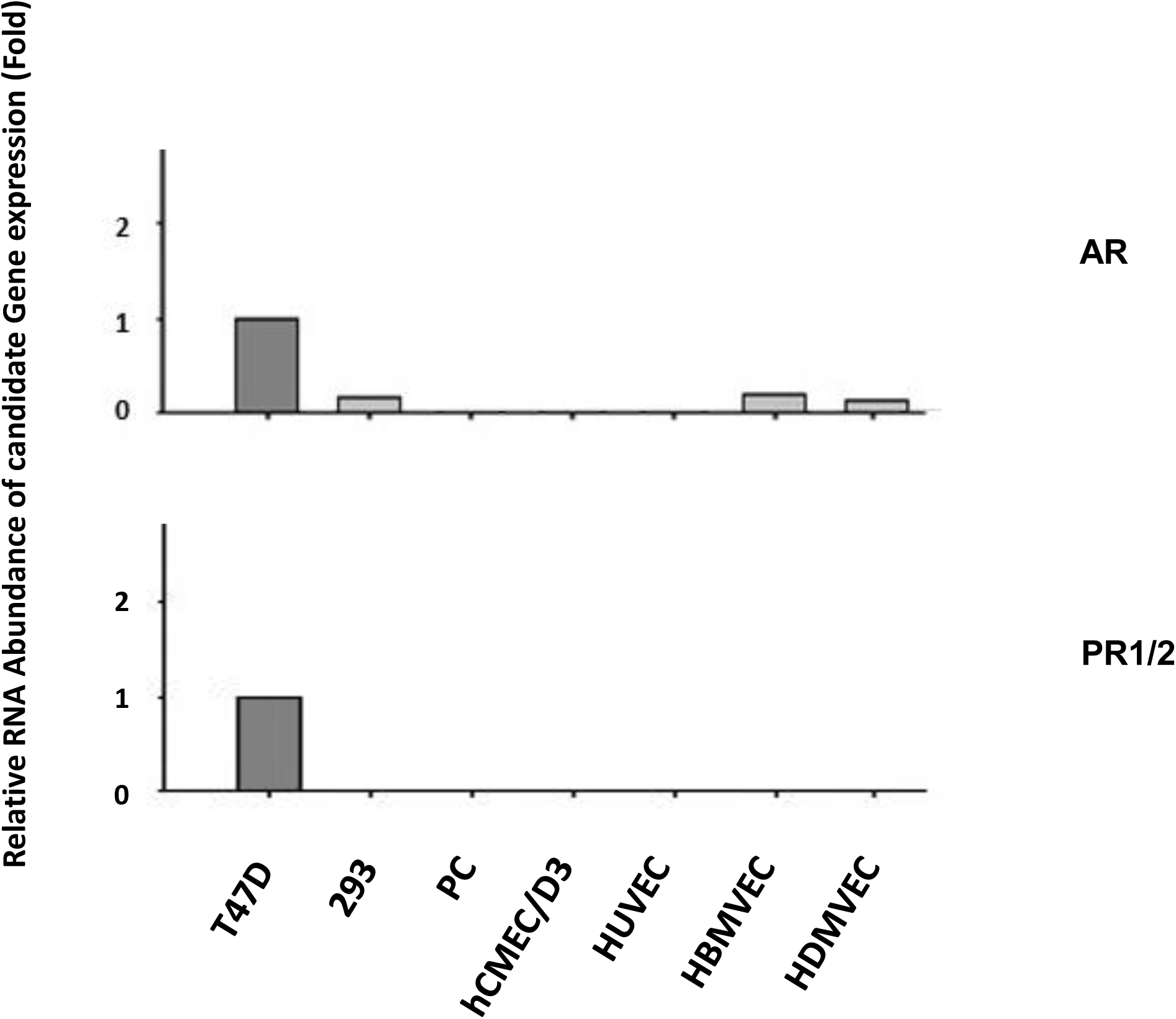
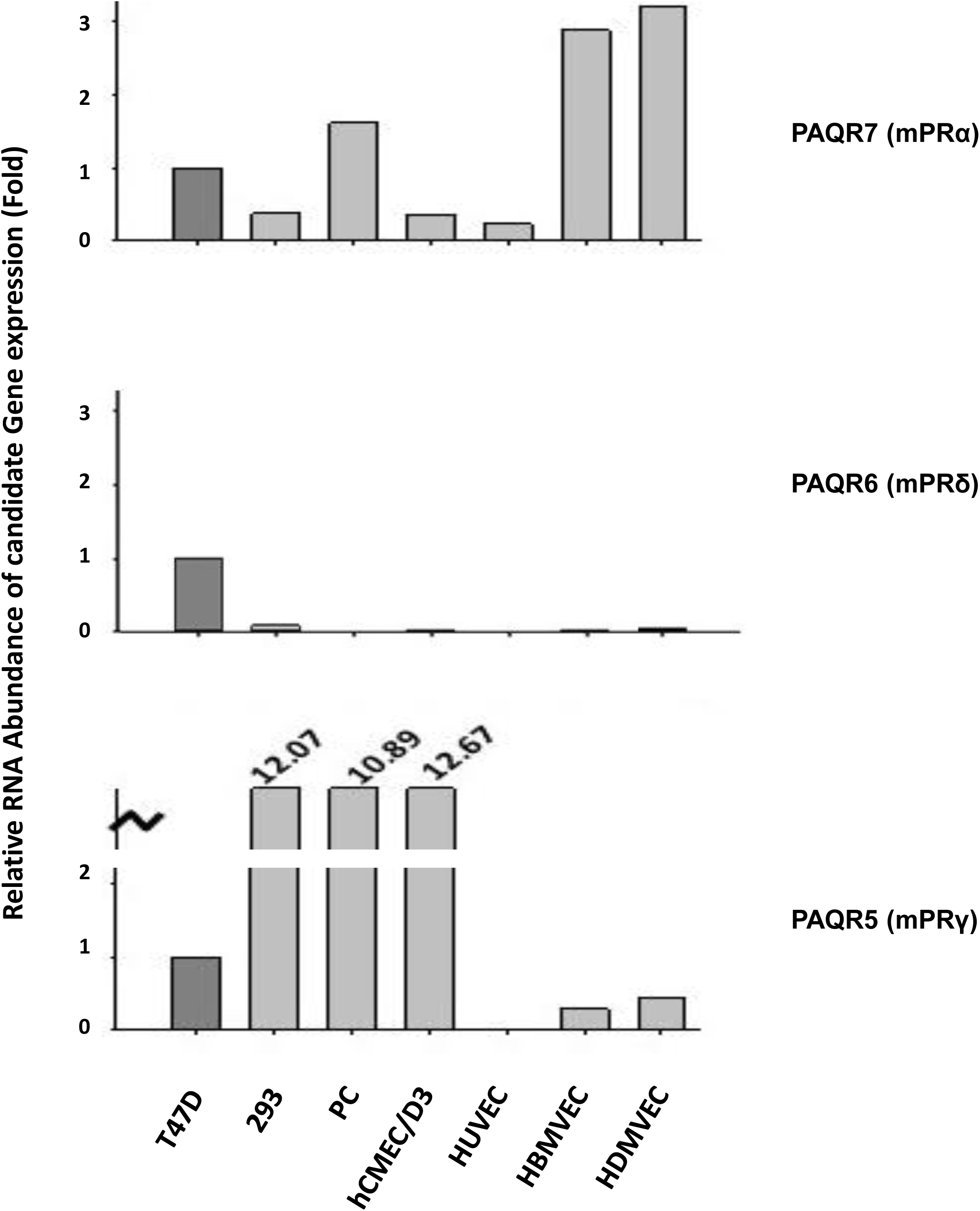
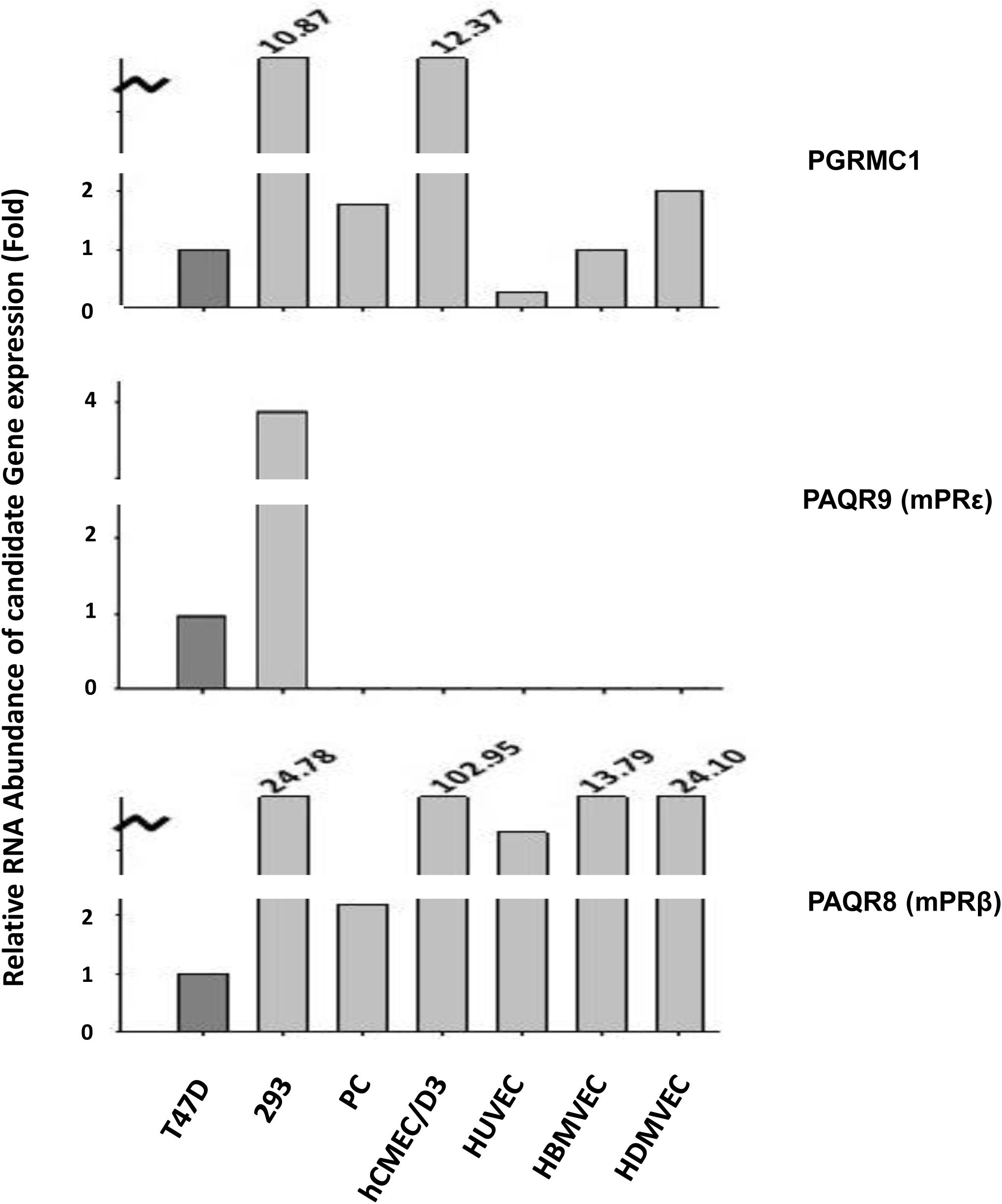
**Relative RNA expression profiling of endogenous classic and non-classic PRG receptors and related sex steroid receptors in PR(+) and PR**(-**) cell lines by qPCR.** The expression levels of AR and PR1/2 (**A**), PAQR5/6/7 (**B**), PAQR8/9 and PGRMC1 (**C**) are presented with bar plots, while β-actin was taken as an inference gene. For endothelial cell lines: HBMVECs, human brain microvascular endothelial cell; HDMVECs, human dermal microvascular endothelial cells; HUVEC, human umbilical vein endothelial cells; hCMEC/D3, immortalized human brain microvascular endothelial cells.

**Fig. S2.**
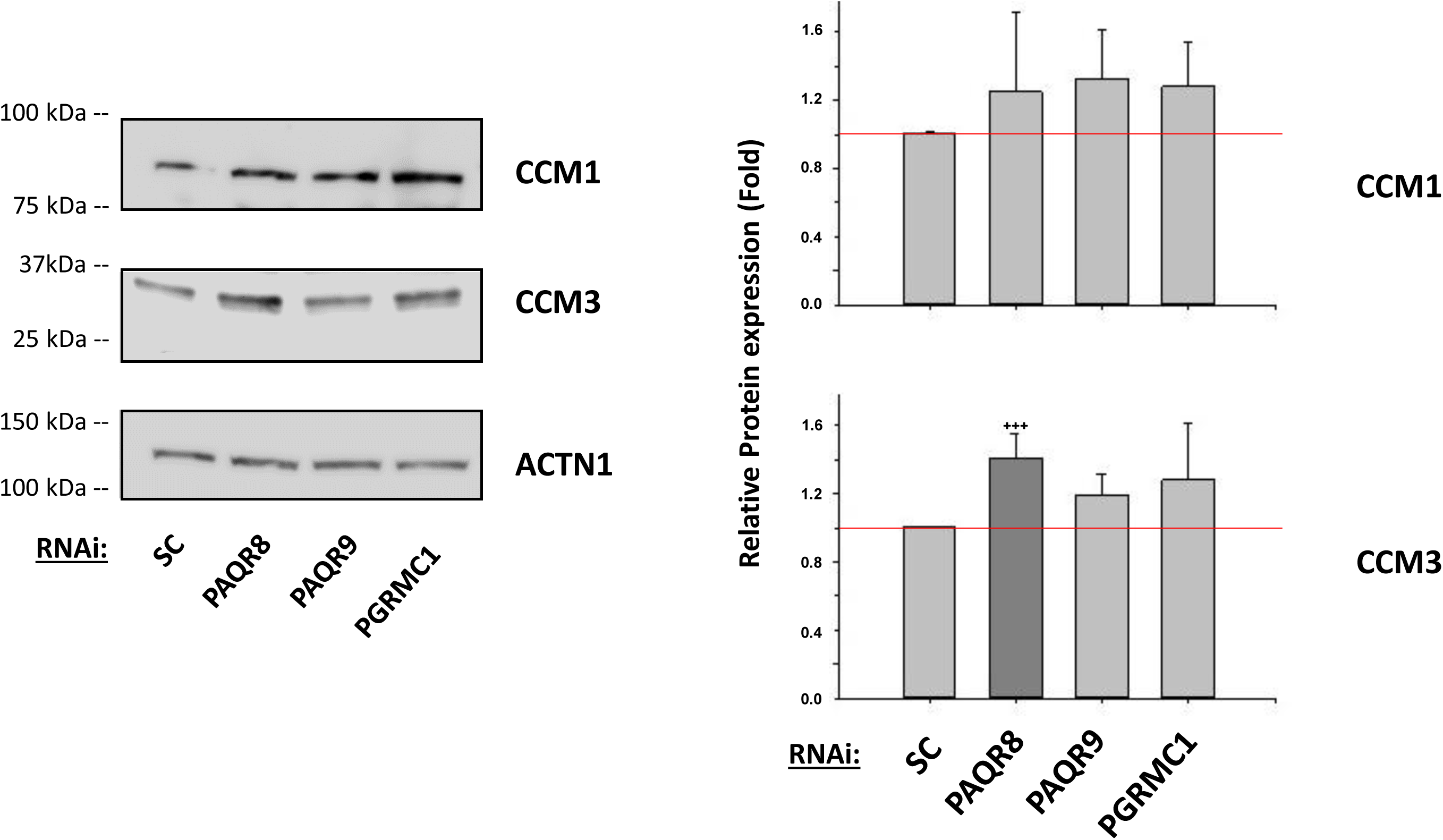
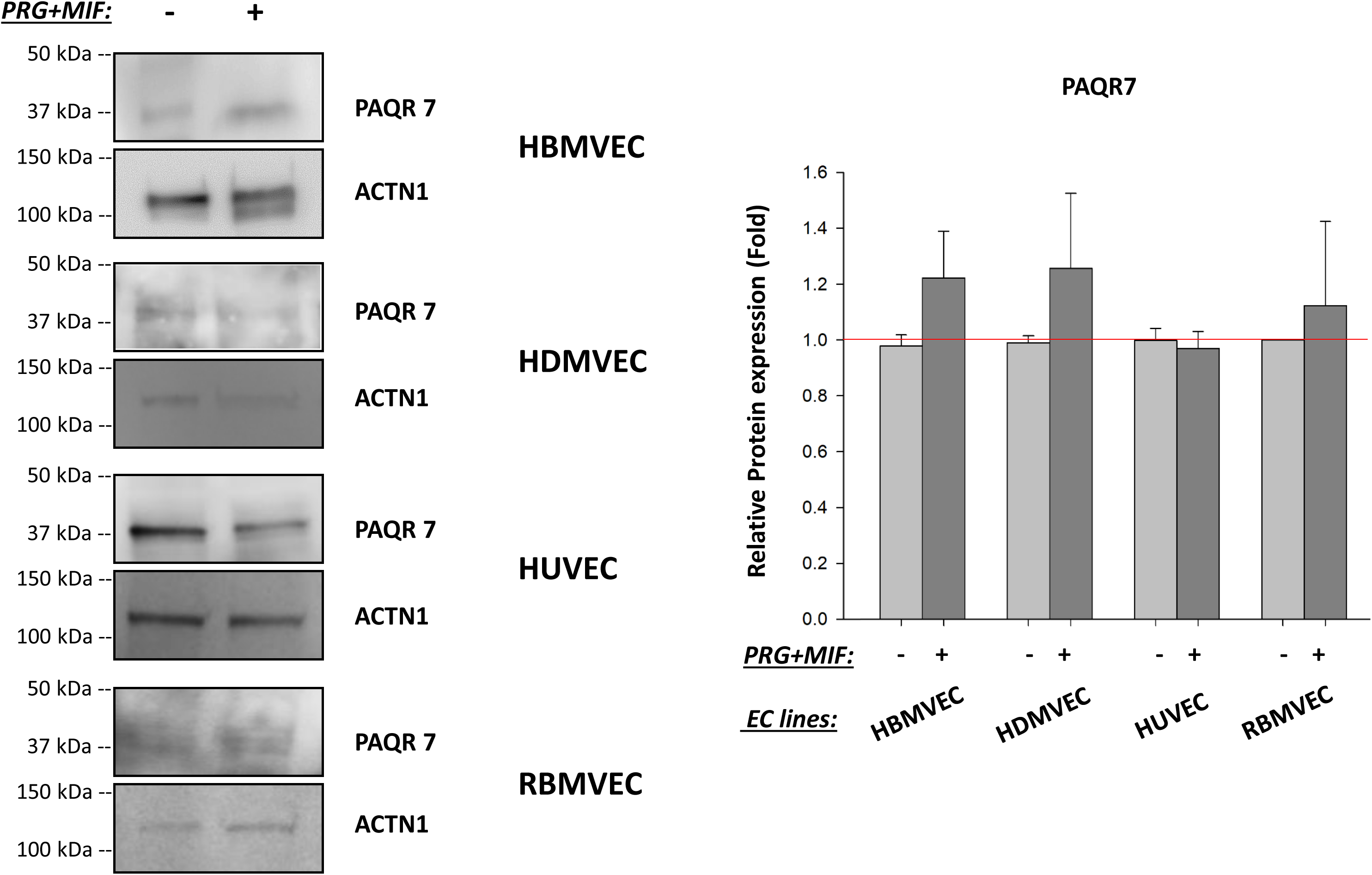
**Relationships among PR1/2, PAQRs, PRG/MIF, and the CSC in PR**(-**) 293T and endothelial cell (EC) lines. A.** Protein expression levels of both CCM 1/3 proteins are barely influenced by depletion of membrane progesterone receptors (PAQR 8, 9 and PGRMC1). After silencing mPRs genes (PAQR 8, 9, PGRMC1) for 48 hrs in 293T cells, the relative expression level of CCM1/3 proteins were barely changed; only increased CCM3 protein level in PAQR8-KD condition was observed (right panel). The relative expression level of CCM 1/3 proteins measured through quantification of band intensities of CCM proteins from three different experiments and normalized against α-actinin (ACTN1) followed by SC controls, and represented with bar plots where light grey bars represent for no change, dark grey bar for significantly increased protein level (n=3). **B.** The expression level of PAQR7 protein in PR(-) EC lines, HBMVEC (human brain microvascular endothelial cells), HDMVEC (human dermal microvascular endothelial cells), HUVEC (human umbilical vein endothelial cells), and RBMVEC (rat brain microvascular endothelial cells), was not influenced by steroid actions. After PRG+MIF treatment (20 µM each) for 48 hrs, no change of relative expression levels of PAQR7 protein was observed among PR(-) endothelial cell (EC) lines (left panel). The relative expression levels of PAQR7 protein were measured through quantification of band intensities of PAQR7 proteins and normalized against α-actinin (ACTN1) followed by vehicle controls (right panel). **C.** The relative RNA expression levels were measured through RT-qPCR from at least three different experiments (triplicates per experiment). The relative protein expression levels were measured through quantification of band intensities of targeted proteins, subtracted from the surrounding background and normalized against control housekeeping proteins followed by vehicle controls. In all bar plots, red line is the control baseline for fold change measurements (-/+). **, *** above bar indicate P ≤ 0.01 or 0.001 for paired t-test, respectively.

**Fig. S3:**
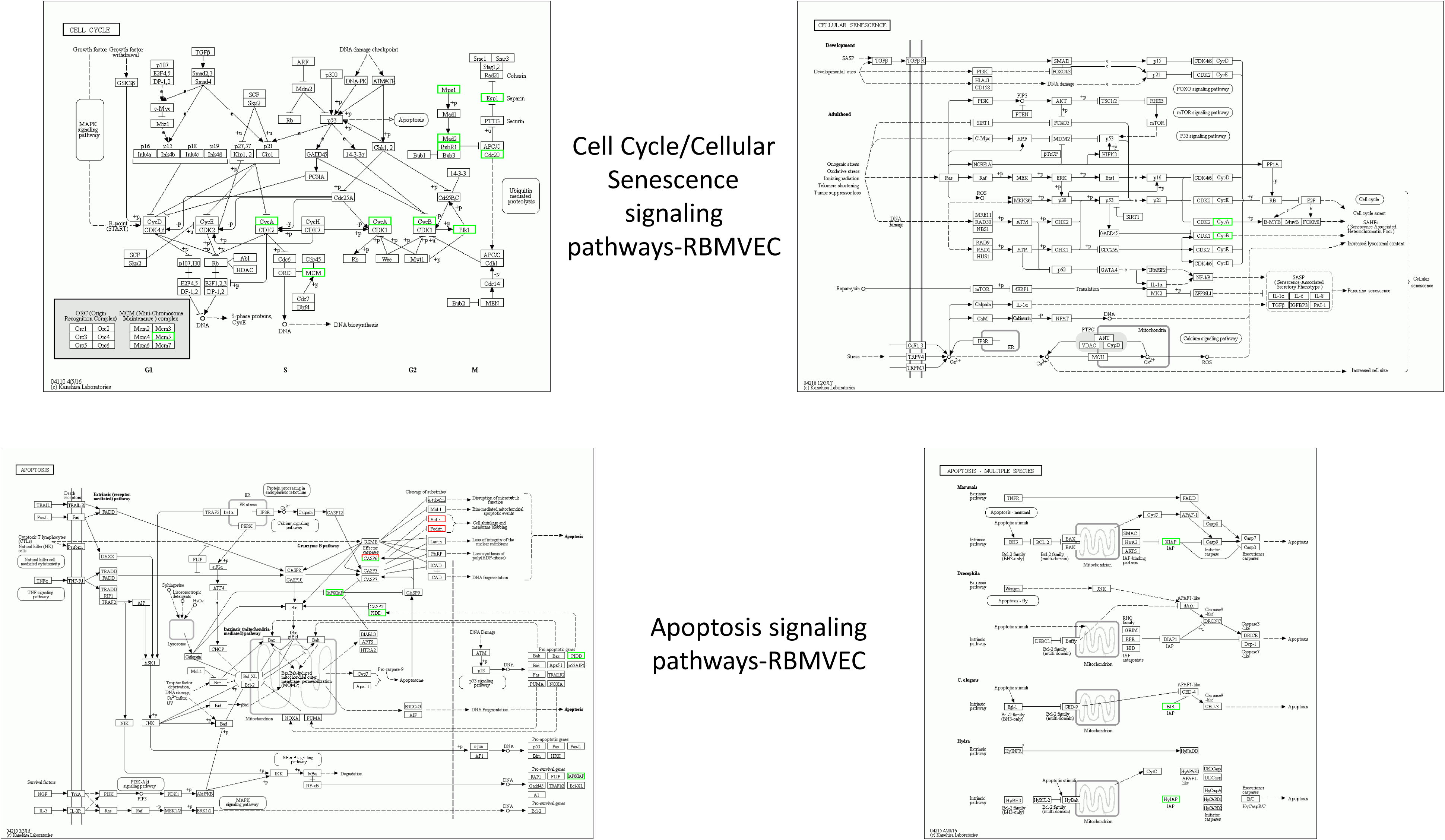
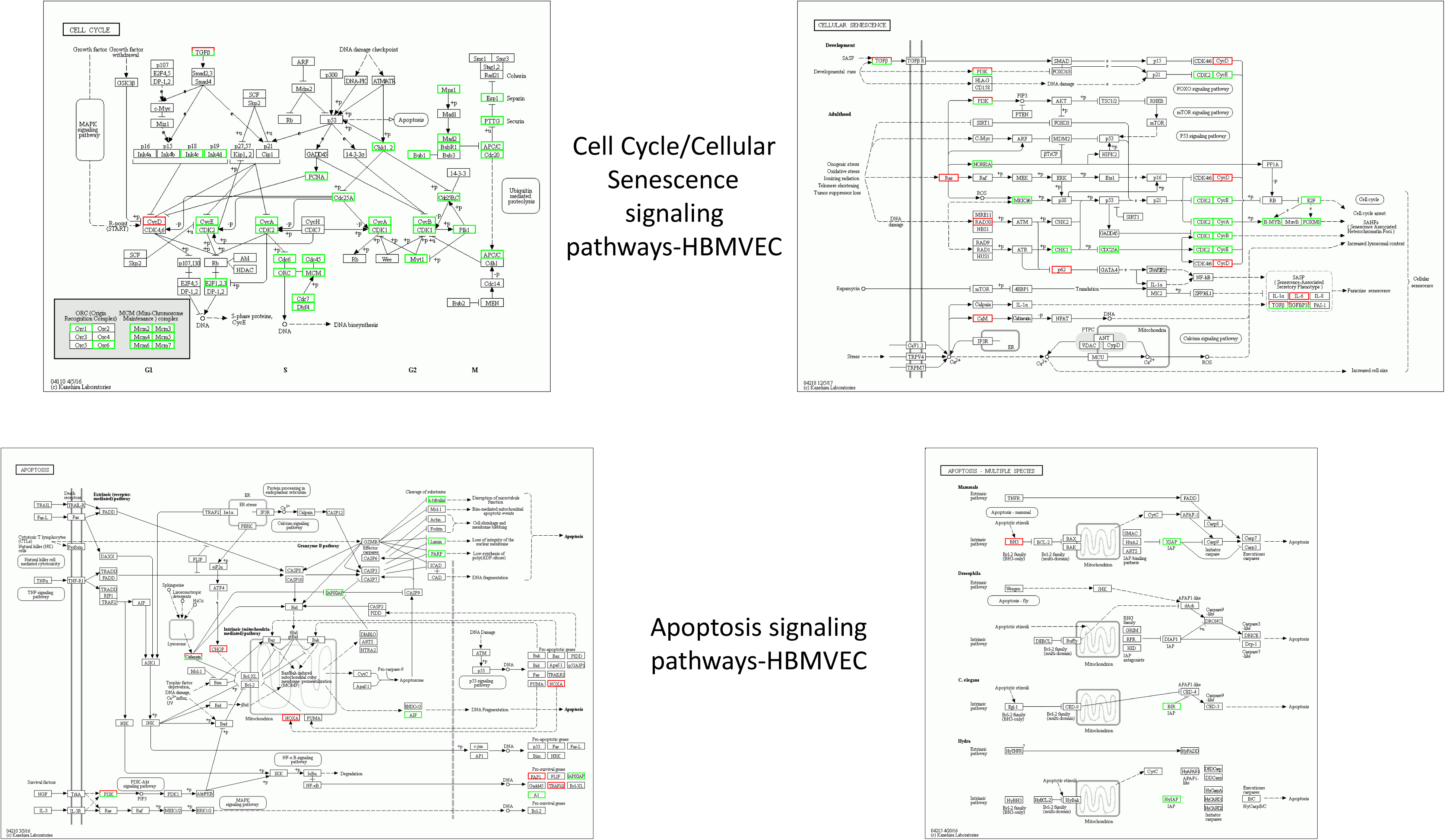
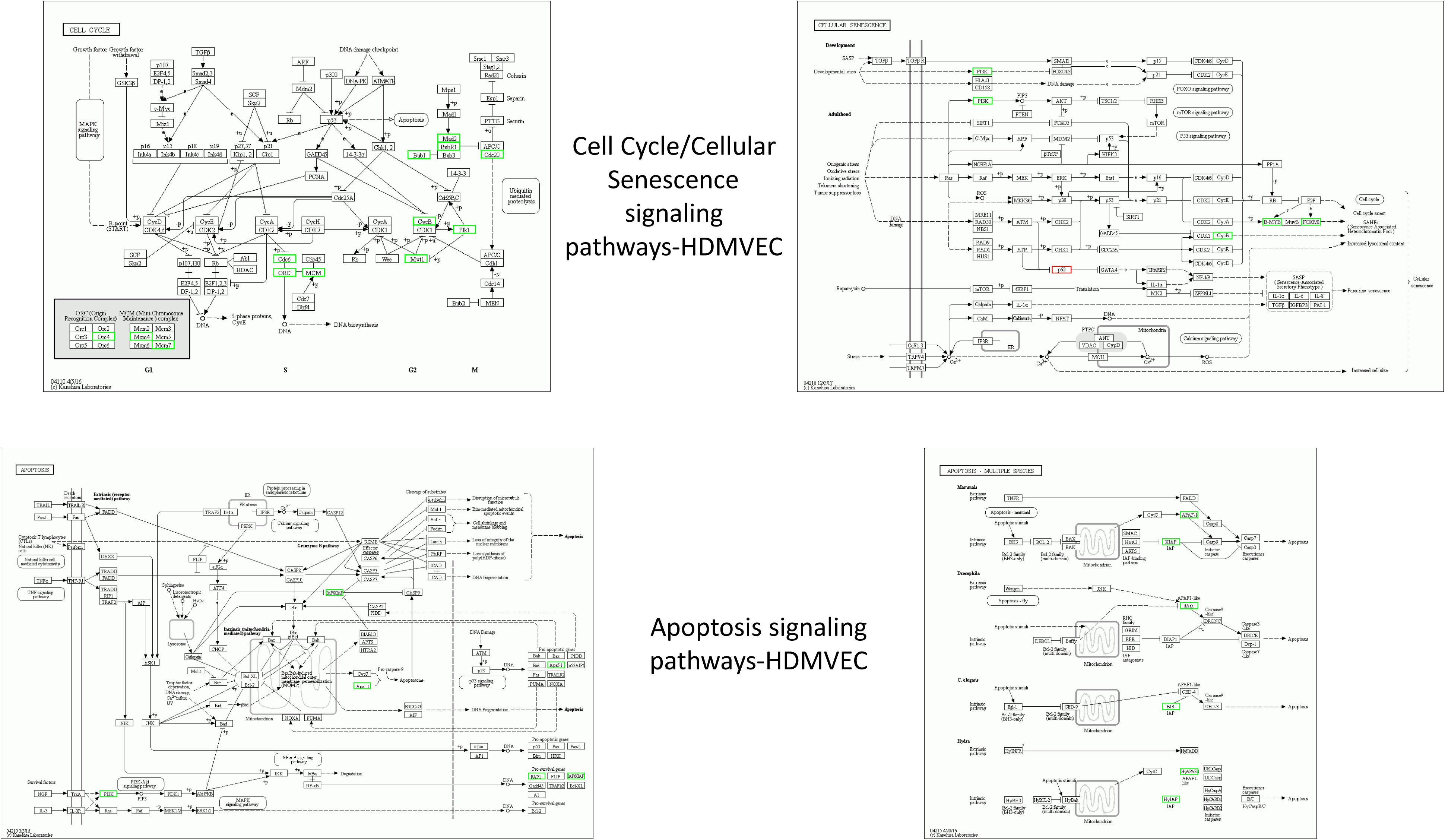
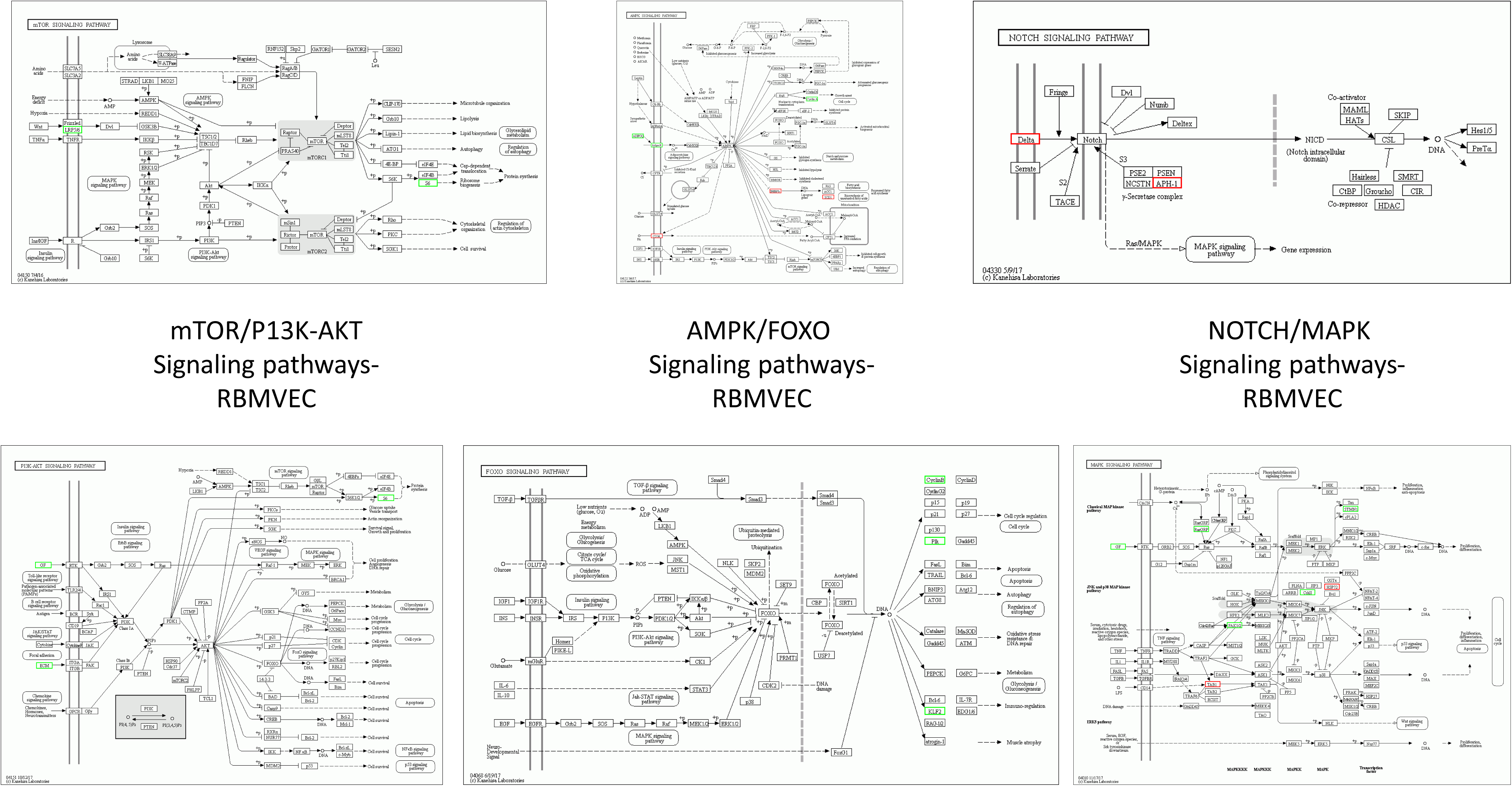
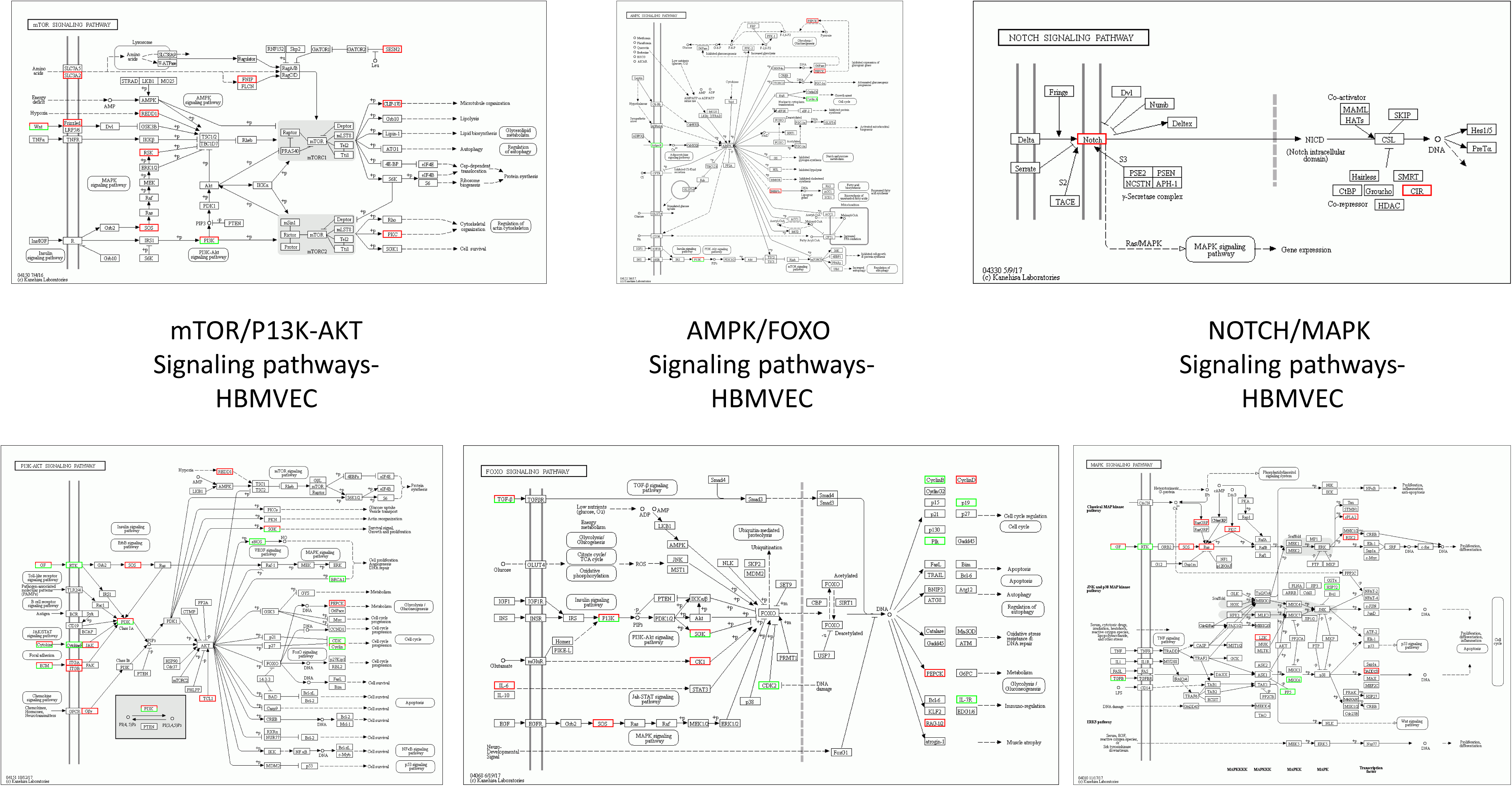
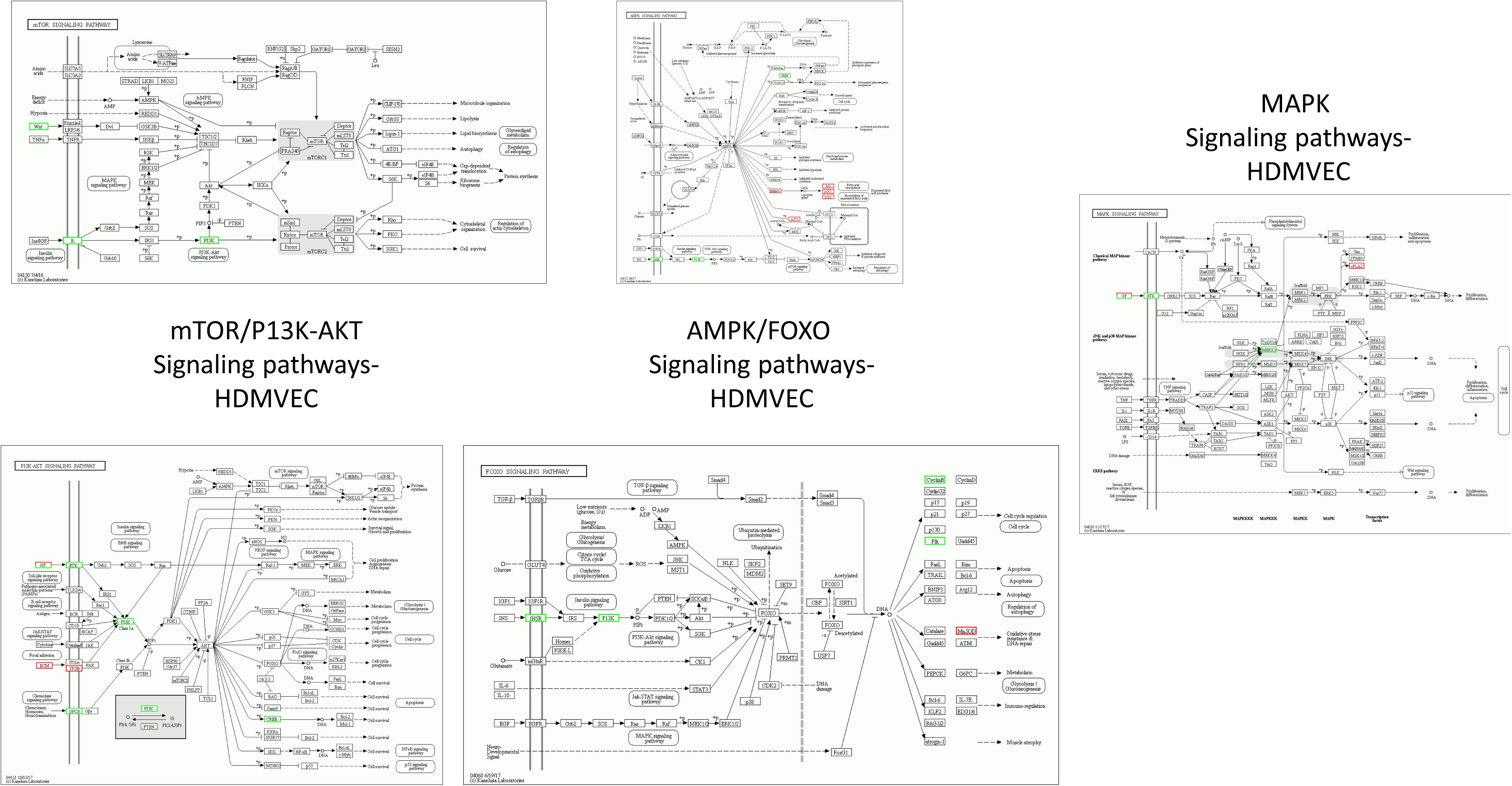
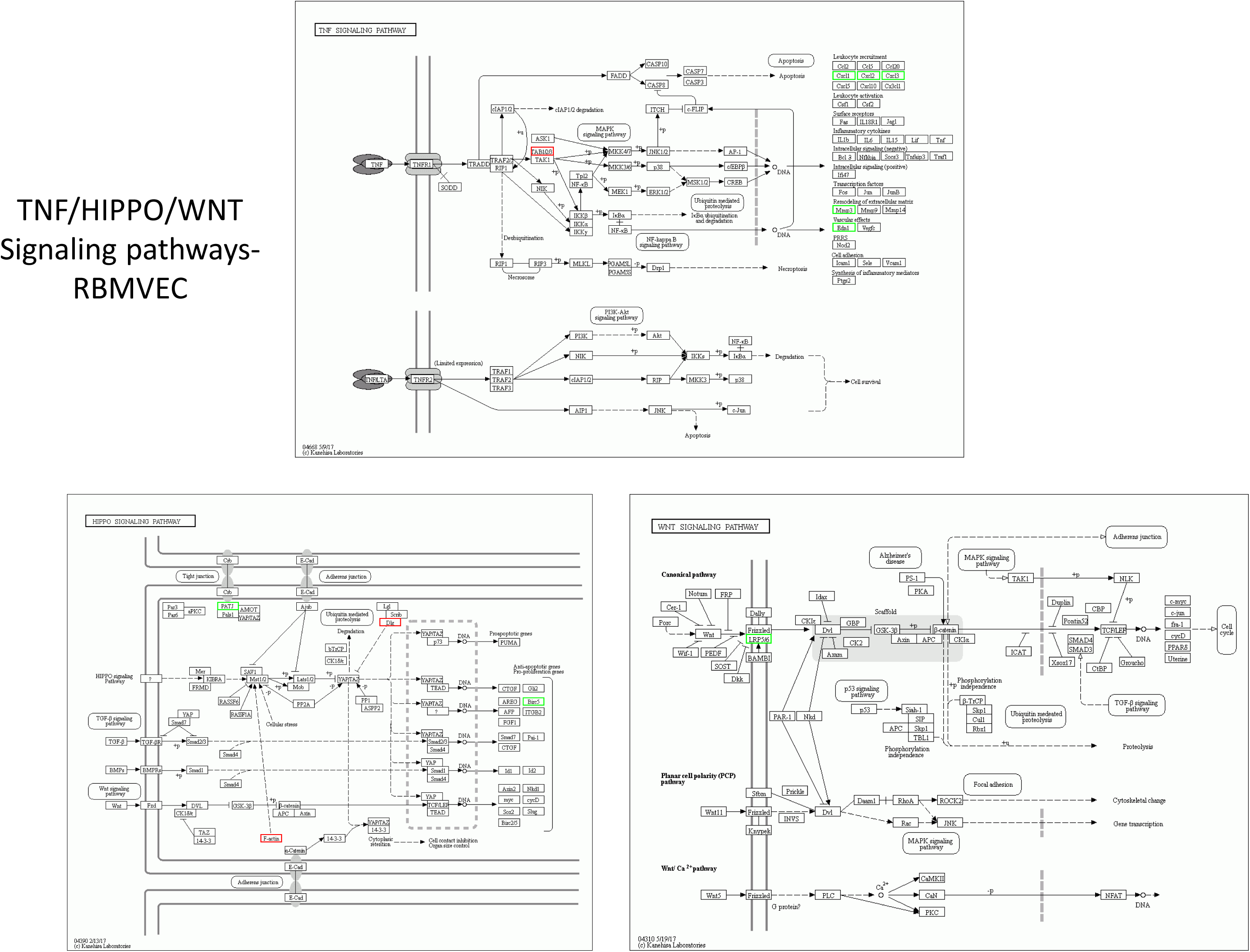
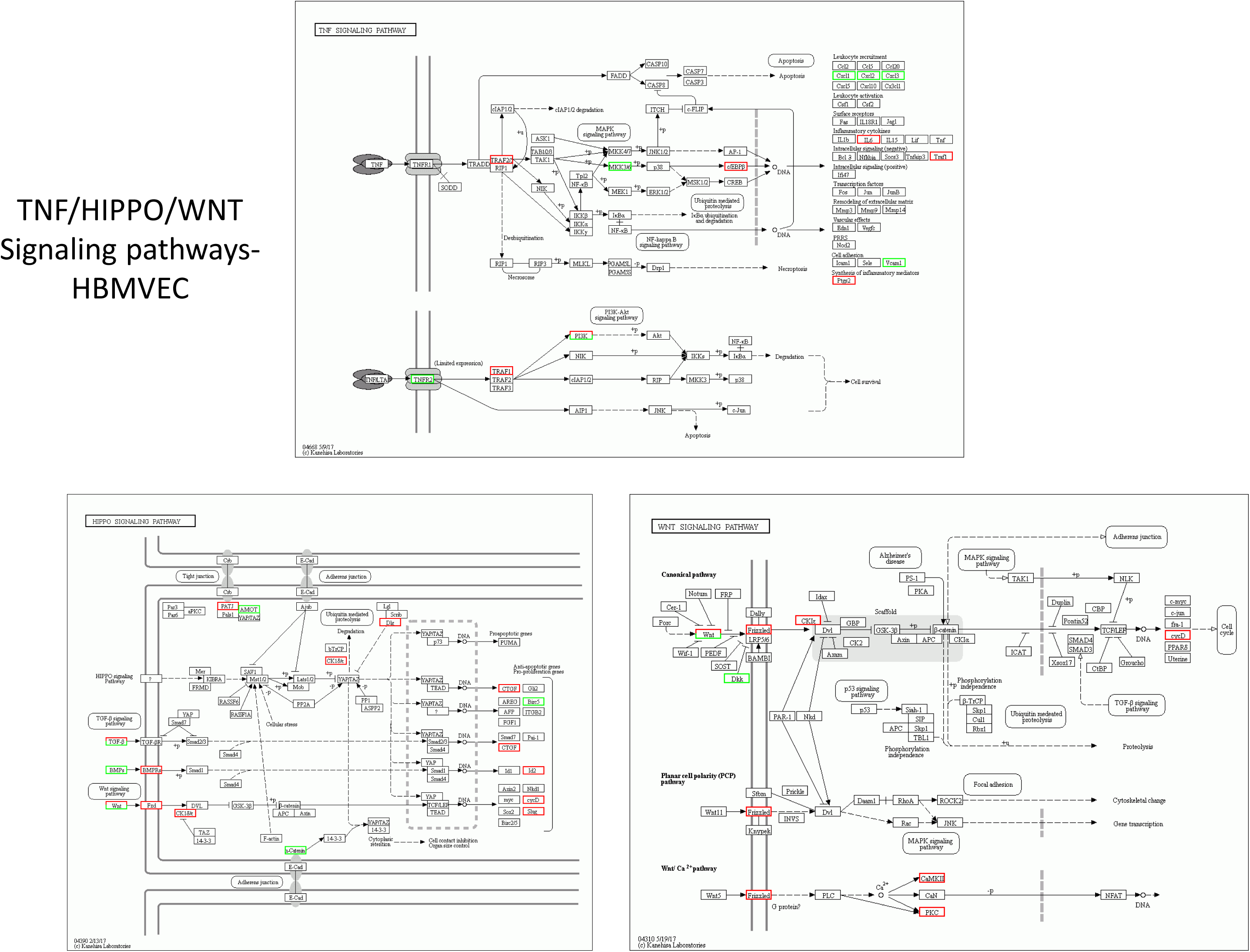
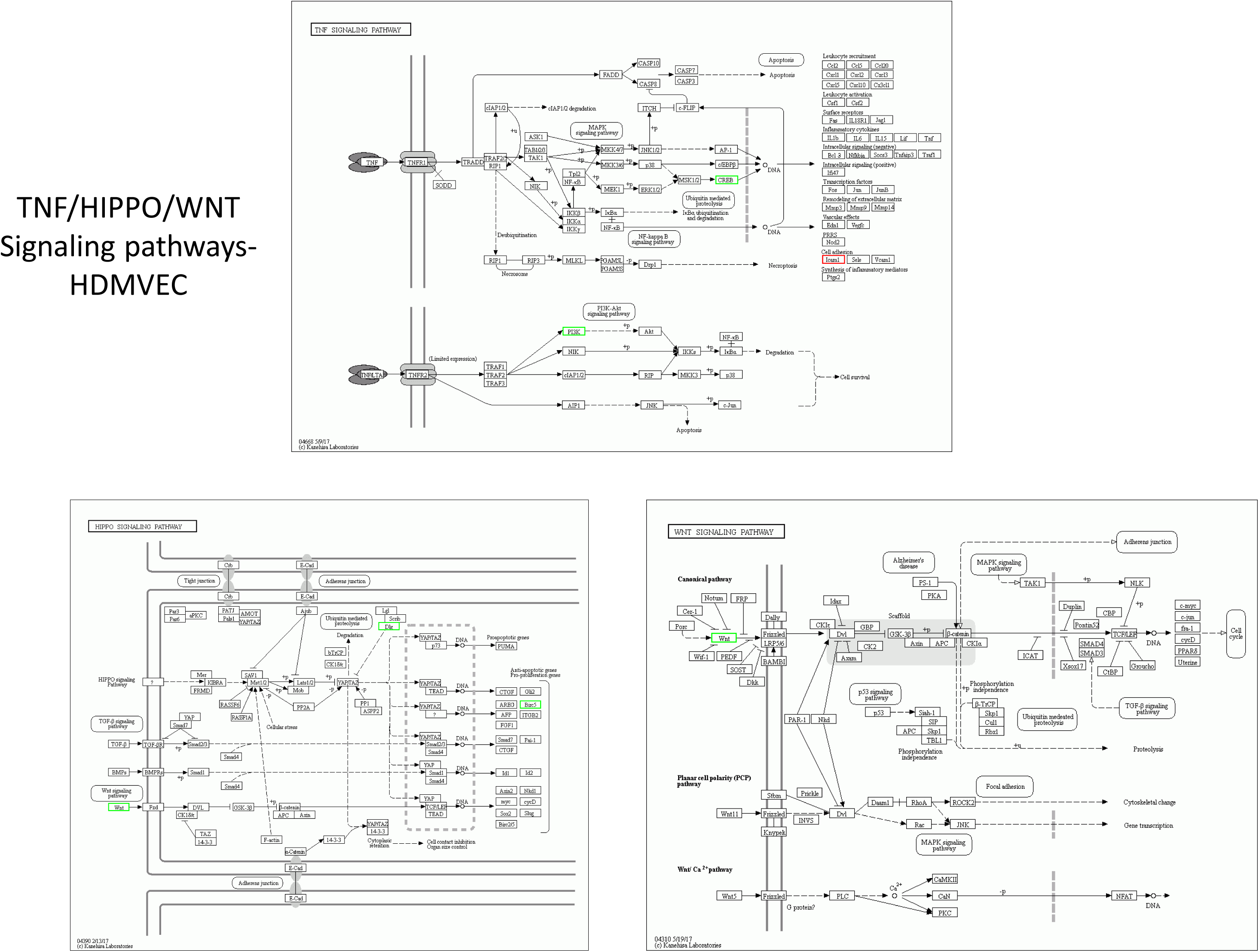
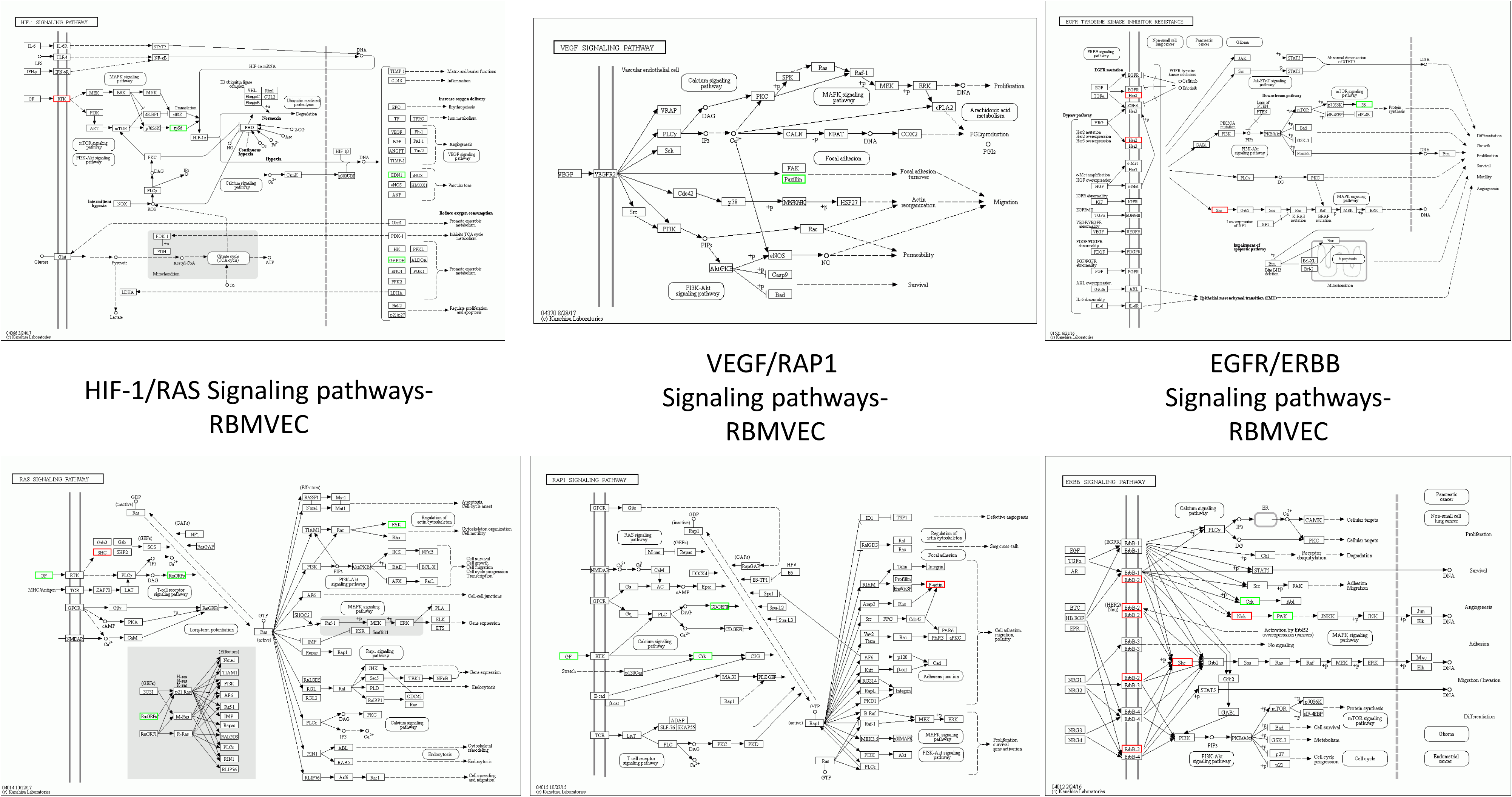
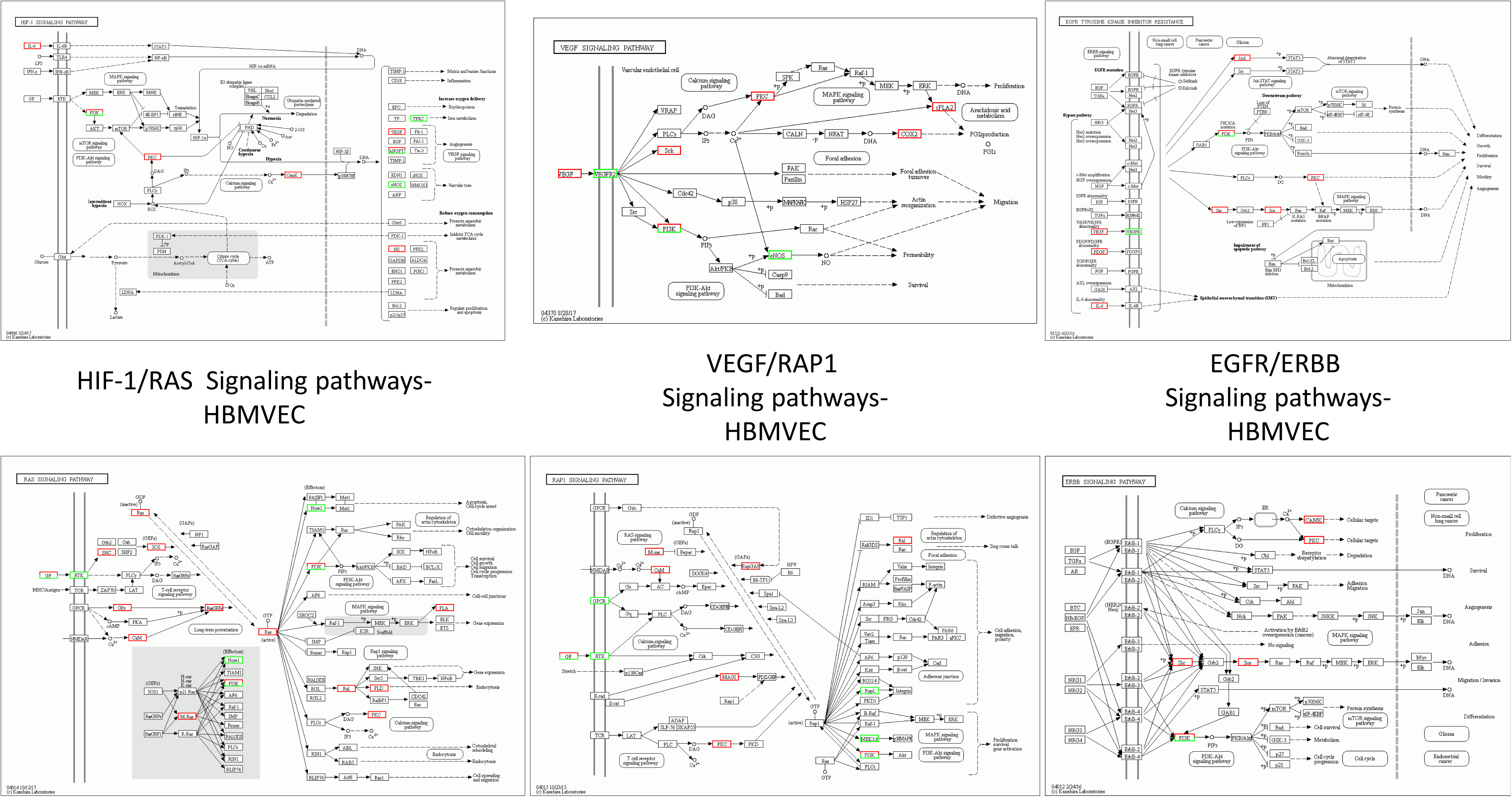
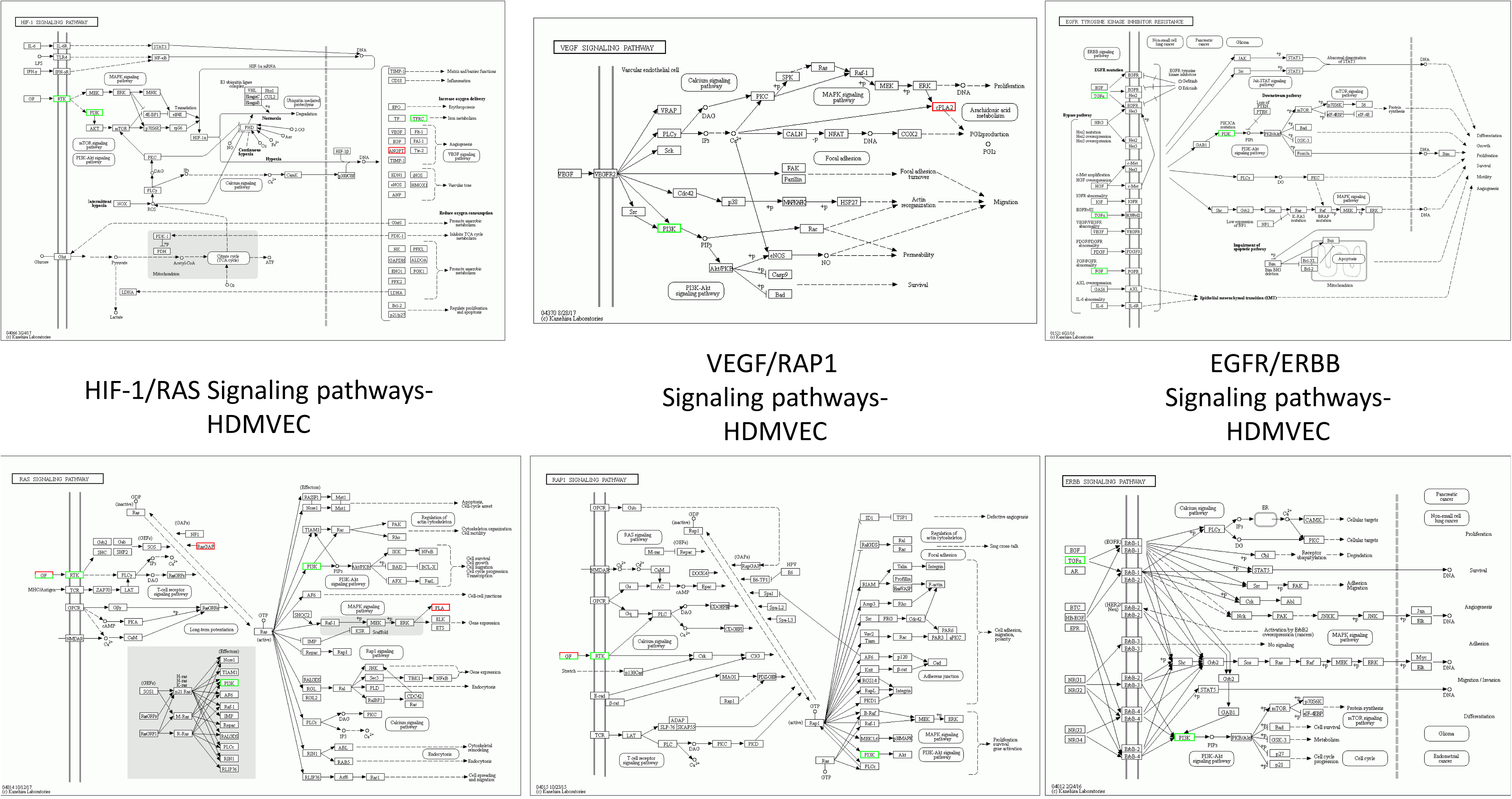
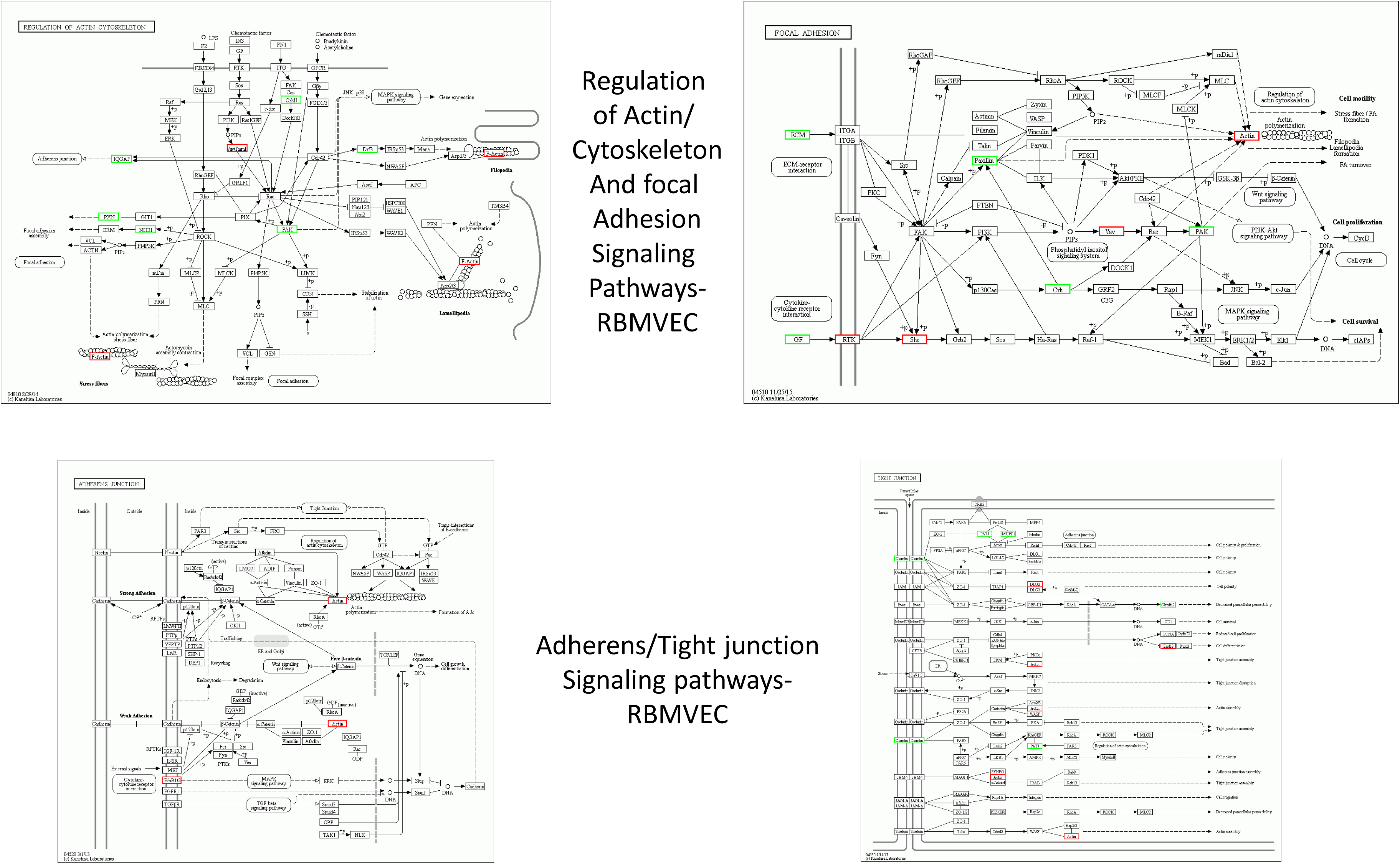
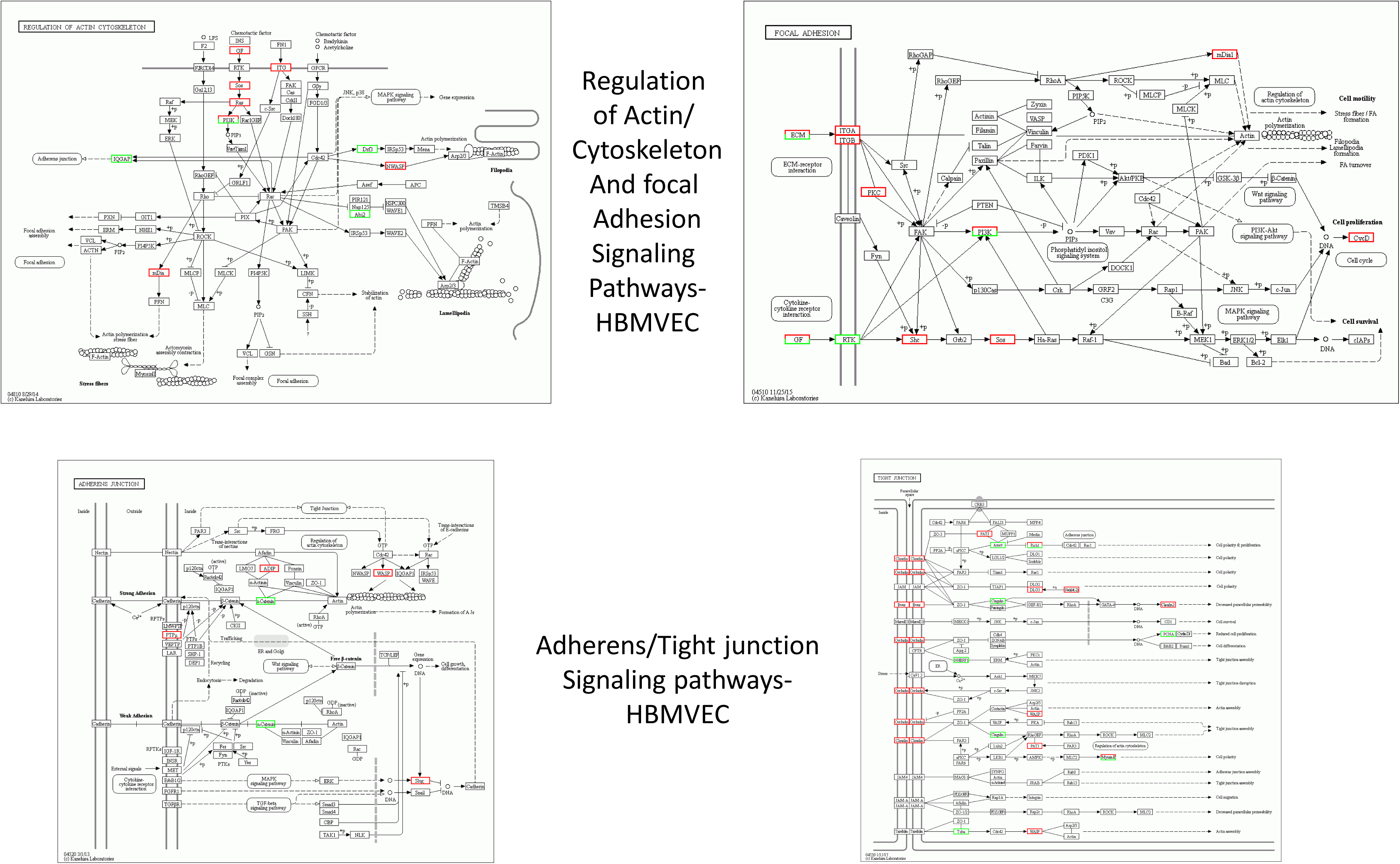
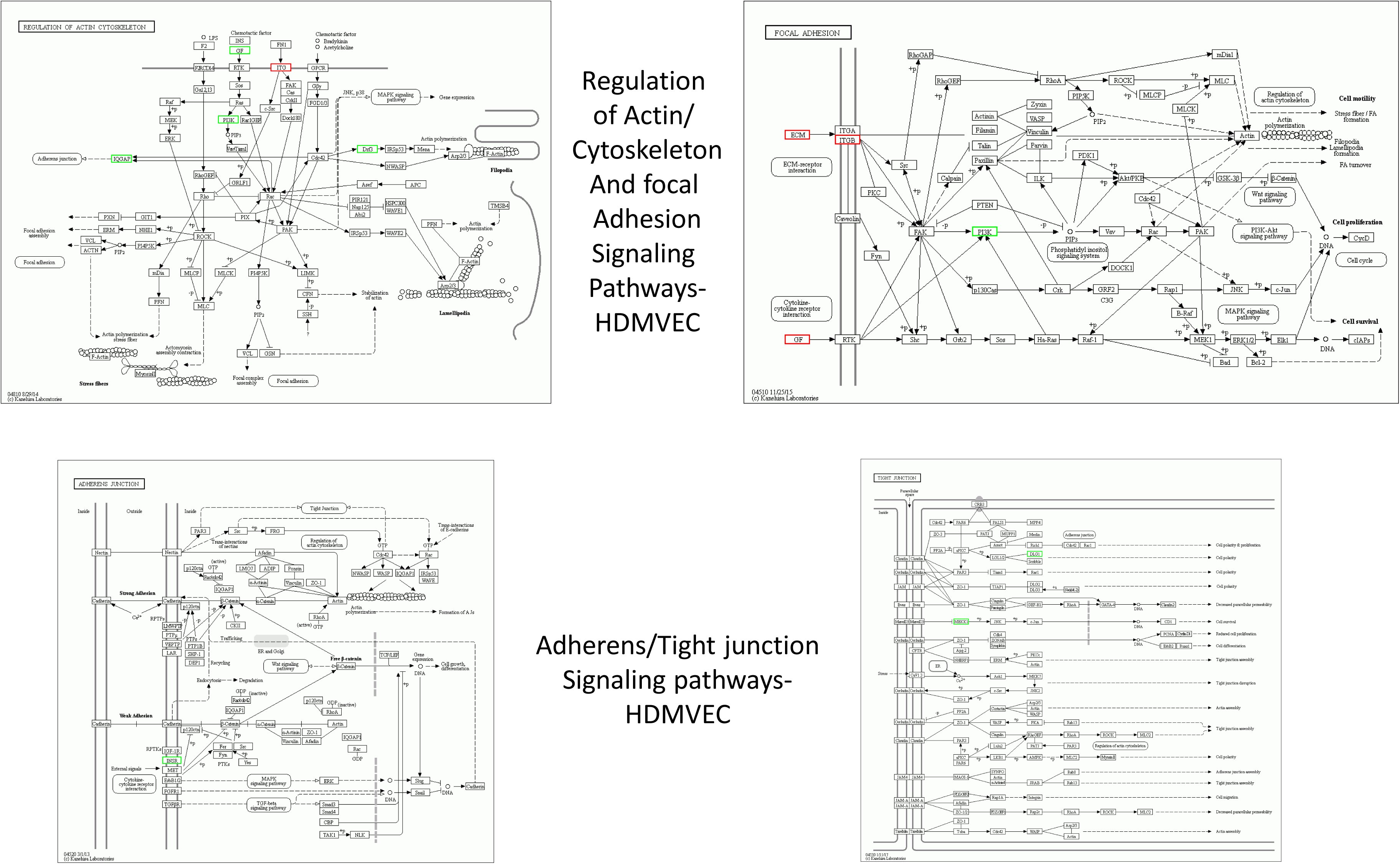
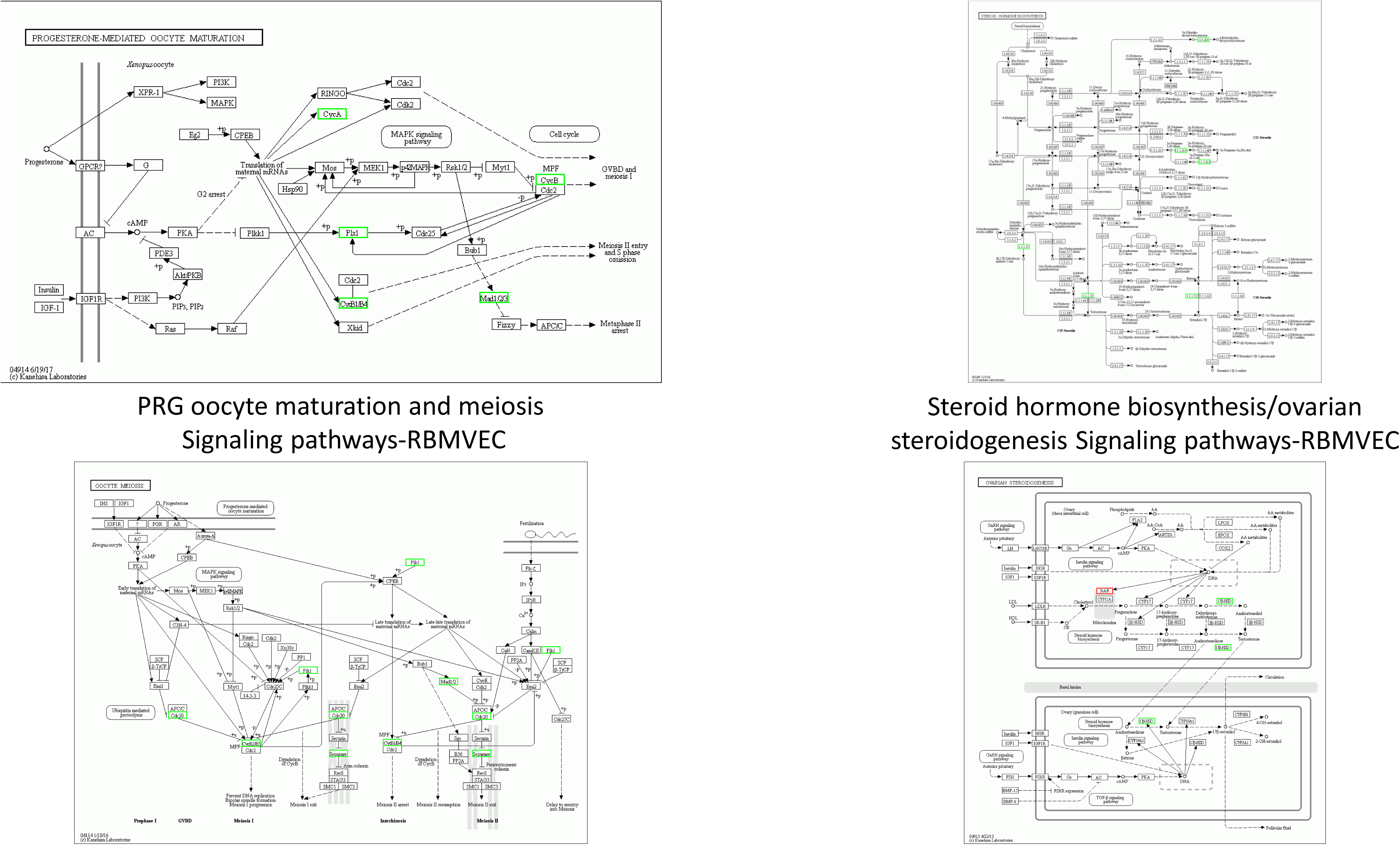
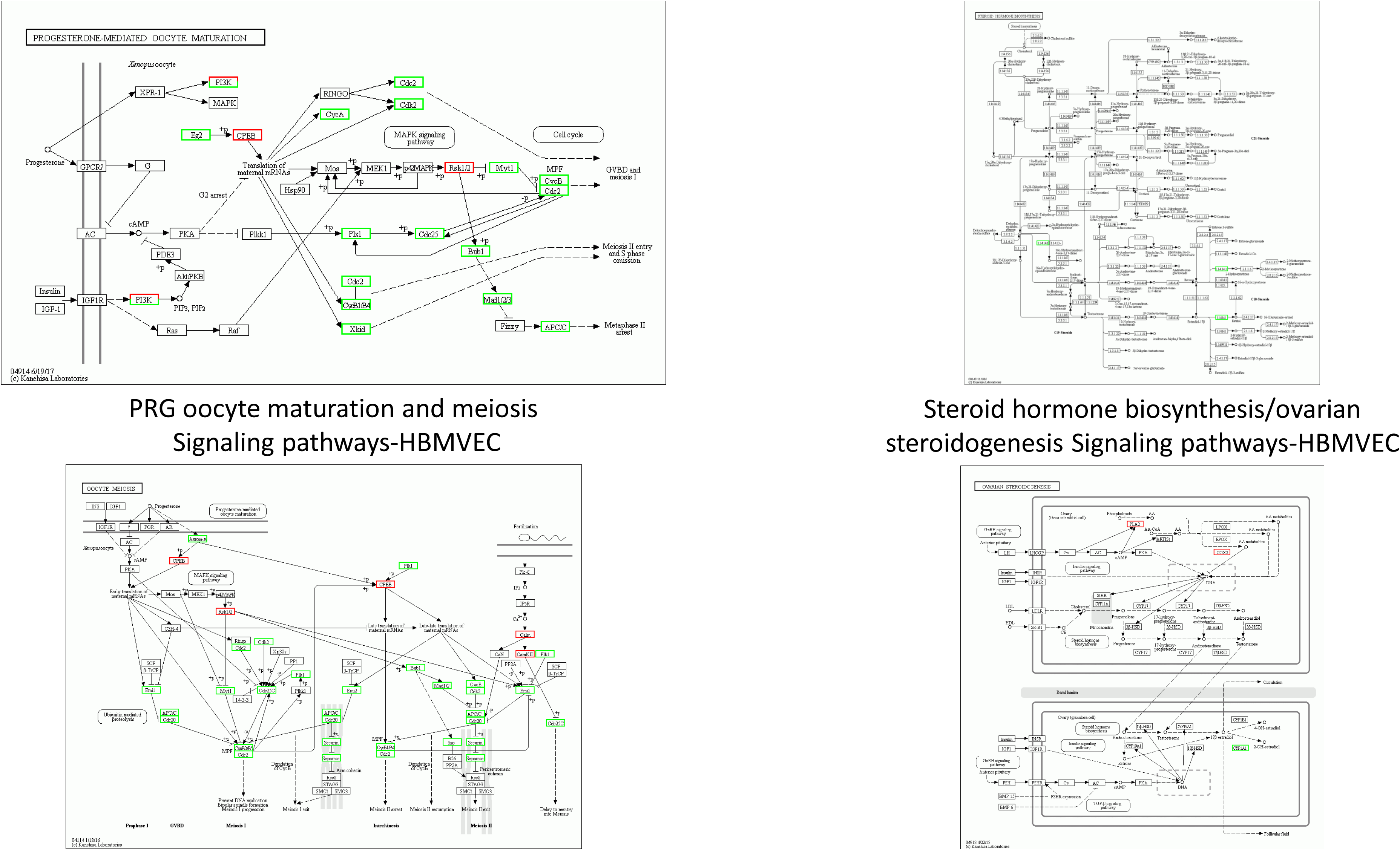
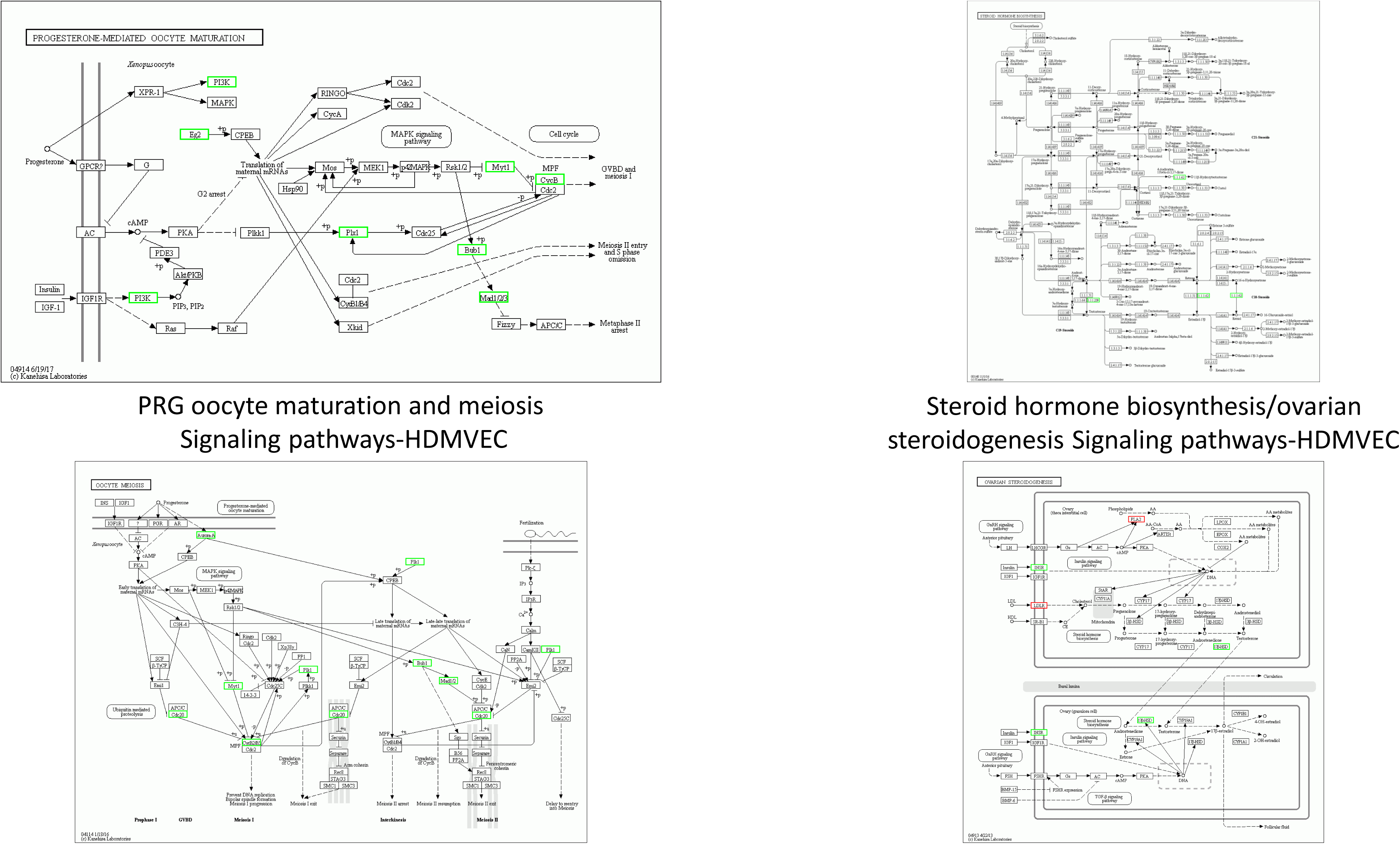
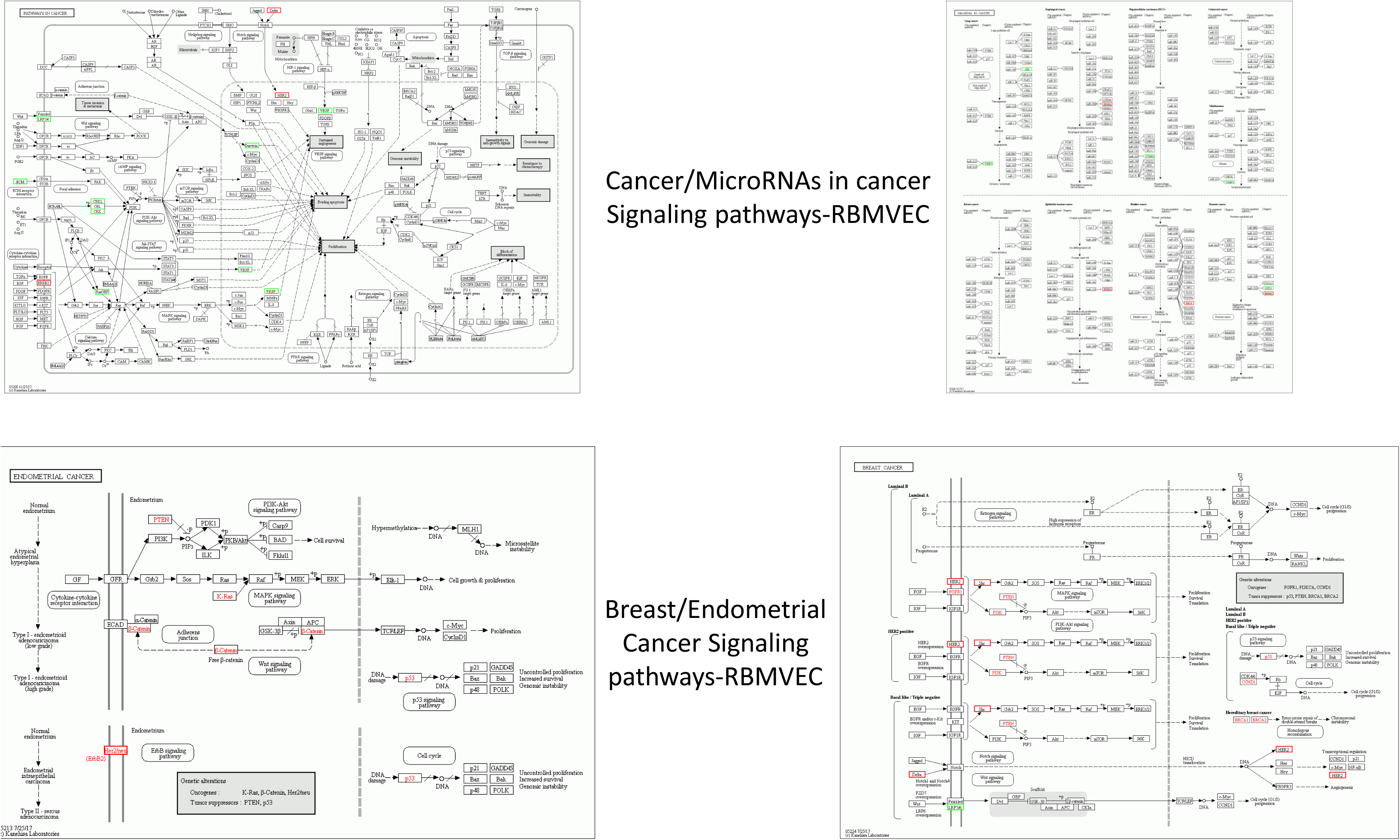
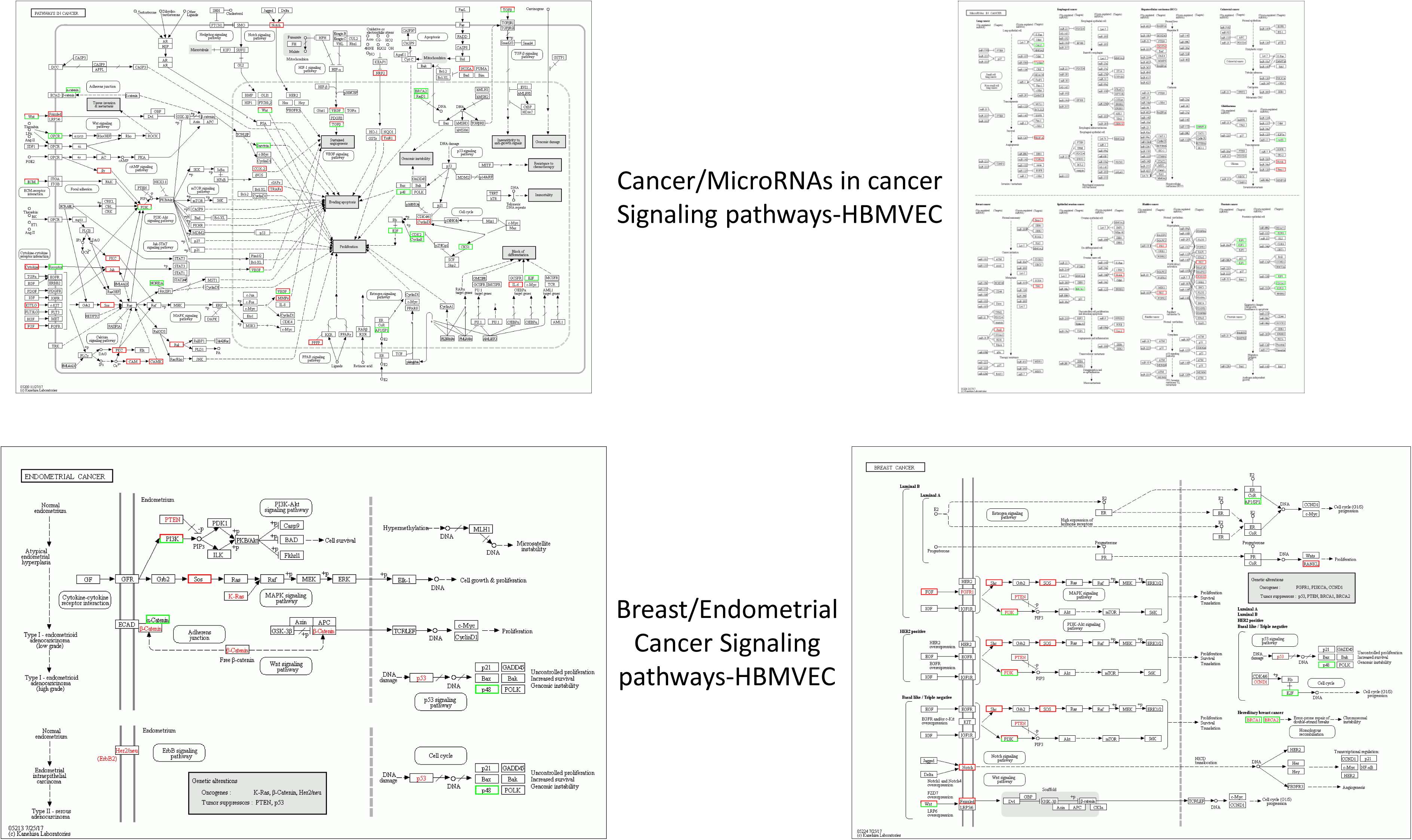
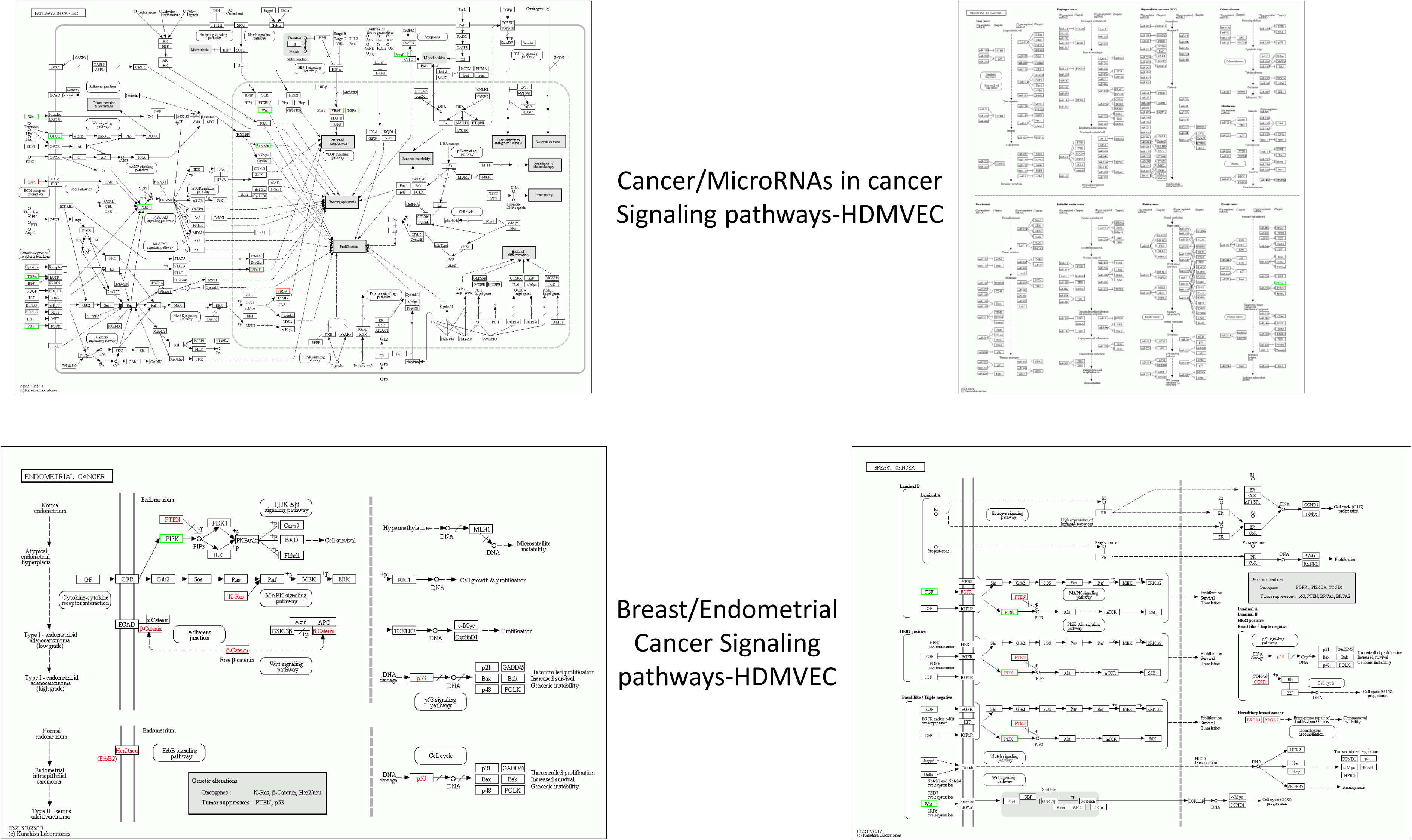
**KEGG pathways analysis of RNAseq data for HBMVEC, RBMVEC and HDMVEC:** Using Differentially Expressed Genes (DEGs), we performed KEGG pathway classification and functional enrichment. With the KEGG annotation results, we classified DEGs according to official classification, and we also performed pathway functional enrichment using phyper, a function of R. **A-C**). Cell cycle, cellular senescence, and apoptosis signaling pathways in RBMVEC (**A**), HBMVEC (**B**) and HDMVEC (**C**). **D-F**). mTOR, P13K-AKT, AMPK, FOXO, NOTCH (RBMVEC and HBMVEC only) and MAPK signaling pathways in RBMVEC (**D**), HBMVEC (**E**) and HDMVEC (**F**). **G-I**). TNF, HIPPO and WNT signaling pathways in RBMVEC (**G**), HBMVE**H**) and HDMVEC (**I**). **J-L**). HIF-1, RAS, VEGF, RAP1, EGFR and ERBB signaling pathways in RBMVEC (**J**), HBMVEC (**K**) and HDMVEC (**L**). **M-O**). Regulation of Actin/cytoskeleton, focal adhesion and adherens/tight junction signaling pathways in RBMVEC (**M**), HBMVEC (**N**) and HDMVEC (**O**). **P-R**). Progesterone oocyte maturation, oocyte meiosis, steroid hormone biosynthesis and ovarian steroidogenesis signaling pathways in RBMVEC (**P**), HBMVEC (**Q**) and HDMVEC (**R**). **S-U**). Cancer, MicroRNAs in cancer, breast cancer and endometrial cancer signaling pathways in RBMVEC (**S**), HBMVEC (**T**) and HDMVEC (**U**). For all figures, up-regulated genes are marked with red borders and down-regulated genes with green borders. Non-changed genes are marked with black borders.

**Fig. S4: Raw data have been deposited in Table S4, no additional data in this figure.**

**Fig. S5.**
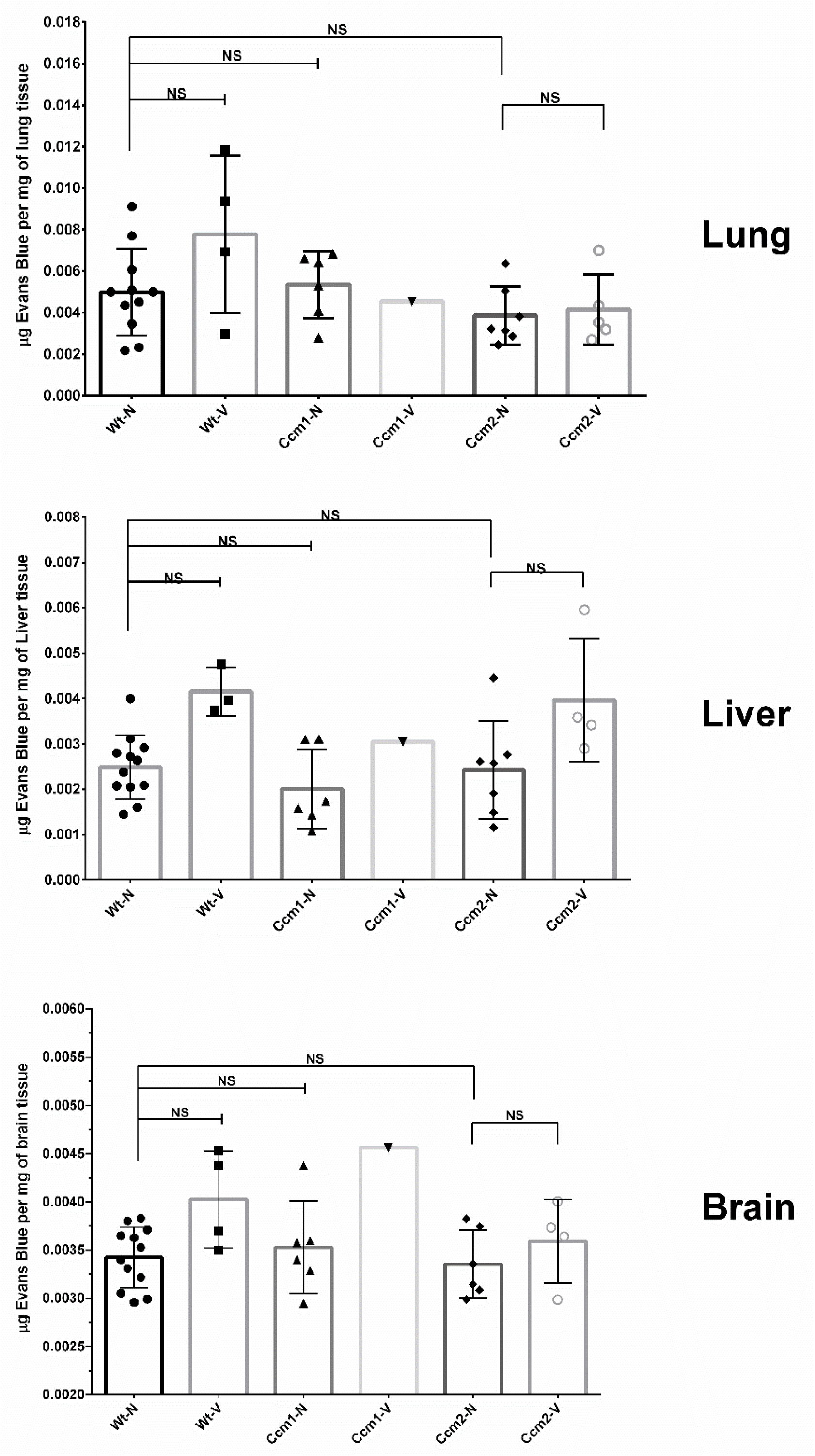
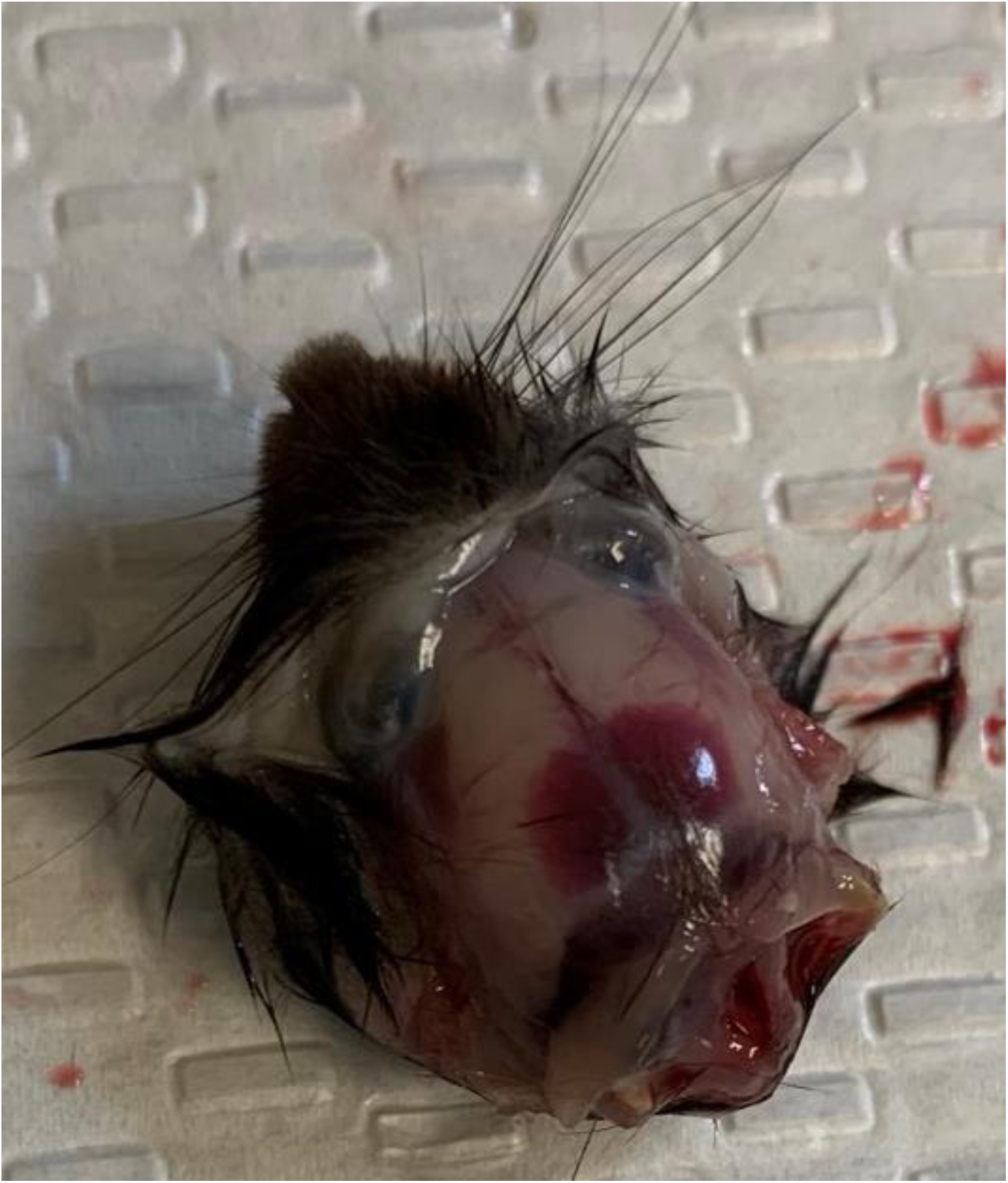
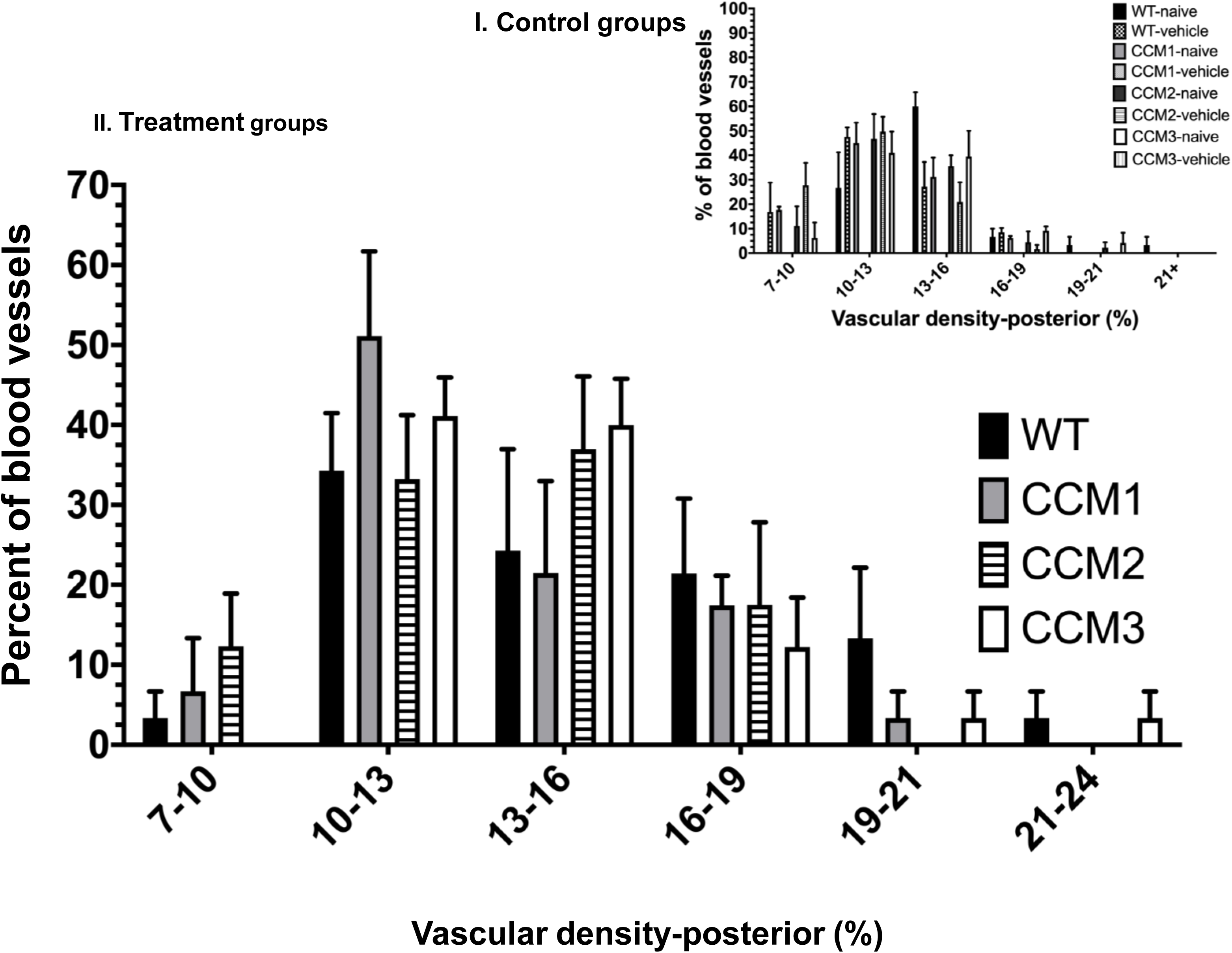
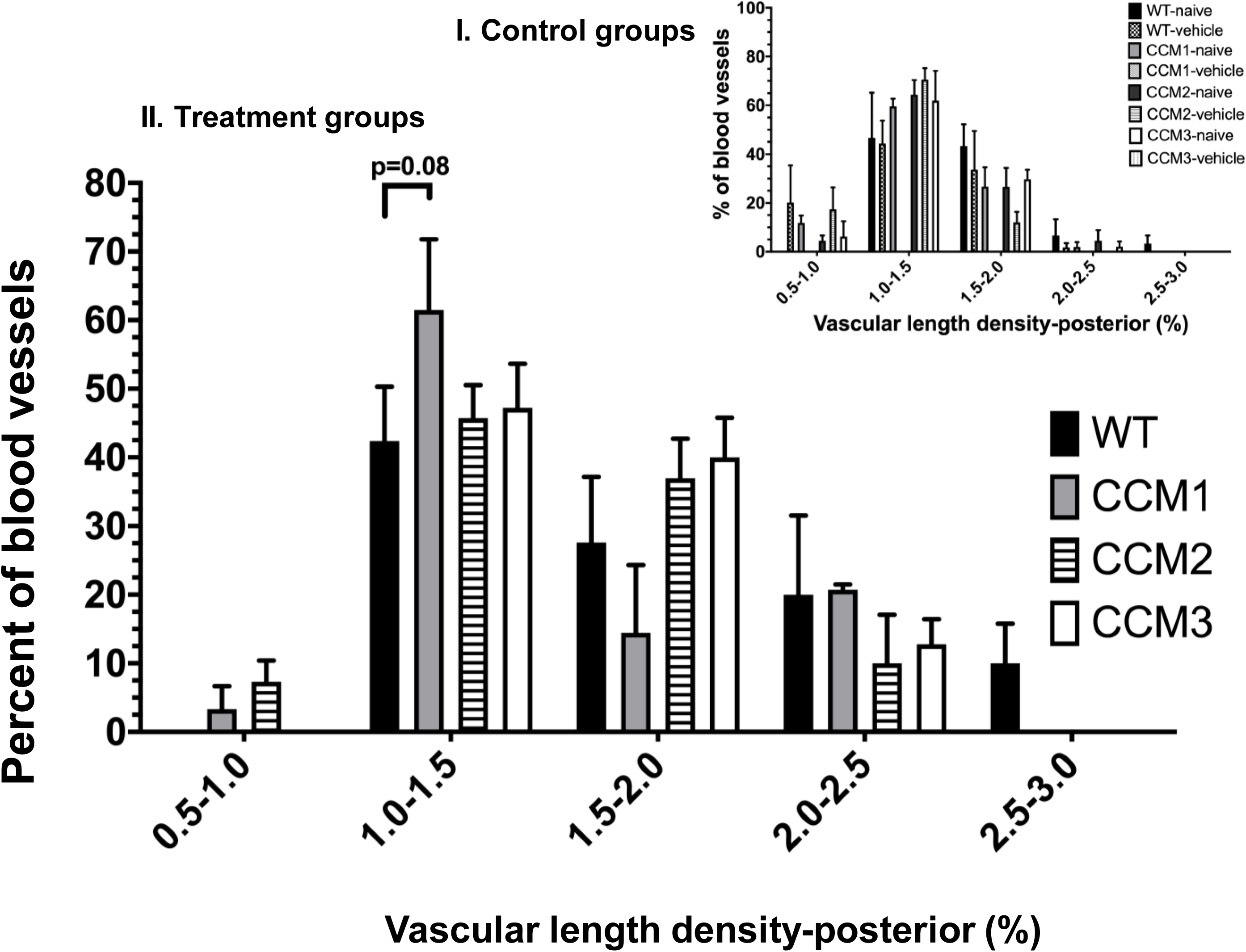
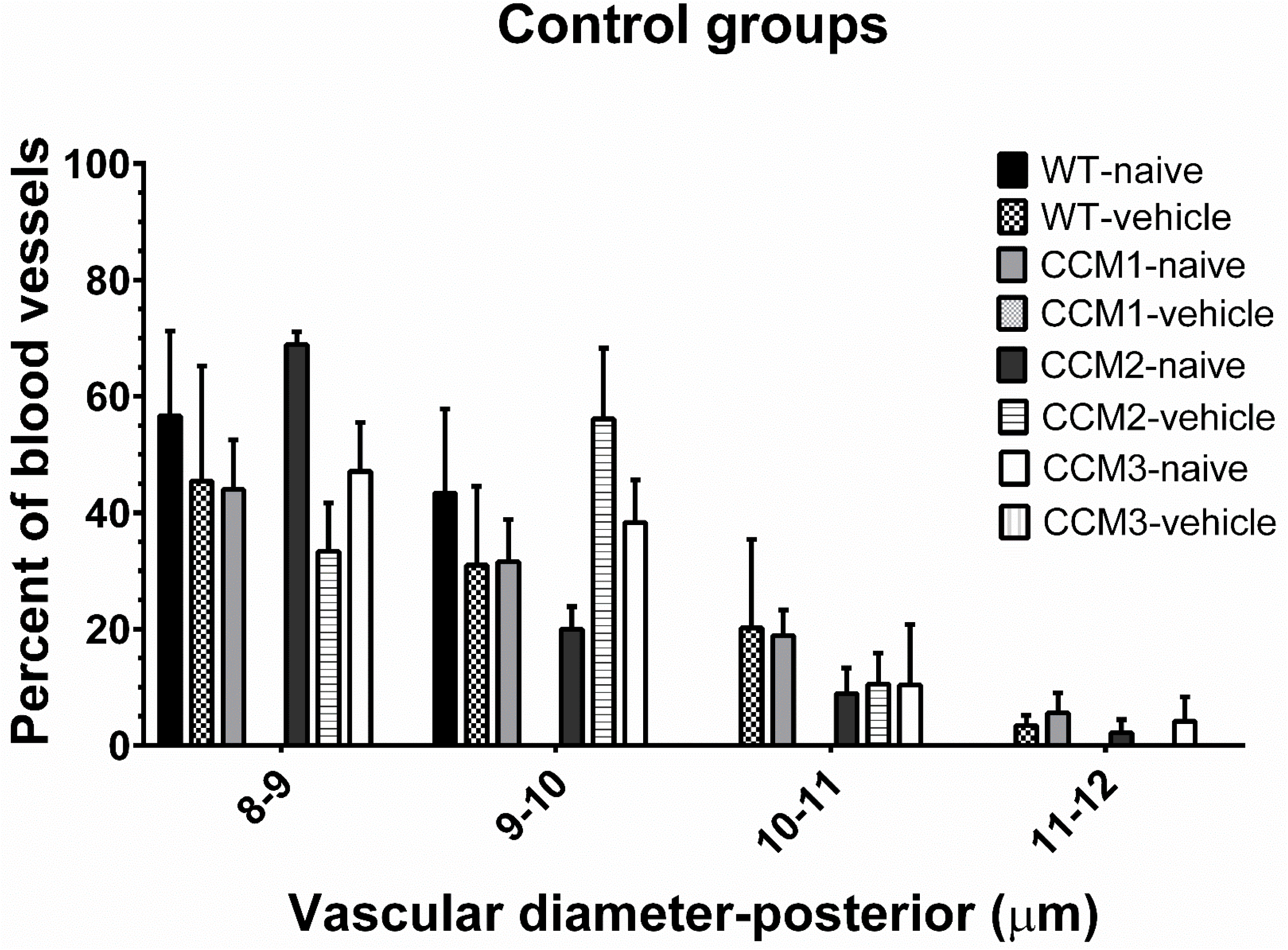
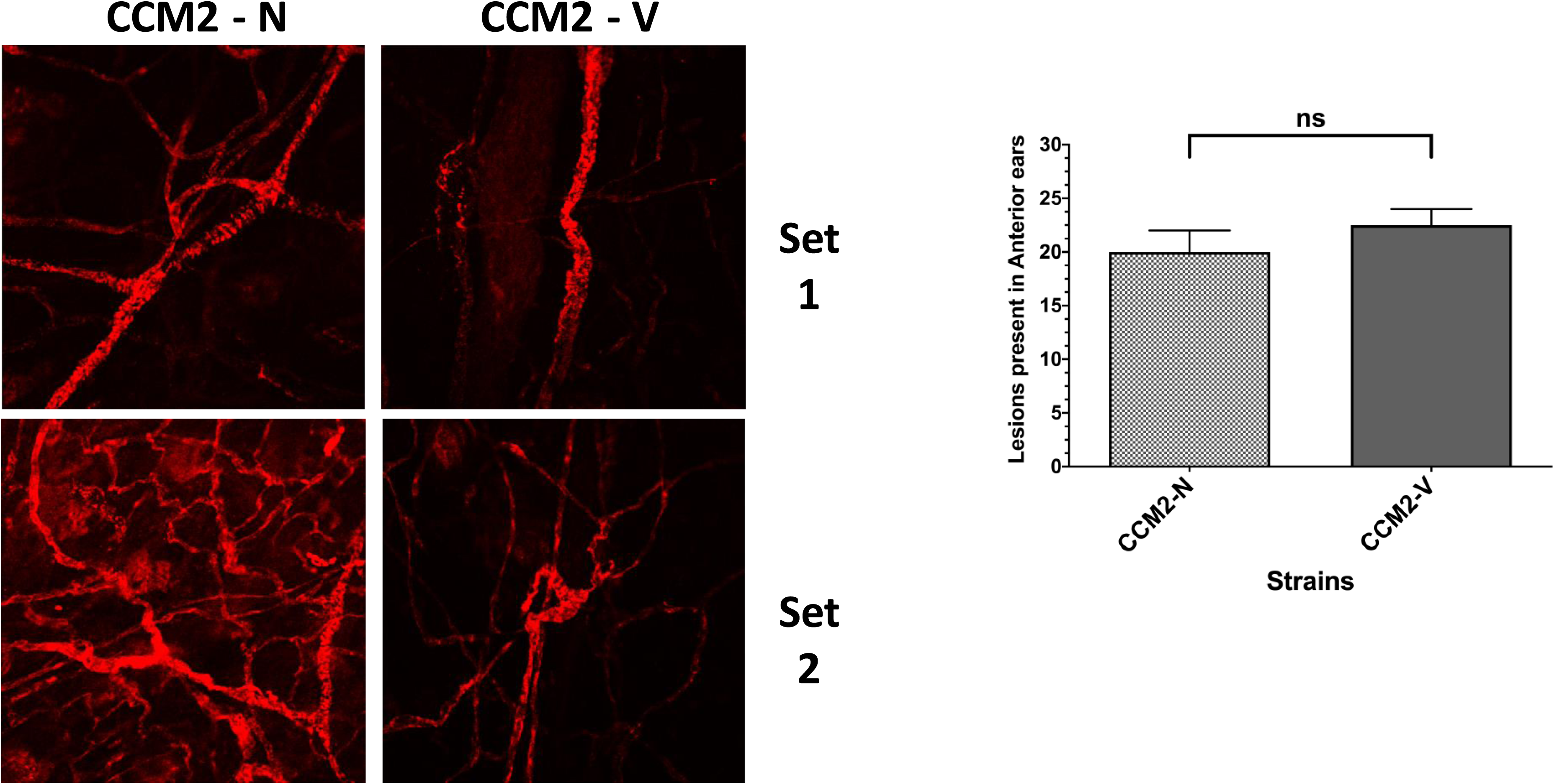
**The combined action of progesterone and mifepristone (PRG+MIF) on mPRs (PAQRs) in PR**(-**) microvascular ECs increase microvascular permeability, and is sufficient for the formation of hemorrhagic CCM lesions *in-vivo*.** A. Increased microvascular permeability is not caused by either by vehicle (peanut oil) or genotypes. Normal microvascular function in the lungs, liver, and brain were observed, with no increased microvascular permeability detected in either vehicle control or naïve groups among Ccms (1, 2, 3) mutant and WT mice. Both Ccms (1, 2, 3) mutant and WT (C57BL/6J) mice were injected with Peanut oil 5 days a week for 30 days for vehicle controls, or without any injections set for naïve controls. **B.** Evidence of intracranial bleeding in Ccm3 mutant mouse in the 90 day treatment group. After Euthanasia, evaluation for early hemorrhagic events was performed to assess any signs of intracranial bleeding, resulting from combined steroid actions. We did not observe any intracranial bleeding in WT PRG+MIF/vehicle treated mice or WT naïve mice. **C.** Vascular density of subcutaneous vessels in posterior sections of ears. Subcutaneous vasculatures, another common location for CCM lesions, were also systematically examined. Percent averages of vessels, within the density categories on the x axis, were then pooled across replicates. Subcutaneous vessels were further classified into six subgroups based on the range of the vessel density (counts/view): group-I (7-10), group-II (10-13), group-III (13-16), group-IV (16-19), group-V (19-21), and group-VI (21-24). I). No significant difference was observed among Ccm1, Ccm2, Ccm3 and WT in either I) control or II) treatment; **D.** Vascular length density in posterior sections of ears was determined through calculation of a ratio of skeletonized vasculature area to total area. Percent averages of vessels, within the length density categories on the x axis, were then pooled across replicates. Subcutaneous vessels were further classified into five subgroups based on the range of the vessel length density (ratio): group-I (0.5-1.0), group-II (1.0-1.5), group-III (1.5-2.0), group-IV (2.0-2.5), and group-V (2.5-3.0). Although increased numbers of Ccm1 mutant was notable in group-III (1.5-2.0), compared to WT, however in general, no significant difference was observed among Ccm1, Ccm2, Ccm3 and WT in either I) control or II) treatment; **E.** Vascular diameter in posterior sections of ears was defined with conversion of pixel color to relative thickness of the vessels in that area. Percent averages of vessels, within the diameter categories on the x axis, were then pooled across replicates. Subcutaneous vessels were further classified into four subgroups based on the range of the vessel size (diameters): group-I (8-9 µM), group-II (9-10 µM), group-III (10-11 µM), and group-IV (11-12 µM); no significant differences was observed among Ccm1, Ccm2, Ccm3 and WT controls; **F.** CCMs lesions of subcutaneous vessels in the anterior side of mouse ears were examined and counted at 40X and any lesion size was counted towards the total numbers for each slide. Ccm2 mutant was chosen to determine the baseline variation between naive (untreated) and vehicle treatment (left panel), no difference was observed between two control groups (right panel) (n=3), indicating no notable influence of vehicle injection on the appearance of vessels (both panels). **G.** in Evans blue assays, statistical analysis was generated using one-way ANOVA with either Kruskal-Wallis test or uncorrected Fisher’s LSD test where appropriate. All treated mice upon completion of the last injection were injected with Evan’s Blue Dye (EBD) (500µG/25G mouse) in peanut oil which was allowed to circulate for 3 hours. Subsequently, organs were harvested and snap frozen until ready for use. All tissue samples went through a series of processing before analysis, each sample was loaded into 8 wells to serve as technical replicates, and analyzed using a flex station 3 (see methods). Fluorescence data are then converted to µg/mL based on standard curves generated in the same PBS/TCA/Ethanol buffer used for extraction, and normalized based off the tissue weight (µg Evans Blue/mg tissue) followed with controls. In ear tissues, statistical significance was performed using unpaired students T-test (*, ** and *** above graphs indicate P ≤ 0.05, 0.01, and 0.001, respectively).

**Fig. S6.**
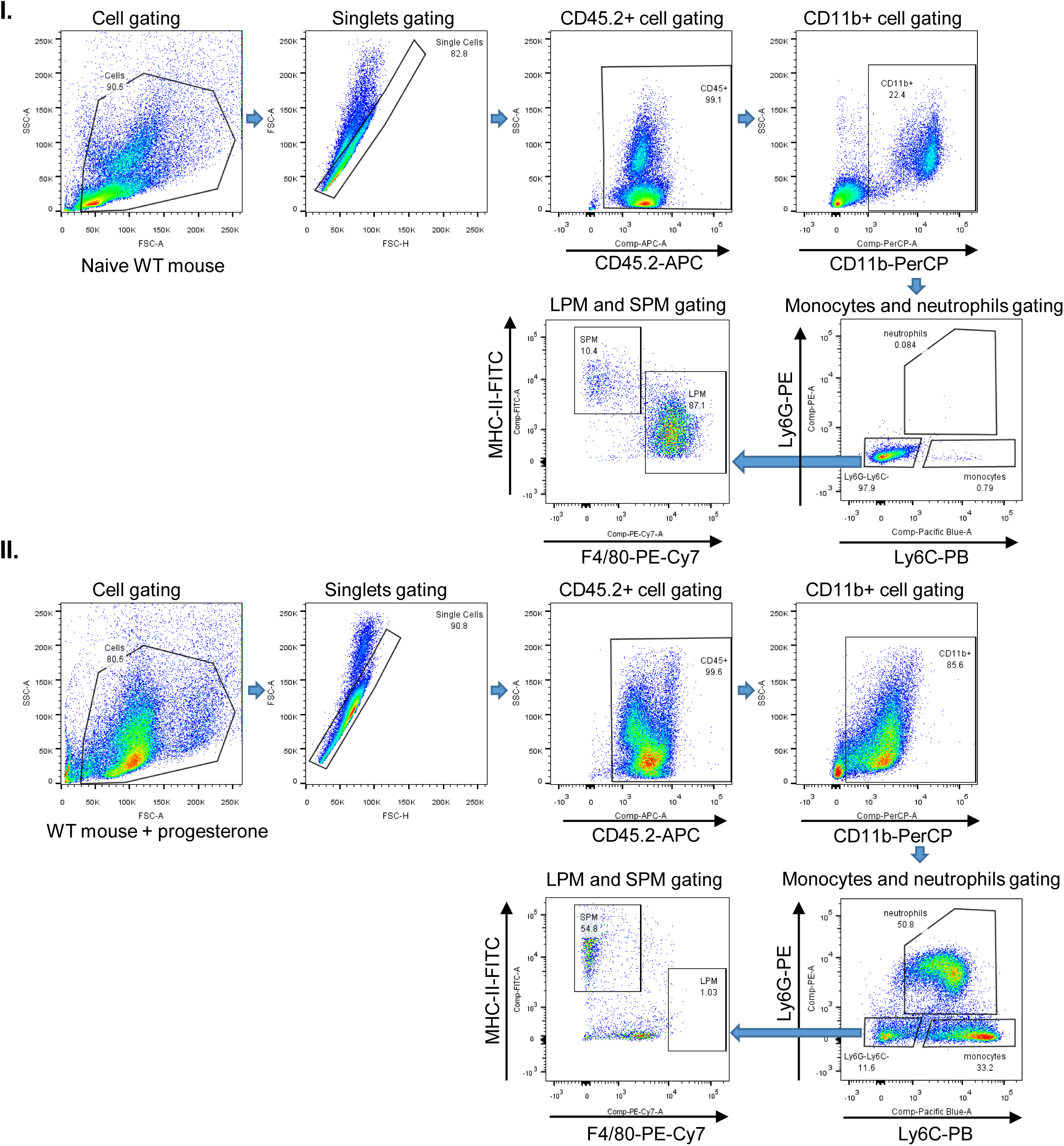
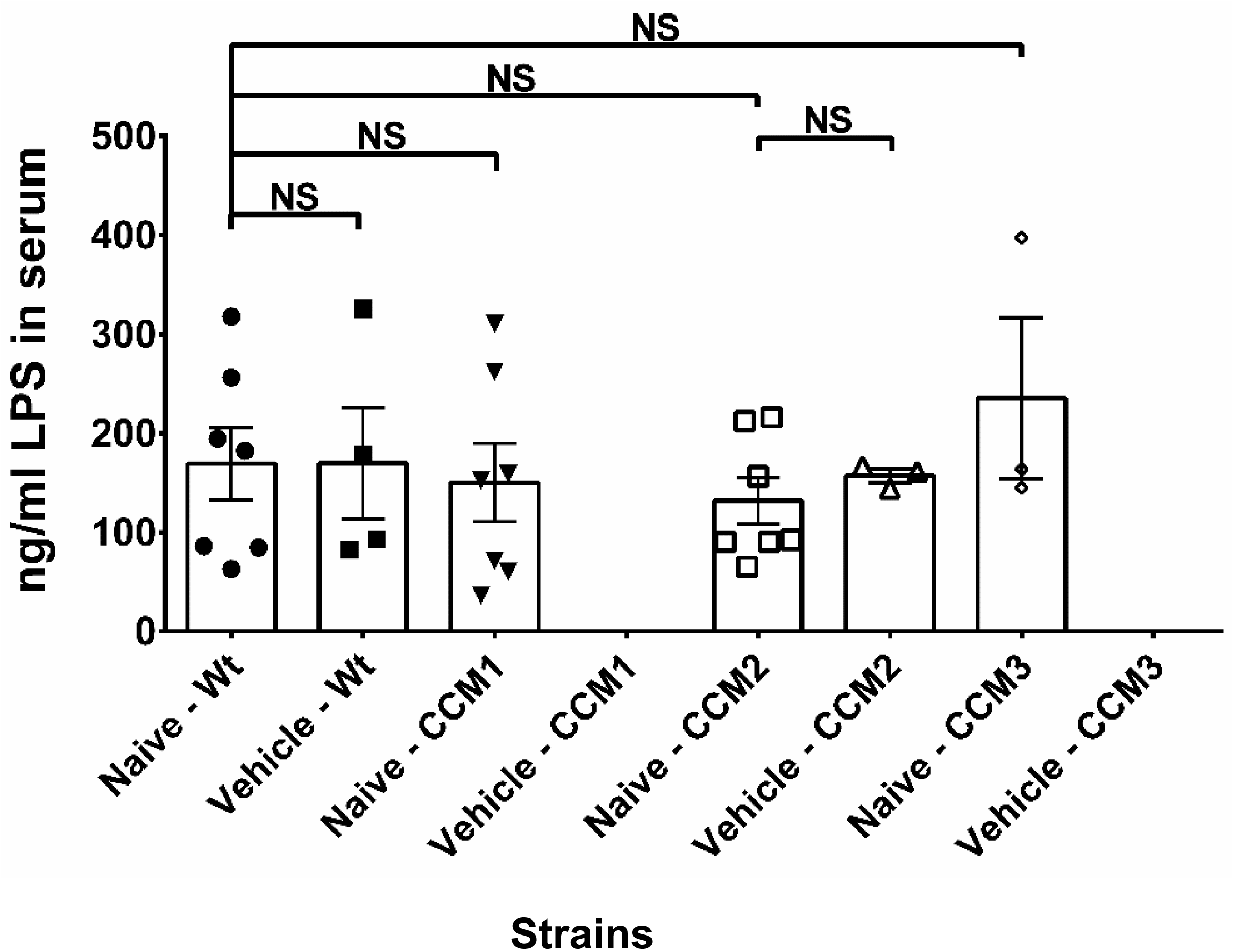
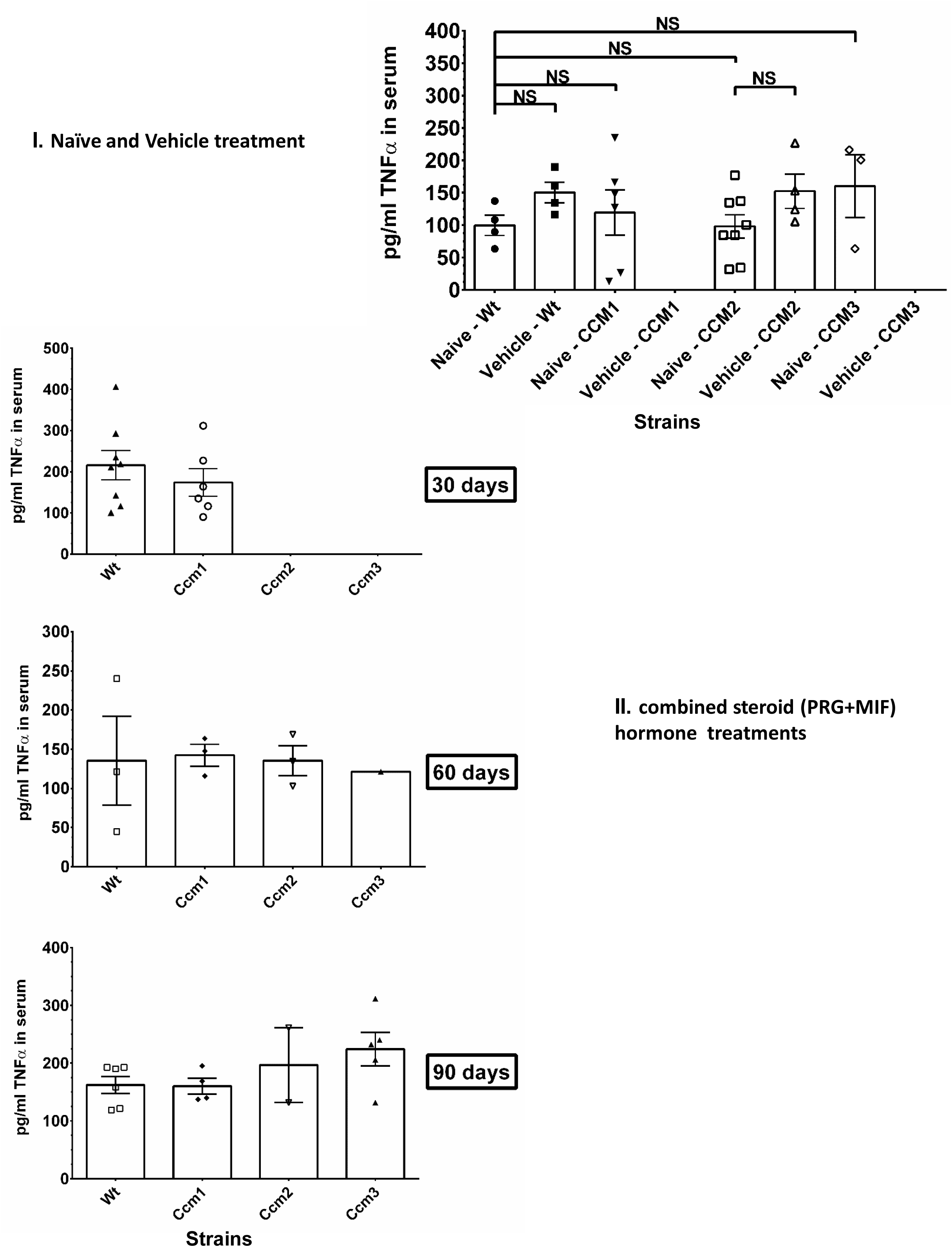
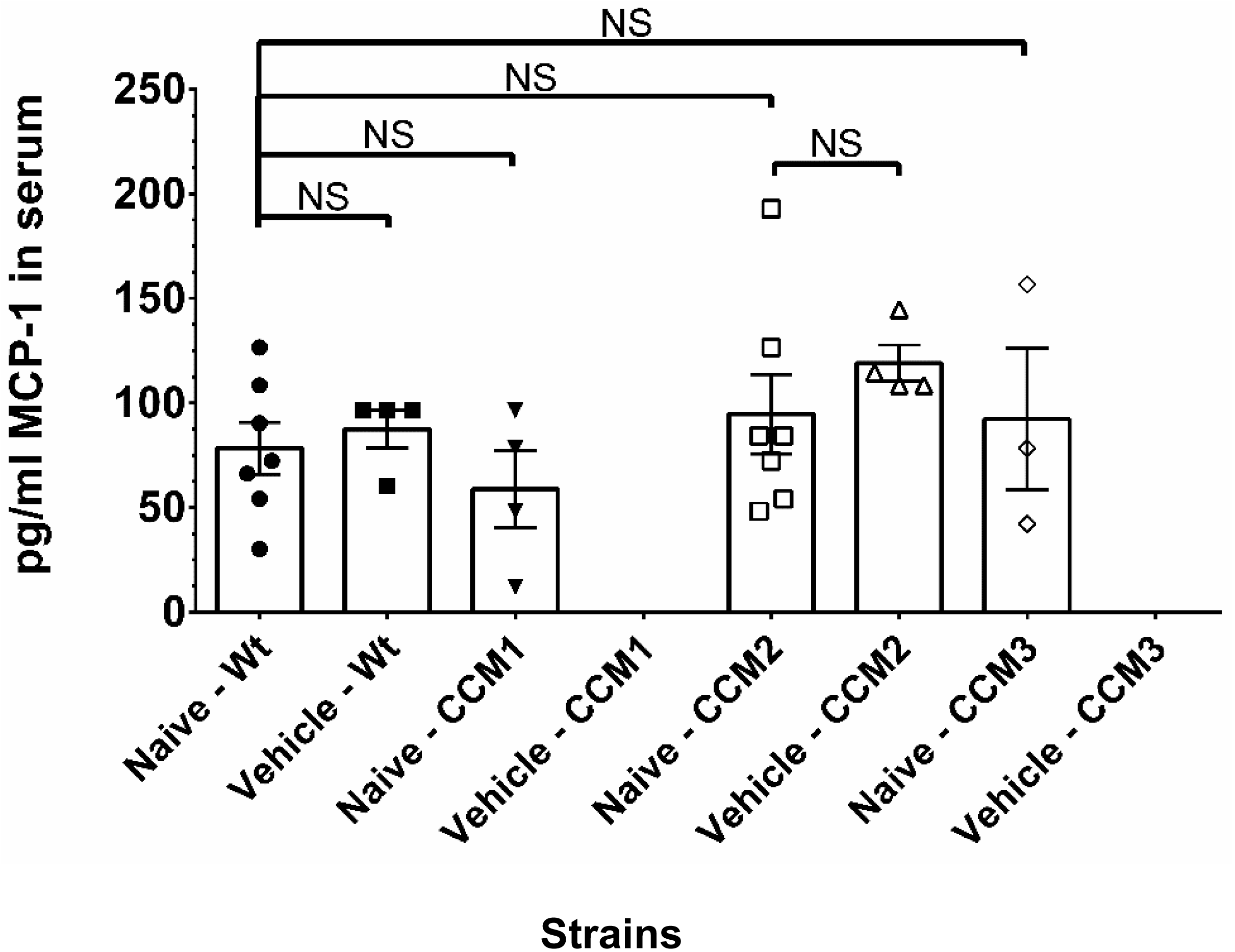
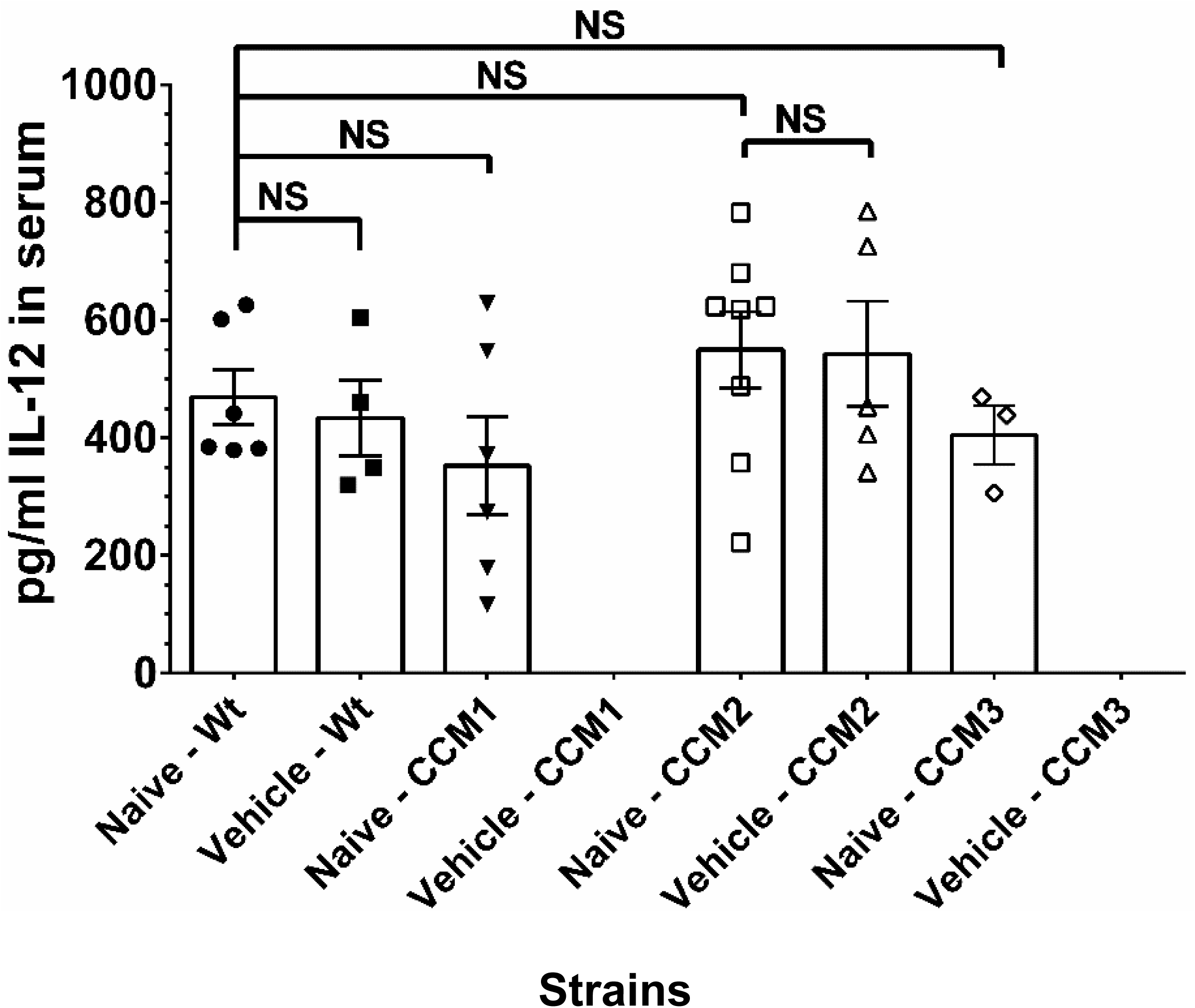
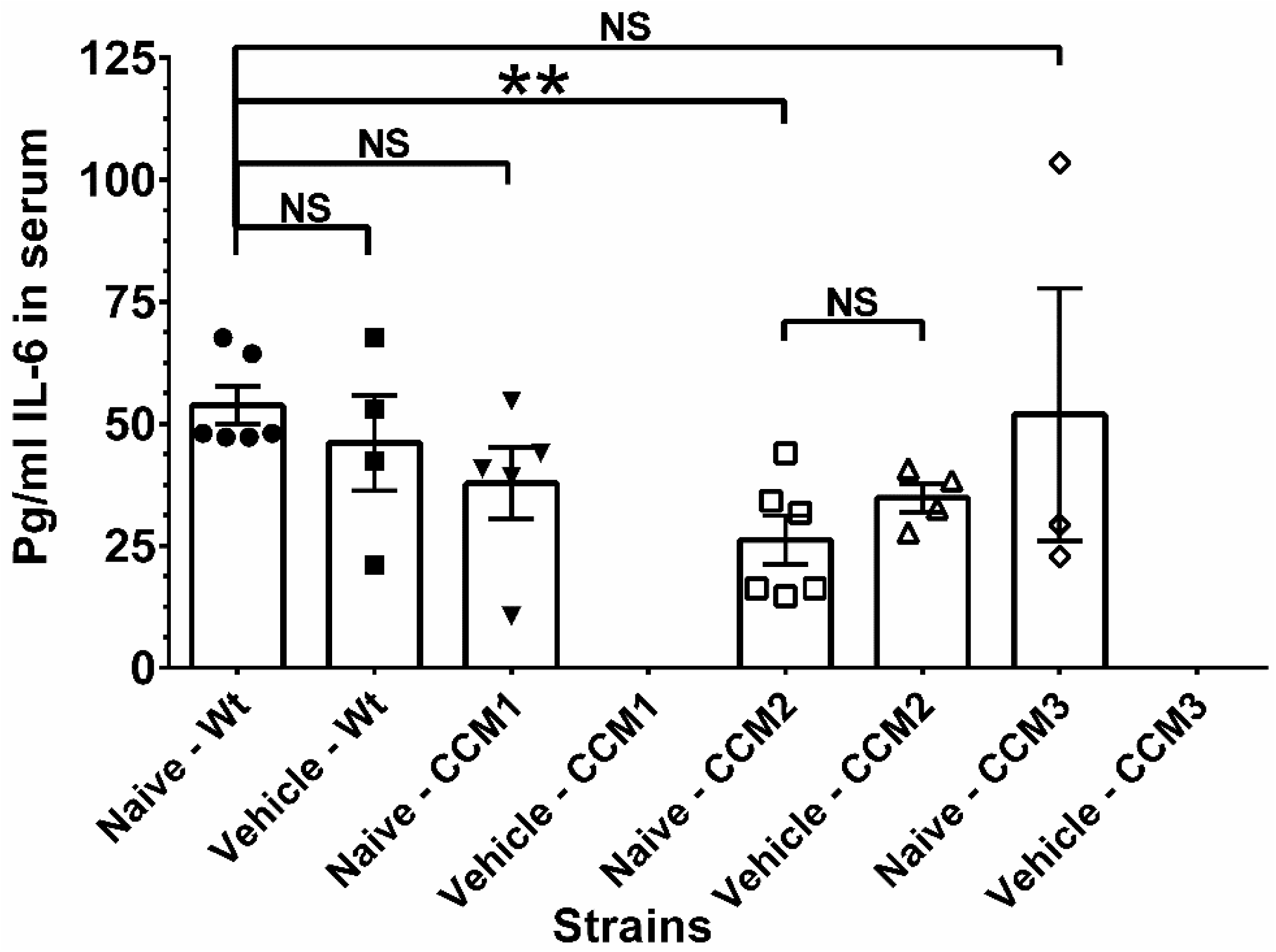
**The disrupted brain-blood barrier (BBB) is neither caused by local inflammatory reactions nor associated with systemic inflammations in Ccms mice. A**. Localized inflammations are not caused by either combined steroid (PRG+MIF) action or genotypes. FACS gating strategy to quantify myeloid cells in the peritoneal lavage of mice under various treatments. Monocytes (CD45.2^+^CD11b^+^ Ly6G^-^Ly6C^high^), neutrophils (CD45.2^+^CD11b^+^ Ly6C^int^Ly6G^high^), large peritoneal macrophages (LPM; CD45.2^+^CD11b^+^ Ly6G^-^Ly6C^-^ F4/80^hi^MHC-II^lo^) and small peritoneal macrophages (SPM; CD45.2^+^CD11b^+^ Ly6G^-^Ly6C^-^F4/80^low^MHC-II^high^) were sorted and quantified. Typical data are shown for I) a naïve WT mouse, and II) a WT mouse injected with combined steroid (PRG+MIF) treatment. **B.** Systemic inflammations are not associated with non-immunogenic LPS from *H. pylori* bacteria in Ccms mutant mice. Systemic inflammations are low and could be suppressed with Ccms mutant genotypes. The disrupted brain-blood barrier (BBB) is not associated with non-immunogenic LPS from H*elicobacter* pylori. Lipopolysaccharide-Based Enzyme-Linked Immunosorbent Assay (LPS-ELISA) was used to measure LPS concentration in mouse serums. LPS concentrations were measured and quantified by ELISA in WT, Ccm1, Ccm2, Ccm3 mutant mice under naïve (untreated) and Vehicle [vehicle (peanut oil)-treated] conditions. No obvious difference was detected among Naïve-WT, Vehicle-WT, Naïve-Ccm1, Naïve-Ccm2, Vehicle-Ccm2, and Naïve -Ccm3, suggesting similar existence of low LPS levels in the serum of all mouse strains. **C**. The serum levels of inflammatory cytokine, TNF-α, is low and not influenced by either steroid actions or genotypes. Nearly equal amounts of TNF-α in the serum of all mouse strains were also observed in 30, 60, and 90-day treatment groups with PRG+MIF, suggesting that TNF-α is not influenced by either steroid actions or genotypes. I). No obvious differences for TNF-α were detected among Naïve-WT, Vehicle-WT, Naïve-Ccm1, Naïve-Ccm2, Vehicle-Ccm2, and Naïve-Ccm3, suggesting similar existence of low TNF-α in the serum of all mouse strains. II). Nearly equal amounts of low TNF-α in the serum of all mouse strains were also observed in 30, 60, and 90-day treatment groups of combined steroid (PRG+MIF) treatment, suggesting that low amounts of MCP-1 is not caused by either combined steroid (PRG+MIF) actions or genotypes. A possible immune suppression was observed in Ccm1 at 30-day treatment group. **D.** The serum level of inflammatory cytokine, MCP-1, is low and not caused by either combined steroid (PRG+MIF) actions or genotypes. The concentrations of the cytokine, MCP-1, were measured and quantified by ELISA in naïve (untreated) and Vehicle [vehicle (peanut oil)-treated] conditions. No obvious differences for MCP-1 were detected among Naïve-WT, Vehicle-WT, Naïve-Ccm1, Naïve-Ccm2, Vehicle-Ccm2, and Naïve-Ccm3, suggesting equal existence of low MCP-1 in the serum of all mouse strains**. E.** The concentrations of the cytokine, IL-12, were measured and quantified by ELISA in naïve (untreated) and vehicle [vehicle (peanut oil)-treated] conditions. No obvious differences for IL-12 were detected among Naïve-WT, Vehicle-WT, Naïve-Ccm1, Naïve-Ccm2, Vehicle-Ccm2, and Naïve-Ccm3, suggesting equal existence of low IL-12 in the serum of all mouse strains. **F.** The concentrations of the cytokine, IL-6, were measured and quantified by ELISA in naïve (untreated) and vehicle [vehicle (peanut oil)-treated] conditions. There is decreased IL-6 in Naïve-Ccm2 mice compared to Naïve-WT. No other obvious differences for IL-6 were detected among Naïve-WT, Vehicle-WT, Naïve-Ccm1, Vehicle-Ccm2, and Naïve-Ccm3, suggesting mostly similar existence of low IL-6 in the serum of most mouse strains.

**Fig. S7.**
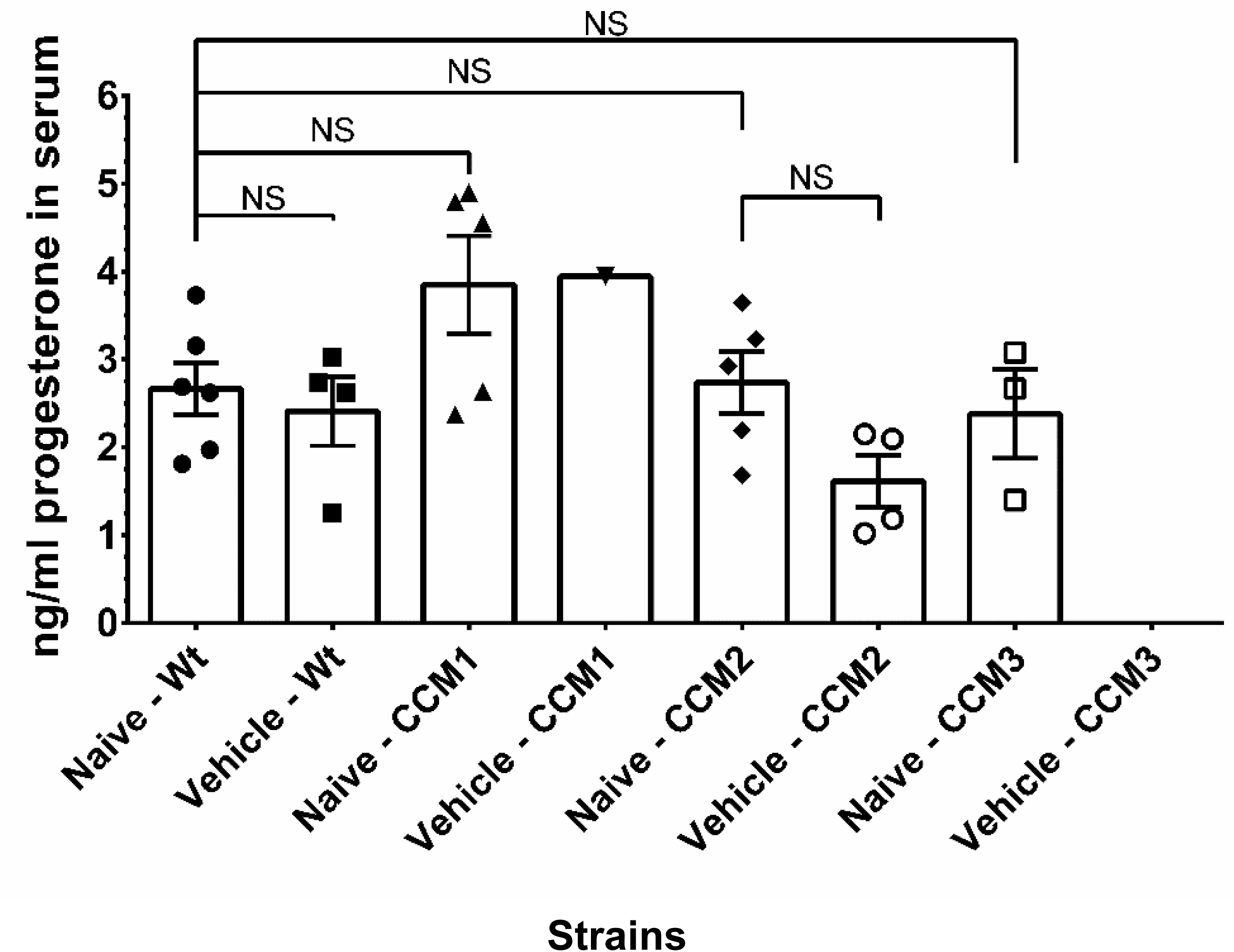
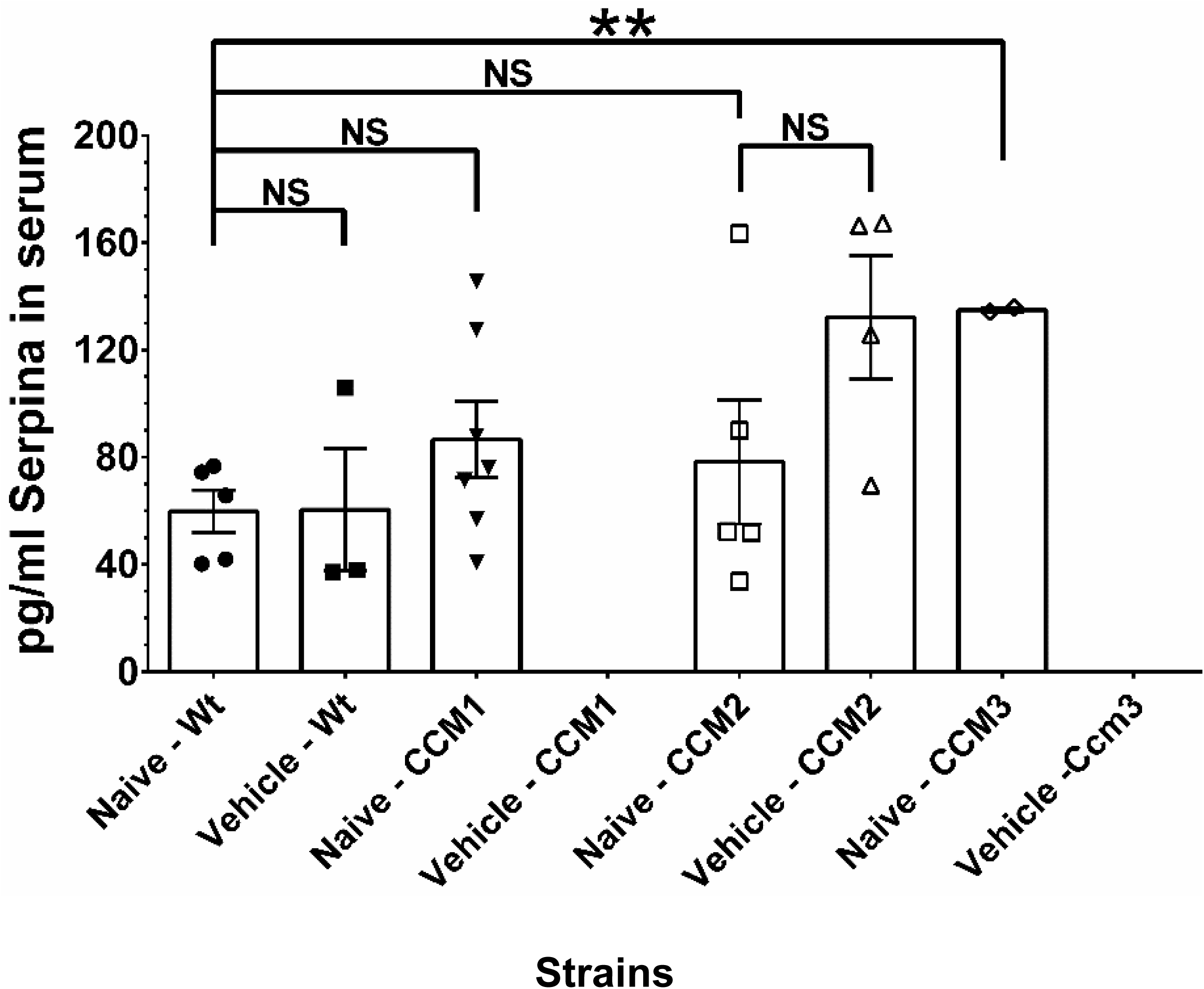
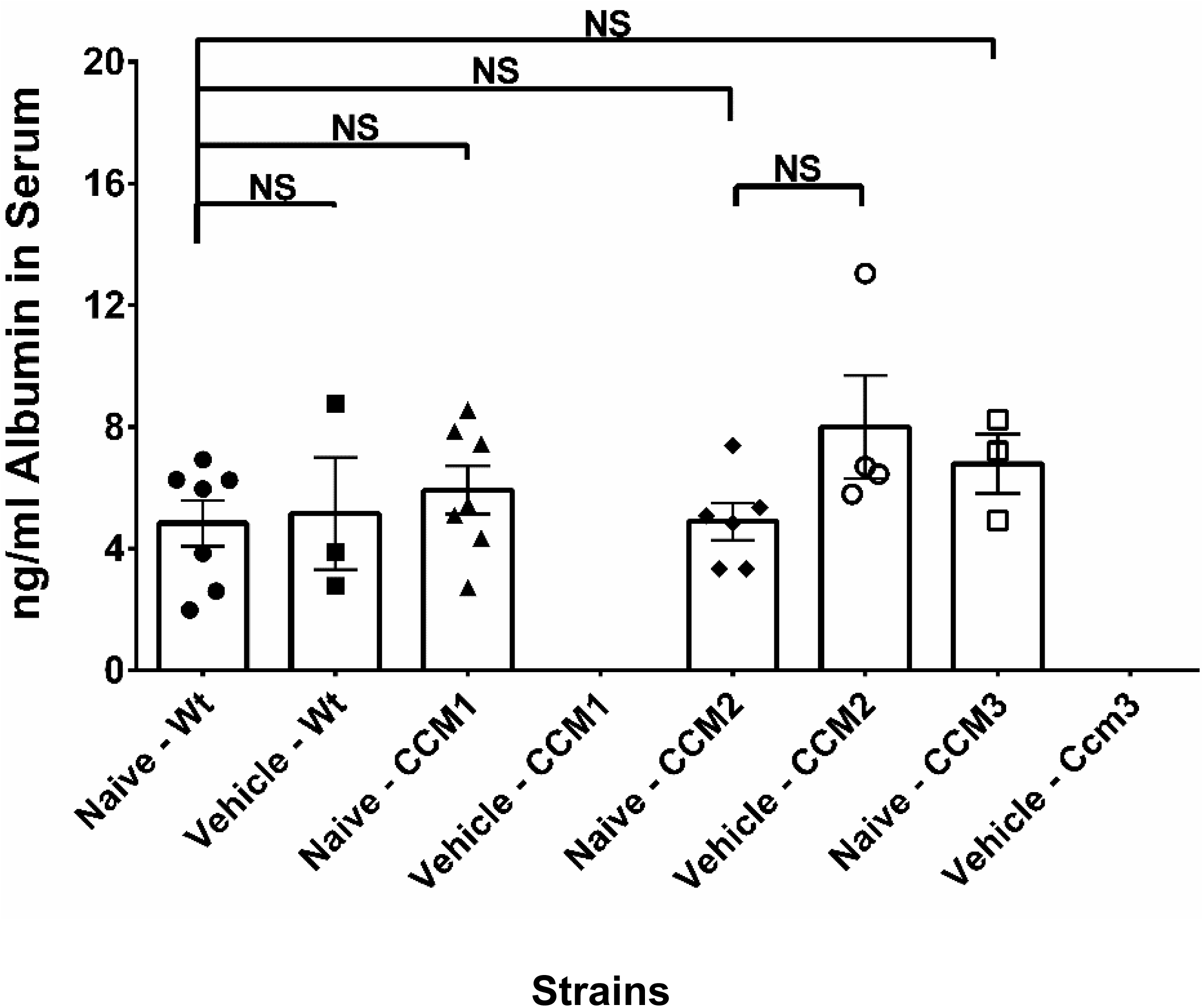
**Perturbation of homeostasis/biogenesis of PRG associated with initial hemorrhagic events in the pathogenesis of CCMs. A.** The concentrations of progesterone (PRG) in mouse serum were measured and quantified by ELISA in naïve (untreated) and vehicle [vehicle (peanut oil)-treated] conditions. No quantitative difference of PRG in mouse serum was detected among naïve (untreated) and vehicle [vehicle (peanut oil)-treated] groups, suggesting equal existence of low PRG in the serum of all control mouse strains. **B.** The concentrations of Serpin A6 in mouse serum were measured and quantified by ELISA in naïve (untreated) and vehicle [vehicle (peanut oil)-treated] conditions. Mostly no quantitative difference of Serpin A6 in mouse serum was detected among naïve (untreated) and vehicle [vehicle (peanut oil)-treated] groups, suggesting mostly similar existence of Serpin A6 in the serum of all mouse strains. Naïve-Ccm3 showed increased levels of Serpin A6 compared to Naïve-WT, possible due to low sample size of naïve-Ccm3 (n=2) **C.** The concentrations of albumin in mouse serum were measured and quantified by ELISA in naïve (untreated) and vehicle [vehicle (peanut oil)-treated] conditions. No quantitative difference of albumin in mouse serum was detected among naïve (untreated) and vehicle [vehicle (peanut oil)-treated] groups, suggesting equal existence of albumin in the serum of all mouse strains.

**Table S1.** **Antibodies used in this study**. Detailed information about application, antigen, clone code, secondary antibody, manufacturer, and category number are listed. * IHC, Immunohistochemistry; IF, immunofluorescence; DAB,HRP/DAB staining, WB, Western Blots, FC, Flow Cytometry; Peritoneal Lavage, PL.

| Application | Antigen |  | Clone | Secondary | Vendor | Cat # |
| --- | --- | --- | --- | --- | --- | --- |
| WB | β-actin |  | C-2 | (anti-mouse) | Santa Cruz | sc-8432 |
| WB | α-actinin |  | H-2 | (anti-mouse) | Santa Cruz | sc-17829 |
| WB | α-actinin-AF488 |  | H-2 |  | Santa Cruz | sc-17829 |
| WB | GAPDH |  | 411 | (anti-mouse) | Santa Cruz | sc-47724 |
| WB | c-Myc |  | 9E/10 | (anti-mouse) | Santa Cruz | sc-40 |
| IF/WB | c-Myc-FITC |  | 9E/10 | (anti-mouse) | Santa Cruz | sc-40 |
| IHC/IF | PECAM-1 -AF488 |  | MEC 13.3 |  | Santa Cruz | sc-18916 |
| IHC/IF | PECAM-1 -AF594 |  | MEC 13.3 |  | Santa Cruz | sc-18916 |
| IHC/IF | Ep-CAM |  | C-10 |  | Santa Cruz | sc-25308 |
| IHC/IF | α-Actin |  | 1A4 |  | Santa Cruz | sc-32251 |
| IHC/IF | α-Actin |  |  |  | ThermoFisher | MA1-06110 |
| IHC/IF | Desmin |  | RD301 |  | Santa Cruz | sc-23879 |
| IHC/IF | Desmin |  |  |  | ThermoFisher | PA5-16705 |
| FC/PL | CD16/CD32 | Fc Shield | 2.4G2 |  | Tonbo Biosciences | 70-0161-U500 |
| FC/PL | CD45.2 -APC |  | 104 |  | eBiosciences | 17-0454-81 |
| FC/PL | CD11b-PerCP |  | M1/70 |  | Biolegend | 101229 |
| FC/PL | MHC class II (I-A/I-E)-FITC |  | M5/114.15.2 |  | Invitrogen | 11-5321-82 |
| FC/PL | Ly6G-PE |  | 1A8 |  | BD Biosciences; | 561104 |
| FC/PL | Ly6C-PB |  | HK1.4 |  | Biolegend | 128013 |
| FC/PL | F4/80 PE-Cy7 |  | BM8 |  | eBiosciences | 25-4801-82 |

**Table S2.** **ELISA kits used in this study.** Detailed information about application, antigen, manufacturers, and category number are listed.

| Gene | Cytokine elisa Kits | Supplier | Catalog # |
| --- | --- | --- | --- |
| IL6 | BD OptEIA Mouse IL-6 ELISA Set | BD biosciences | 555240 |
| IL12 | BD OptEIA Mouse IL-12 (p40) ELISA Set | BD biosciences | 555165 |
| MCP-1 | BD OptEIA Mouse MCP-1 ELISA Set | BD biosciences | 555260 |
| TNF-a | BD OptEIA Mouse TNF (mono/mono) ELISA Set | BD biosciences | 555268 |
| Progesterone (PRG) | Progesterone ELISA | DRG international | EIA-1561 |
| Lipopolysaccharides (LPS) | Mouse Lipopolysaccharides (LPS) ELISA Kit | MyBioSource | MBS7700668 |
| Serum Albumin | Mouse Serum albumin ELISA Kit | EIAAB Science | E1028m |
| SERPINA6 (CBG) | Mouse Corticosteroid-binding globulin (SERPINA6) ELISA Kit | abbkine | KTE70454 |

**Table S3. RNAseq analysis across three endothelial cell lines under steroid actions. A: RAW data for RNAseq analysis for HBMVEC, RBMVEC and HDMVEC:** Based on the gene expression level, we can identify the DEGs (Differentially expressed genes) between samples or groups. We used Possion Distribution algorithms to detect the DEGs. The columns in the table represent the following information: GeneID: Gene ID, Length: Gene Length, Group1-Expression: Gene expression in group1, Group2-Expression: Gene expression in group2, log2FoldChange (group2/group1): The log2 value of ratio of Group1-Expression to Group2-Expression, FDR: Adjusted p-value, Up/Down-Regulation: Up/down-regulated, Pvalue: p-value, Symbol: Gene Symbol ID (Gene Name). **B: Differentially Expressed Genes (DEGs) (RNAseq) identified between Steroid treatment among HBMVEC, RBMVEC and HDMVEC.** Unique and overlapped genes displayed in Venn diagrams in figure 3Bi-iii (Top panels) are provided with the GeneID. There is a separate tab for each section of the V.enn diagrams (3 tabs for each Venn, 9 tabs total. **C: Shared functional enrichment pathways identified between Steroid treatments (Transcriptome) for HBMVECs, RBMVEC and HDMVEC.** Overlapped Biological Processes, Molecular Functions and KEGG pathways displayed in Figure 3D are provided with the Direction, adjusted p-value, number of genes (all significant), pathway description as well as the genes involved (gene symbols) in the overlapped pathways that were differentially expressed in our samples.

**Table S4: Multi-omics analysis in HBMVECs with a disrupted CSC under steroid actions. A: Unique/shared Differentially Expressed Proteins (DEPs) between Steroid treatment (proteome) and Disrupted CSC (individual Ccm KDs, Proteome) for HBMVECs.** DEPs displayed in Venn diagrams in figure 4A (i-iii) are provided with the accession number, fold change by category, and T-Test for each comparing group. Tabs are divided into UP, DOWN, and REST (opposite expression) for each comparison that was made as well as a RAW data tab that contains all the accession number, fold change by category, and T-Test for all 4 groups (Ccm1-KD, Ccm2-KD, Ccm3-KD and MIF+PRG treated). Proteins with significance in 1 of the comparing groups are color coded red and proteins that are significant among both comparing groups have been highlighted yellow. **B: Unique/shared significant genes identified between Steroid treatment (transcriptome and proteome) and Disrupted CSC (Pooled, Proteome) for HBMVECs.** Unique and overlapped genes displayed in Venn diagrams in figure 4B-C (Top and bottom left panels) are provided with the alternate ID (protein), GeneID and symbol (RNA) and Fold Change (most protein data and overlapped RNAseq data) (Unique genes in RNAseq and protein hormone groups do not have fold change, since they are unique to only one treatment group). Unique proteins (black color) found in CSC-KD (pooled CCM1-KD, CCM2-KD, CCM3-KD), Hormone treated samples (compared to vehicle) and the shared genes for each group (red colored) are displayed. **C: Shared functional enrichment pathways identified between Steroid treatment (Transcriptome and Proteome) and Disrupted CSC (Pooled CCM1-KD, CCM2-KD, CCM3-KD, Proteome) compared to Steroid treatments (Transcriptome) for HBMVECs.** Overlapped Biological Processes, Molecular Functions and KEGG pathways displayed in Figure 4B-C (Top and bottom right panels) are provided with the Direction, adjusted p-value, number of genes (all significant), pathway description as well as the genes involved (gene symbols) in the overlapped pathways that were differentially expressed in our samples.

**Video S1. Stroke/loss of motor function phenotype.** Evidence of stroke and loss of motor function in Ccm2 mutant mouse in the 90-day treatment group. Daily evaluations for early hemorrhagic events was performed before and after injections to assess any signs of early hemorrhagic events, resulting from combined steroid actions. We did not observe any stroke and loss of motor function in WT PRG+MIF/vehicle treated mice or WT naïve mice in any of the treatment groups.

**Video S2. Seizure phenotype in Ccm2 mutant, 90-day treatment group.** Evidence of seizure in Ccm2 mutant mouse in the 90-day treatment group. Daily evaluations for early hemorrhagic events was performed before and after injections to assess any signs of early hemorrhagic events, resulting from combined steroid actions. We did not observe any seizure phenotype in WT PRG+MIF/vehicle treated mice or WT naïve mice in any of the treatment groups.

**Video S3. Seizure phenotype in all three Ccms mutants, 90-day treatment group.** Evidence of seizure in Ccm1, Ccm2 and Ccm3 mutant mice in the 90-day treatment group. Daily evaluations for early hemorrhagic events was performed before and after injections to assess any signs of early hemorrhagic events, resulting from combined steroid actions. We did not observe any seizure phenotype in WT PRG+MIF/vehicle treated mice or WT naïve mice in any of the treatment groups.

**Video S4. Seizure phenotype in Ccm2/3 mutants, 90-day treatment group.** Evidence of seizure in Ccm1, Ccm2 and Ccm3 mutant mice in the 90-day treatment group. Daily evaluations for early hemorrhagic events was performed before and after injections to assess any signs of early hemorrhagic events, resulting from combined steroid actions. We did not observe any seizure phenotype in WT PRG+MIF/vehicle treated mice or WT naïve mice in any of the treatment groups.

Table S3 and S4 contain a large amount of raw data for Omic, analytical and bioinformatics analysis for Fig. 3 and Fig. 4. They were submitted in the compressed forms.

